# Inferring context-dependent computations through linear approximations of prefrontal cortex dynamics

**DOI:** 10.1101/2023.02.06.527389

**Authors:** Joana Soldado-Magraner, Valerio Mante, Maneesh Sahani

**Author notes:** **Correspondence:** Dr. Joana Soldado Magraner Neuroscience Institute Carnegie Mellon University Pittsburgh, PA, USA. Denotes shared senior authorship (Valerio Mante, Maneesh Sahani).

## Abstract

The complex neural population activity of prefrontal cortex (PFC) is a hallmark of cognitive processes. How these rich dynamics emerge and support neural computations is largely unknown. Here, we infer mechanisms underlying the context-dependent selection and integration of sensory inputs by fitting dynamical models to PFC population responses of behaving monkeys. A class of models implementing linear dynamics driven by external inputs accurately captured the PFC responses within each context, achieving performance comparable to models without linear constraints. Two distinct mechanisms of input selection and integration were equally consistent with the data. One implemented context-dependent recurrent dynamics, as previously proposed, and relied on transient input amplification. The other relied on the subtle contextual modulation of the inputs, providing quantitative constraints on the attentional effects in sensory areas required to explain flexible PFC responses and behavior. Both mechanisms consistently revealed properties of inputs and recurrent dynamics missing in more simplified, incomplete descriptions of PFC responses. By revealing mechanisms consistent with rich cortical dynamics, our modeling approach provides a principled and general framework to link neural population activity and computation.

## Introduction

A fascinating aspect of our daily existence is that, in a blink of an eye, we can effortlessly change our course of action, switch between tasks or wander in between lines of thought. To achieve this flexibility, brain circuits must be endowed with mechanisms to perform context-dependent computations, so that behavior is quickly adapted to each situation and the correct decisions can be taken. The mechanisms underlying this flexibility are still poorly understood.

A brain structure known to mediate flexible computations is the Prefrontal Cortex (PFC)^1^. PFC is part of an extensive and highly distributed network of cortical and subcortical areas comprising the decision-making circuitry of the brain^2^. It is involved in complex cognitive functions such as planning, selective attention and executive control^3,4^. PFC is thought to hold the representation of goals, contexts and task rules^5,6^ and in primates is required to switch behaviors according to different task instructions^7^. Finally, PFC’s crucial role in ignoring task distractors suggests that it actively filters out irrelevant information^8,9^. This makes PFC of special importance for studying contextual decision-making.

Previous work suggested that flexible prefrontal computations emerge from the concerted interaction of large, interacting neural populations^1^. Surprisingly, during contextual decisions requiring monkeys to integrate noisy sensory information towards a choice, irrelevant information did not appear to be gated at the level of inputs into PFC. Instead, irrelevant inputs may be dynamically discarded through recurrent computations occurring within PFC. A possible mechanism for such dynamical gating was revealed by reverse-engineering recurrent neural networks (RNNs) trained to solve the same contextual decision-making task as the monkeys. Remarkably, the trained RNNs reproduced key features of the PFC population activity, even though the networks were not explicitly designed to match the dynamics of the data. The match with the recorded data, however, was only qualitative, as these networks do not reproduce all aspects of the rich and heterogeneous responses of individual PFC neurons. It is not known whether a model explicitly designed to capture the complex PFC dynamics in its entirety would rely on the same contextual decision-making mechanism as the RNNs.

In this study, we took the approach of fitting linear dynamical system models (LDS) directly to the PFC data, allowing us to infer interpretable low-dimensional (low-d) linear systems that approximate the neural population activity in each context. We characterized the nature of computations implemented in each context by analysing the properties of the fitted models, whose dynamics closely matched those of the PFC population. To validate our assumption of linear dynamics, we compared the LDS to a novel low-rank factorization of the data, Tensor Factor Regression (TFR), which can capture non-linear dynamics. Both models performed comparably, implying that a linear model is sufficient to explain PFC activity in a given context.

We fitted different LDS model classes corresponding to different hypotheses about the nature of context-dependent computations in PFC. One class could implement context-dependent recurrent dynamics but received fixed inputs, mimicking the design of RNNs developed in past work^1^. Another class had fixed recurrent dynamics, but could implement context-dependent inputs. In both models, we inferred external input signals directly from the data. Surprisingly, these two model classes explained the PFC responses similarly well, meaning that both contextual decision-making mechanisms are consistent with the data. Both mechanisms shared some features with the RNN solution, but also differed from it in important ways, revealing previously unknown properties of PFC inputs and recurrent dynamics underlying contextual decision-making computations.

Our data-driven approach to analyzing neural dynamics, based on fitting LDS models to neural population responses, can be applied across different brain areas, neural data sets, and computational mechanisms, providing a general tool to test specific hypotheses about the nature of computations implemented by neural circuits.

## Results

We analysed PFC recordings from two monkeys performing a contextual version of the classic random dots motion task^1,10^. The monkeys had to report the overall color or motion of the dots, depending on context (Fig. 1a). Since both types of sensory evidence were simultaneously presented, the monkeys had to actively ignore the irrelevant sensory input in order to form a decision based only on the relevant input. We analysed only correct trials and focused on the random dots presentation period (750 ms), during which the motion and color evidence needed for a correct decision were presented^1^. In the next sections, we present an in-depth analysis of the PFC data from one of the monkeys (monkey A). Findings from monkey F are presented in the supplementary material, and confirm the key insights gained from monkey A.

**Figure 1.**
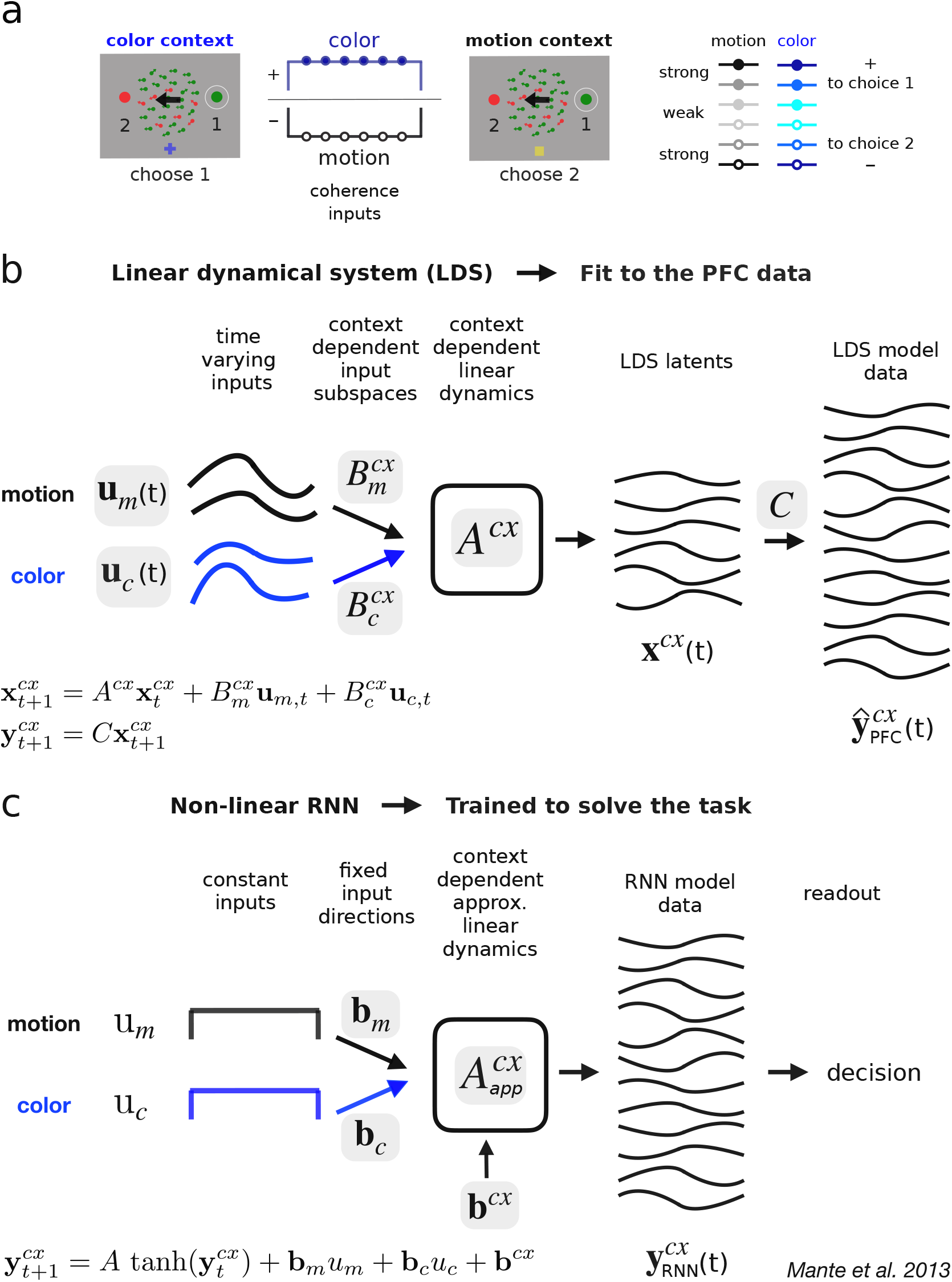
Linear dynamical systems (LDS) model-fitting approach. **a**, Task. The monkeys chose one of two targets based on evidence from either the motion or the color of a random dots display depending on context, indicated by the fixation point (blue cross/yellow square). The motion/color evidence could point towards (choice 1) or away from (choice 2) the target placed on the receptive field of the recorded neurons (white circles). The evidence strength was given by the color/motion coherence level of the random dots, yielding 6 coherence conditions (3 levels, color shades; 2 choices, filled circles/positive values, hollow circles/negative values). Here, strong color and motion evidence (dark shades) point at opposite targets (positive vs. negative). In the color context, the monkey should choose the target matching the overall color of the dots (green; right target, choose 1), but in the motion context, the target indicated by the overall motion direction of the dots (left arrow; left target, choose 2). **b**, An LDS model is fit to the PFC data from both contexts learning either joint or context-dependent linear dynamics *A*^*cx*^ and input subspaces 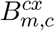. External inputs **u**_*m,c*_(*t*) are also learned (one for each coherence condition in **a**), are fixed across contexts and can vary in time. These parameters determine the evolution of a low-d latent process **x**(*t*) that approximates the dynamics of the high-d PFC data ŷ_*P FC*_(*t*) (via the orthonormal mapping *C*). **c**, Non-linear RNNs were trained by Mante et al. on the same task as the monkeys. External inputs were hand-crafted noisy signals with mean *u*_*m,c*_ constant over time and proportional to the coherence level. Input dimensions were 1d and fixed across contexts **b**_*m,c*_. A context-dependent input vector **b**^*cx*^ switched the dynamics of the fixed non-linear recurrent network between two approximately linear regimes 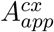. A linear readout pooled network responses to generate a decision signal for training. Network population responses **y**_*RNN*_ (*t*) were only qualitatively compared to the PFC responses. Grey shadings: learned parameters due to training or data fitting.

### Two classes of models implementing context-dependent computation

To infer possible mechanisms underlying PFC population dynamics, we fitted several LDS models to the measured responses (Fig. 1b). Each LDS had three components: a dynamics matrix *A*, which determined the recurrent contribution to a low-d “latent” activity state **x**(*t*); external motion and color inputs **u**_*m*_(*t*) and **u**_*c*_(*t*); and motion and color input subspaces *B*_*m*_ and *B*_*c*_, specifying dimensions along which the external inputs modulated the latent state. The dynamics matrix and the input subspaces were fixed over time, whereas the external inputs could be time-varying. The condition-averaged z-scored peri-stimulus-time-histograms (PSTHs) of individual units in PFC were then reconstructed as a linear combination of the low-d latent dynamics (matrix *C* in Fig. 1b).

Critically, we fitted any given LDS model jointly to the PFC data from the two contexts, whereby only some of the model parameters varied across contexts. We considered primarily two model classes. In what we refer to as the *A*^*cx*^, *B* models, the dynamics matrix *A*^*cx*^ could differ across contexts (Fig. 1b, *cx* = mot/col context), whereas the input parameters were fixed. In the *A, B*^*cx*^ models, on the other hand, the dynamics matrix was fixed, but the motion and color subspaces 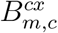 were allowed to vary across contexts (in both direction and norm). These two classes effectively amount to two distinct mechanisms for processing inputs flexibly across contexts.

The *A*^*cx*^, *B* models retain key properties of previously proposed RNNs^1^. As in the RNNs, the motion and color inputs are fixed across contexts, meaning that context-dependent computations must be achieved by the recurrent dynamics (Fig. 1c). The *A, B*^*cx*^ models instead rely on contextually modulated inputs, a mechanism that appeared unlikely based on past analyses of the PFC responses^1^. Both model classes differ from the RNNs in several ways. First, whereas the RNNs were trained on the task, with hand-crafted external inputs that were constant over time, all LDS parameters were fitted to the data (Fig. 1b, grey boxes), including the time-varying inputs **u**_*m*_(*t*) and **u**_*c*_(*t*). Second, whereas the RNNs received one-dimensional inputs, the LDS could learn multi-dimensional input subspaces *B*_*m,c*_ and could thus produce rich activity patterns directly through the inputs^11,12^. To avoid solutions that relied entirely on input driven activity, we fitted the LDS with a regularization favoring weak inputs (Methods). Activity patterns that do not directly represent the motion and color coherence, such as the integrated relevant evidence or activity related to the passage of time, would then have to emerge through the transformation of the inputs by the recurrent dynamics in all LDS models.

We found that the two LDS model classes could explain the PFC responses similarly well (Fig. 2a, *A*^*cx*^, *B* and *A, B*^*cx*^; cold color lines), implying that two very different mechanisms of context-dependent computation are consistent with the observed activity. A third model class that had contextual flexibility in both the recurrent dynamics and the inputs (referred to as *A*^*cx*^, *B*^*cx*^) did not improve the fits. A model that could change only the initial conditions across contexts (Methods), but not the recurrent dynamics or the inputs (referred to as *A, B*) instead performed significantly worse. We estimated the dimensionality of the latent dynamics and inputs based on generalization performance using leave-one-condition-out cross-validation (LOOCV^13^, Fig. 2b). All models performed substantially better for input dimensionality higher than 1d (Fig. 2a), meaning that they required multi-dimensional input signals. The best performing LDS models required 3 dimensions for both the color and motion inputs. The LDS models needed between 13 and 18 latent dimensions to best fit the data (Supplementary Table 1), many more than the 4 dimensions required to describe the task (motion, color, context, and decision).

**Figure 2.**
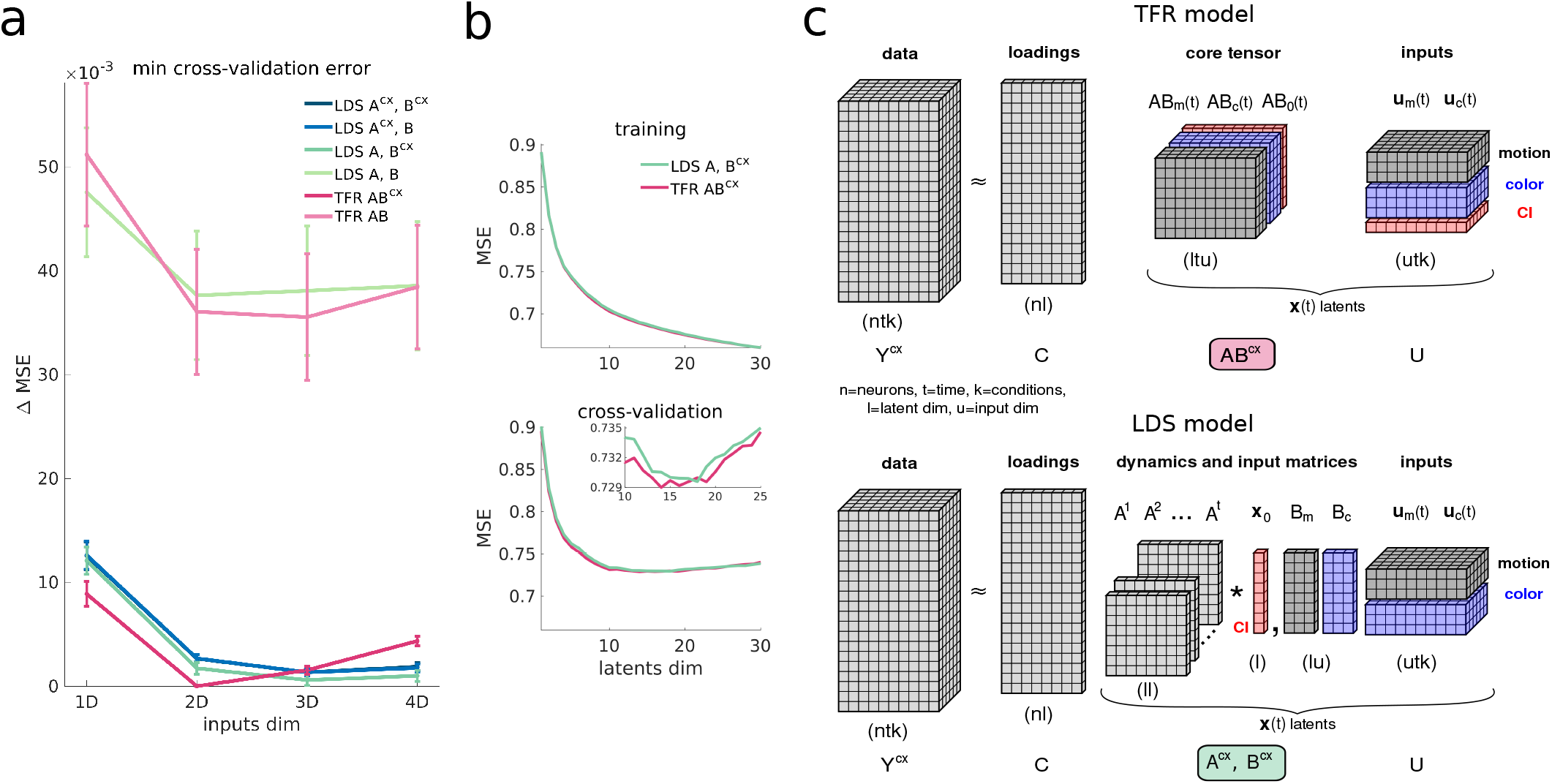
Several LDS model classes capture the PFC data equally well, and comparably to a more flexible Tensor Factor Regression (TFR) model. **a**, Leave-one-condition-out cross-validation performance (LOOCV)^13^ for LDS and TFR models with different input dimensionalies and contextual constraints. Shown are minimum cross-validation mean squared errors (MSE) across all latent dimensionalities, relative to the best performing model (*A, B*^*cx*^ model with 3d inputs and 18 latents). The latent dimensionality for which the minimum was attained is in Supplementary Table 1. Error bars, standard error mean (sem) across LOOCV folds (36 task conditions, Methods). *A*^*cx*^, *B*^*cx*^ line is below *A*^*cx*^, *B*’s. **b**, Training and LOOCV performance for the best LDS and TFR models across different latent dimensionalities (TFR 2d *AB*^*cx*^ and LDS 3d *A, B*^*cx*^; min LOOCV latent dim = 14 and 18). Monkey A data. **c**, TFR model (top). The data tensor *Y* is factorized into 3 low-rank tensors, all learned. The loadings *C* (an orthonormal matrix) sets the rank of the factorization and maps the low-d core tensor *AB* into the high-d neural space. The low-d latents **x**(*t*) are generated by multiplying the core tensor and the input tensor *U*, which captures motion, color and condition independent (CI) signals. For clarity, two indicator tensors are omitted, one recreating an LDS-like temporal convolution of the core tensor and inputs, and another one used to repeat the inputs across the 36 task conditions (Methods). To generate context-dependent activity *Y* ^*cx*^, the core tensor can change across contexts *AB*^*cx*^. The LDS model (bottom) is nested within the TFR model class (Methods). The core tensor *AB* is replaced by a smaller set of parameters, *A* and *B*. The data temporal structure is restricted to be captured by powers of *A* over time (Methods). The latents are recreated by convolving (asterisk symbol) the dynamics matrix powers with the inputs and adding the initial conditions *x*_0_. Inputs are also repeated across task conditions.

### PFC dynamics in each context are well approximated by a linear system

Several observations imply that fits of the *A*^*cx*^, *B* and *A, B*^*cx*^ models provide accurate descriptions of the PFC responses.

First, both models closely approximated the highly heterogeneous responses of individual PFC neurons (Extended Data Fig. 1). The fits captured a substantial fraction of the variance in the data (27%, corresponding to MSE=0.73 on z-scored responses, Fig. 2b), which included poorly fit neurons with weak or sparse responses (Extended Data Fig. 1, firing rates *<* 1Hz), and were not smoothed or “de-noised”^1^.

Second, the best LDS models performed comparably to a more powerful model class that we refer to as Tensor Factor Regression (TFR) (Fig. 2a,b, warm color lines). TFR is based on a new low-rank factorization of the data that partitions the data tensor into several low-d tensors, including a core tensor and an input tensor (Fig. 2c). The core tensor contains time-varying parameters that provide TFR with greater fitting flexibility compared to the LDS models, which are constrained to generate linear dynamics. The LDS models are nested within the TFR class (Methods), simplifying model comparisons^14^. TFR incorporated input parameters and contextual constraints equivalent to those of the LDS (Fig. 2c, Methods). The fitted latent dimensionality was similar in the TFR and LDS fits (13-18d, Supplementary Table 2), but the optimal input dimensionality was lower for TFR compared to the LDS fits (2d vs 3d; Fig. 2a). The extra input dimension in the LDS fits could imply limitations of the linear dynamical constraints. Beyond this difference, however, the greater flexibility in TFR provided little or no advantage to the fits (Fig. 2a,b), confirming that linear dynamics provide an accurate description of the data.

Third, the best LDS models qualitatively captured salient features of the population dynamics equally well. In particular, both the *A, B*^*cx*^ and the *A*^*cx*^, *B* models reproduced PFC population trajectories in the activity subspace capturing most variance due to motion, color and choice across contexts^1^ (Fig. 3). TFR fits were comparable, both at the population (Extended Data Fig. 2a) and single neuron level (Extended Data Fig. 1). One key implication of this finding is that the properties of population trajectories in the considered low-d activity subspace are not sufficient to distinguish between mechanisms that rely on inputs that are fixed or variable across contexts.

**Figure 3.**
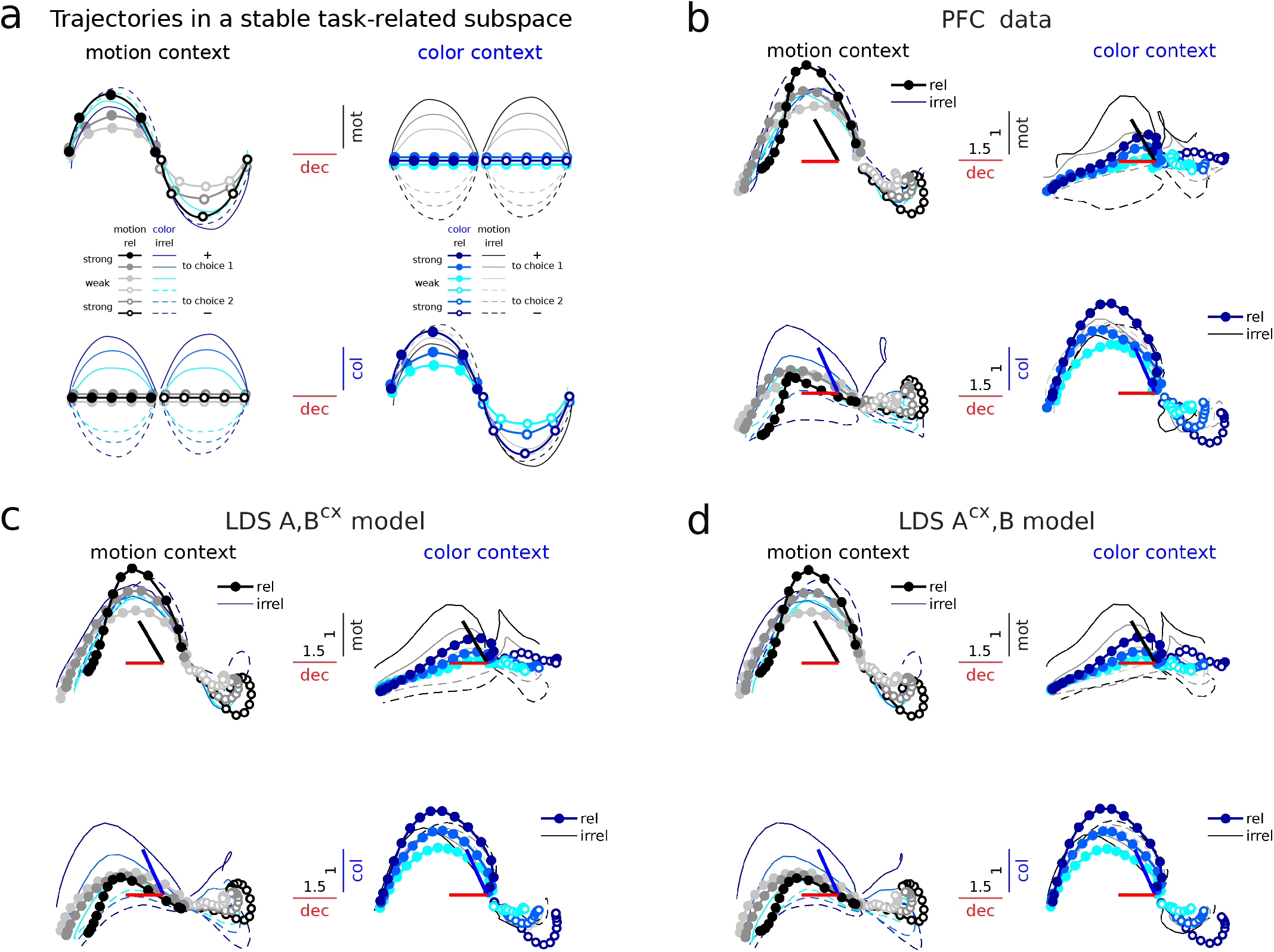
LDS models with both fixed and context-dependent input dimensions capture PFC trajectories in a contextually-stable task-related subspace. **a**, Expected population trajectories in a hypothetical subspace, fixed across contexts, capturing most variance due to motion, color and choice. Trajectories are sorted by choice and coherence conditions (as in Fig. 1a). The same trajectories in each context are sorted twice, by either motion or color, and then plotted along different task-related dimensions (top, motion and choice; bottom, color and choice). Stable input dimensions encode coherence information (sign and strength) of a given input regardless of context, but not of other inputs (top, motion dimension encodes motion coherence, in black, but not color coherence, in blue; bottom, the reverse); i.e., stable input dimensions encode coherence signals regardless of whether these are relevant or irrelevant in a given context (thick doted vs. thin lines). In contrast, a stable decision-related dimension reflects the sign (and the integrated strength) of the relevant input in each context, but not the irrelevant one—a signature of selective integration^1^ (decision axes separate filled vs. hollow circles, the relevant input sign, but not filled vs. dashed lines, the irrelevant; i.e. motion in the motion context, left, and color in the color context, right); Thus, the movement of the trajectories along the choice axis is coupled to the evolution along the relevant input axis, but not the irrelevant. This suggests that choice-related activity emerges from the relevant input signal. Input signals are assumed transient along the input dimensions. **b**, PFC trajectories in the task-related subspace found by Mante et al. using targeted dimensionality reduction (TDR)^1^, for monkey A. The subspace captures motion, color and choice-related variance along a set of orthonormal axes that are stable across contexts. Colored thick bars, angle between TDR axes before orthogonalization. Numbers on bars, scaling factor to ease visualization^1^. Trajectories are sorted by choice and motion/color coherence conditions, with color/motion conditions averaged out^1^. **c** Cross-validated model trajectories (LOOCV) for the best LDS *A, B*^*cx*^ model (3d inputs, 18d latents). **d** Same for the *A*^*cx*^, *B* model (3d inputs, 16d latents). Trajectories have been smoothed with a Gaussian filter for visualization (sliding window size, 5-bins).

To understand the mechanisms of context-dependent integration in the two model classes, below we first separately characterize the inputs and recurrent dynamics in the *A, B*^*cx*^ and the *A*^*cx*^, *B* models, and then ask how their combined effects can account for contextual integration in PFC. We assessed the robustness of the inferred mechanisms by fitting 100 models for each class (random initialization) with the dimensionality of inputs and latent state set by the above cross-validation results (inputs: 3d; latent: 18d and 16d, for *A, B*^*cx*^ and *A*^*cx*^, *B*; Fig. 2a, Supplementary Table 1).

### Input signals span curved manifolds and are largely stable across contexts

In the models, the strength of contextual modulation can be summarized by the norm of the latent activity, which we term the model “output” (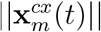 and 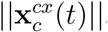, Fig. 4a). We computed the output in each context by setting either the color or motion input to zero. The model output is essentially identical across model classes: it increases throughout the trial and is much larger for the relevant compared to the irrelevant input, reflecting context-dependent integration (Fig. 4b,c, bottom panels, thick vs. thin curves; green bars: p*<*0.001, Wilcoxon rank-sum test, N=100 models). In the *A*^*cx*^, *B* model, this context dependence is due to differences in the recurrent dynamics across contexts, whereas in the *A, B*^*cx*^ model it reflects contextual modulation of the strength and/or the direction of the inputs.

**Figure 4.**
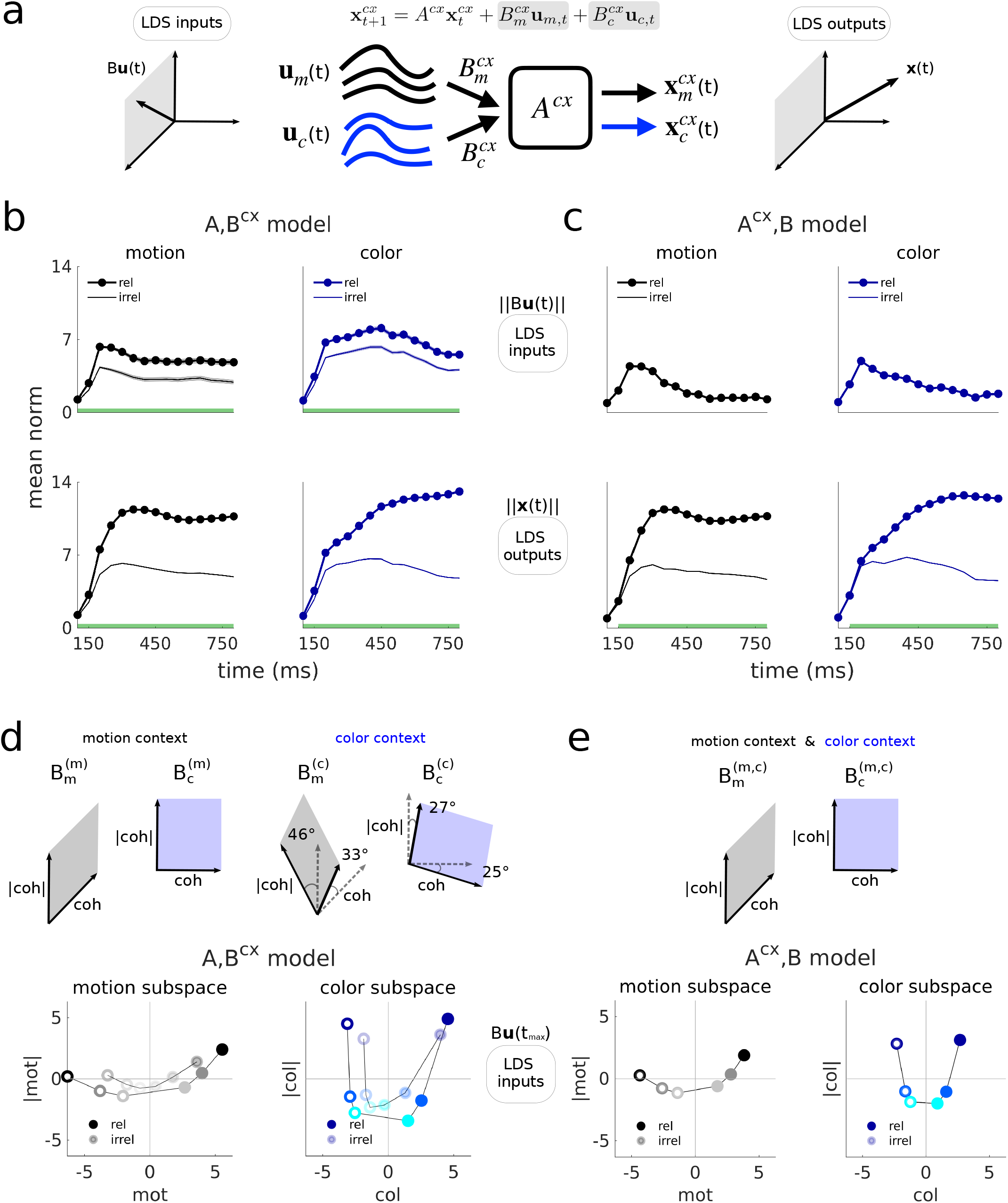
LDS inputs are integrated selectively by both models, are largely stable across contexts and span curved manifolds. **a**, Schematic illustrating the inputs and outputs of the LDS. The strength of the external inputs over time is defined as the norm of the input vectors at each time step ||*Bu*(*t*)||, which accounts for the norm of B. Input vectors lay in the subspace spanned by the columns of *B*, meaning that the input trajectories are confined within the input subspaces (left, here a 2d subspace for illustration). The latents span the full low-d LDS subspace (right, here 3d) and can be seen as the output of the LDS model, since they result from the convolution of the inputs with the dynamics (Methods). **b** Top, external input strength over time inferred for the *A, B*^*cx*^ model across contexts (relevant vs. irrelevant), here shown for the strongest positive coherence. Bottom, same but for the outputs (latents’ norm). Mean across 100 models. Shades = sem (not visible in the outputs). Green bars, times when relevant and irrelevant inputs/outputs are significantly different (Wilcoxon rank-sum test, p*<*0.001). **c** Same but for the *A*^*cx*^, *B* model. Inputs across contexts are the same, by construction. **d**,**e**, Orthonormal 2d subspaces that demix coherence and coherence magnitude variance (coh and |coh|), found within each 3d input subspace in both models, by linearly regressing the inferred external input values against the experimental coherence values and their magnitudes. Shown are mean inputs across 100 models for all coherences, at t=250ms (after input norm pick strength, Fig. 4b,c) and projected onto the 2d coh-|coh| planes. These form a curved representation of coherence information. Lines are drawn to ease visualization. For the *A, B*^*cx*^ model input projections are shown onto the plane bisecting the two input planes found for each context, which were highly aligned (angles between dashed and filled lines). Color and motion planes were nearly orthogonal within each context for both models. For all input values the mean across conditions has been subtracted out to remove condition independent signals (CI). The outputs here are computed running through the dynamics the CI subtracted motion and color inputs independently. Monkey A data.

We computed the strength of the motion and color inputs by pooling contributions from all the corresponding input dimensions (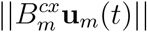 and 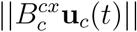; Fig. 4b,c, top panels). Both inputs were somewhat more transient and weaker in the *A*^*cx*^, *B* compared to the *A, B*^*cx*^ model, but were otherwise similar across the two model classes. By definition, input strength was fixed across contexts in the *A*^*cx*^, *B* model. In the *A, B*^*cx*^ model, the irrelevant inputs were weaker than the relevant ones, but only modestly (Fig. 4b, top panels, thick vs. thin curves, green bars: p*<*0.001, Wilcoxon rank-sum test; avg. decrease of 38 ± 14% and 22 ± 8% for mot and col at t*>*200ms, mean ± std, N=100 models).

Even though our cross-validation procedure inferred 3-dimensional input subspaces, most of the inferred input variance was contained in a 2d plane (Extended Data Fig. 3a). Indeed, LDS models with 2d and 3d-inputs performed similarly (Fig. 2a, Extended Data Fig. 2c,d), whereas models with 1d-input performed worse (Fig. 2a, Extended Data Fig. 2e,f). The input plane was spanned by dimensions that separately captured variance related to input coherence (mot, col) and coherence magnitude (|mot|, |col|), implying that inputs were represented along a curved 1d-manifold within the plane^15^. Such curved representations were found in both models (Fig. 4d,e), both contexts of the *A, B*^*cx*^ model (Fig. 4d), and also in the PFC data (Extended Data Fig. 3b, Extended Data Fig. 4).

The input planes were highly aligned between model classes (16-31°, average planes, N=100 models per class; Extended Data Fig. 3c), an effect not expected by chance (Extended Data Fig. 3d). In the *A, B*^*cx*^ model, the motion and color planes varied across contexts, but only modestly (33° ± 10 mot, 46° ± 16 |mot|, 25° ± 5 col, 27° ± 9 |col| dims, mean ± std, N=100 models; Fig. 4d, Extended Data Fig. 3c) and less than expected by chance (Extended Data Fig. 3d). These small changes in input direction across contexts, together with the concurrent, modest change in input strength (Fig. 4b, top), fully account for changes in the output of the *A, B*^*cx*^ model (Fig. 4b, bottom).

In both models, the time-course (Fig. 4b,c, top; Extended Data Fig. 4) and structure of the inputs (Fig. 4d,e) is thus relatively simple. This finding alleviates a possible confound inherent in fitting LDS with time-dependent inputs. In principle, the fitted inputs could be very rich, and effectively approximate on their own the dynamics of a very complex, non-linear dynamical system. As we retrieved inputs of much lower dimensionality than the recurrent dynamics (3d vs. 16-18d), this scenario appears unlikely. Indeed, constraining the inputs to be fixed over time results in only a small drop in performance (Extended Data Fig. 5a,b, Supplementary Note 1). The observed complexity of PFC responses thus need not be inherited from the external inputs, but rather can be explained as resulting from approximately linear recurrent dynamics.

### Input integration relies on high-dimensional linear dynamics

We analysed the recurrent dynamics with an approach originally introduced for the non-linear RNNs trained to solve the context-dependent integration task^1^. Context-dependent computations in the RNNs reflect four key features of the local linear approximations of the dynamics (Fig. 5a, left panel). First, the dynamics has one eigenvalue with norm close to one, and all other eigenvalues smaller than one, implying integration along a line-attractor^16^. Second, inputs are selected for integration by changing the direction of the leading left-eigenvector of the dynamics (the “input-mode” associated with the largest eigenvalue, i.e. slowest dynamics), such that it is orthogonal to the contextually irrelevant input. Third, the direction of the leading right-eigenvector of the dynamics (the “output-mode” associated with the largest eigenvalue), which determines the direction of the line-attractor, is fixed across contexts. Fourth, the leading right and left eigenvectors have different directions, implying “non-normal” dynamics^17,18^. We refer to these four features of the dynamics as the “RNN-mechanism” and below compare them to the dynamics of the fitted models.

**Figure 5.**
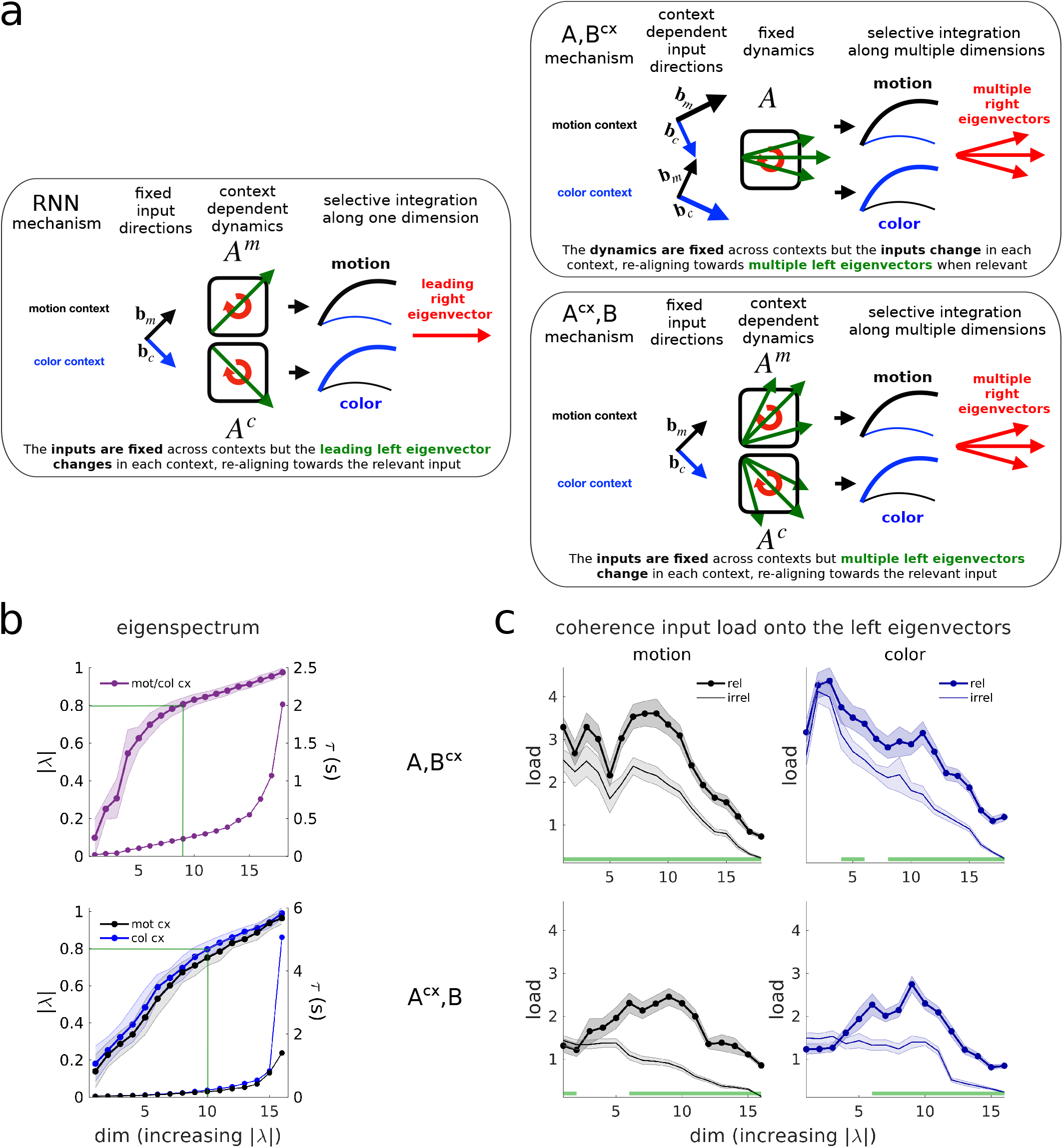
Selective integration requires multiple linear dynamics modes. **a**, The RNN (left) was build with fixed input directions across contexts *b*_*m,c*_. The dynamics in each context switched between two approximately linear regimes, defined by the linearized dynamics matrices *A*^*m*^ and *A*^*c*^. The leading left eigenvector of *A*^*m,c*^ was realigned towards the relevant inputs in each context, loading them onto the slowest output mode of the dynamics (the leading right eigenvector, with associated eigenvalue close to 1), which defined a one-dimensional integrator or line-attractor system^1^. The two LDS models (right) performed a realignment of either the inputs (*A, B*^*cx*^) or the left eigenvectors (*A*^*cx*^, *B*) across contexts, which loaded the inputs onto multiple modes. The *A, B*^*cx*^ model also increased the relevant input’s norm (Fig. 4b, bigger input arrows in cartoon). **b**, Average eigenvalues norm across 100 models (shades=std). The norm sets the rate of decay of each mode (time constant *τ*), and determines the stability of the dynamics (|*λ*| *>* 1 expanding mode, |*λ*| *<* 1 decaying mode, |*λ*| = 1 integration mode). Slow modes have norms close to one (0.8 *<* |*λ*| ≤ 1, *τ >* 224ms, green bars, Methods). The *A*^*cx*^, *B* model learns similar eigenvalues across contexts. **c**, Average coherence input loads onto the eigenmodes of the dynamics across 100 models (shades=sem). The loads were defined by the non-normalized projection of the coherence inputs onto the left eigenvectors, averaged across all time steps (Methods), here shown for the strongest positive coherence inputs. The relevant input loads are significantly higher than the irrelevant loads across multiple eigenmodes (green bars, Wilcoxon rank-sum test, p*<*0.05). Monkey A data.

In agreement with the RNN-mechanisms, both model classes inferred a largest eigenvalue with norm close to 1 (0.98 ± 0.02, *A, B*^*cx*^; 0.96 ± 0.03 / 0.99 ± 0.03, mot/col context, *A*^*cx*^, *B*; mean±std, N=100 models), implying decay time-constants longer than the trial duration (2.5s and 1.2 / 5s). However, both models inferred a surprisingly large number of additional eigenvalues associated with relatively slow decay (Fig. 5b; |*λ*| *>* 0.8, i.e. *τ >* 224ms, for a 750ms trial). Such “slow modes” were most prominent in the *A, B*^*cx*^ compared to the *A*^*cx*^, *B* models (55 ± 7% vs. 35 ± 8% mot cx/ 41 ± 8% col cx; mean±std, N=100 models). The large number of slow modes in both models suggest that PFC dynamics may be higher-dimensional than predicted by the RNN-mechanism.

We assessed context-dependent relations between the recurrent dynamics and the inputs by focusing on the input dimension representing coherence, while ignoring the representation of coherence magnitude (Fig. 4d,e). In the considered linear models, only the coherence component of an input can contribute to choice-dependent responses. To assess the alignment of inputs and recurrent dynamics, we first computed the “load” (the non-normalized projection, Methods) of the coherence component onto each left-eigenvector at each time, and then averaged over times (Fig. 5c). Consistent with the RNN-mechanism, these input loads were overall larger for the relevant vs. the irrelevant input, in both models (Fig. 5c, green bars: p*<*0.05, Wilcoxon rank-sum test, N=100 models). The load along the leading left-eigenvector was close to zero for the irrelevant input, as in the RNN-mechanism. Surprisingly, however, the largest loads overall were consistently obtained for eigenvectors with intermediate eigenvalues (|*λ*| = 0.7 − 0.8, *τ* = 140 − 224ms) and thus relatively fast decay time-constants (Fig. 5b,c). Notably, the prominent differences in input loads across contexts reflect very different mechanisms in the two models: changes in the input strength and direction in the *A, B*^*cx*^ model, and changes in the recurrent dynamics in the *A*^*cx*^, *B* model (Fig. 5a, right panels).

### Non-normal dynamics makes model-specific contributions to selective integration

The qualitative similarity in the eigenvalues (Fig. 5b, Extended Data Fig. 6a) and input loads (Fig. 5c) for the *A, B*^*cx*^ and *A*^*cx*^, *B* models masks a key difference in the recurrent dynamics they implement. Specifically, the two models implement dynamics with very different degrees of non-normality. We assess the strength of non-normality through one of its possible consequences, namely the transient amplification of perturbations of the activity^17,18^ (Supplementary Note 2, Extended Data Fig. 7). We simulated dynamics resulting from a short perturbation or pulse of activity at trial onset, along random state-space directions. For the *A, B*^*cx*^ model, the perturbations gradually decay over the course of the trial (Fig. 6a, top, dashed lines, average across pulses in random directions). For the *A*^*cx*^, *B* model, instead, activity following a perturbation is transiently amplified, i.e. the gradual decay is preceded by a transient increase in activity (Fig. 6a, bottom, dashed lines). For perturbations along the left-eigenvectors, transient amplification is even more pronounced in the *A*^*cx*^, *B* model, but still largely absent in the *A, B*^*cx*^ model (Fig. 6a, doted lines). Dynamics is thus strongly non-normal in the *A*^*cx*^, *B* model, as in the RNN mechanism, but less so in the *A, B*^*cx*^ model (Fig. 6c, Extended Data Fig. 6d).

**Figure 6.**
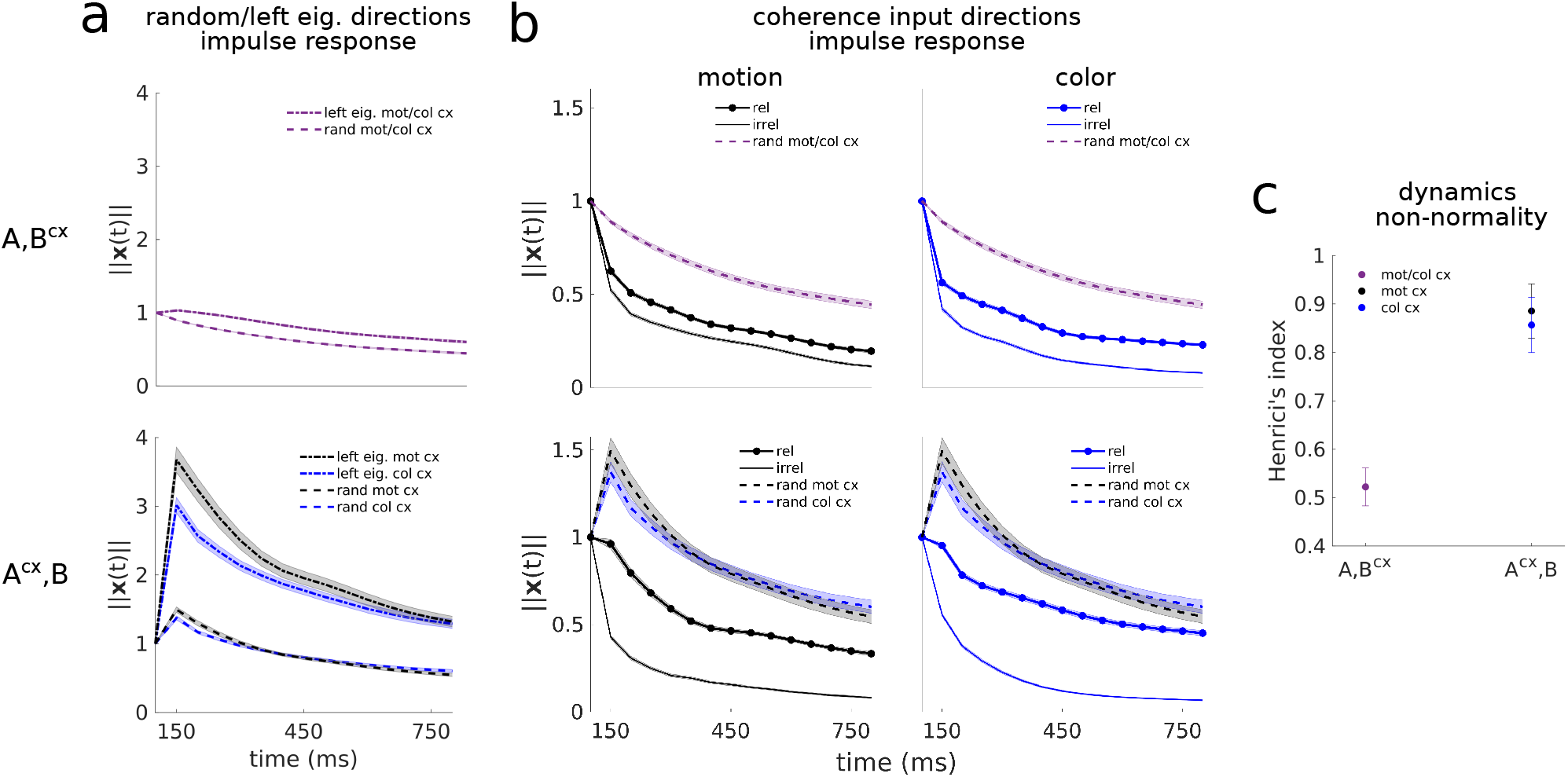
Non-normal transient dynamics contributes to selective integration in the *A*^*cx*^, *B* model. **a**, Models *A*^*cx*^, *B* and *A, B*^*cx*^ mean impulse response for perturbations along random directions (dashed lines) and along the left eigenvectors (dotted lines), averaged across 100 models and across left eigenvectors or perturbations (num pert. = num left eigv. = 16/18). This measure shows how the dynamics matrix transforms a perturbation (or input) of unit norm, by tracking the state norm of the system ||**x**(*t*)|| over time. Note that the *A*^*cx*^, *B* system has a different impulse response for each context, since the dynamics matrix *A* changes in each context. Shades, sem across 100 models. **b**, Impulse response for unit norm perturbations along the motion and color coherence input dimensions. For the *A, B*^*cx*^ model the dynamics matrix is the same across contexts, and thus, the difference in the impulse response between perturbations along the relevant and the irrelevant input dimensions arises due to the fact that these input dimensions subtly change across contexts. For the *A, B*^*cx*^, the perturbations are applied along the same input directions across contexts, since these are fixed, but the dynamics matrix changes, which causes a different transformation of the same input pulse in each context. Note that the impulse response along the input directions is different from the average impulse response along random directions (dashed lines, same as in **a**), which indicates processing selectivity of the dynamics along the input directions. Shades, sem across 100 models. **c**, Degree of non-normality of the two model classes (Henrici’s index, Methods). Error bars, std across 100 models. Monkey A data.

These differences in recurrent dynamics between models are also apparent in their responses to input perturbations (along the coherence dimension, Fig. 6b). In the *A, B*^*cx*^ model, input pulses are not transiently amplified, but rather immediately decay, whether they are relevant or not (Fig. 6b, top, thick and thin curves). In the *A*^*cx*^, *B* model, the relevant input is transiently “persistent”, due to non-normal dynamics (Supplementary Note 2), whereas the irrelevant input quickly decays (bottom). Also at longer time-scales, the decay of a relevant input pulse is faster in the *A, B*^*cx*^ compared to the *A*^*cx*^, *B* model, indicating less accurate input integration. Overall, the recurrent dynamics in the *A, B*^*cx*^ model thus cannot sustain relevant input pulses as well as in the *A*^*cx*^, *B* model (Fig. 6b, top vs. bottom thick curves). This difference explains why the *A, B*^*cx*^ model infers inputs that are stronger and less transient than in the *A*^*cx*^, *B* model (Fig. 4b,c, top).

The features of the dynamics considered so far imply that the two LDS models implemented mechanisms of selection and integration that share key properties of the RNN mechanism. Like the RNNs, all LDS models ultimately relied on a context-dependent realignment between the inputs and a subset of the modes of the recurrent dynamics, either through a change of the inputs (*A, B*^*cx*^) or of the recurrent dynamics (*A*^*cx*^, *B*). And like the RNNs^1^, the *A*^*cx*^, *B* model (but not the *A, B*^*cx*^ model) implemented strongly non-normal recurrent dynamics. However, while the RNNs implement only a few slow modes (and approximate a “line attractor”^1^), both LDS models inferred overall higher-dimensional dynamics, with a comparatively large number of slow modes. As we show below, the functional consequences of these slow modes become apparent when considering how the neural trajectories emerge from the recurrent dynamics.

### Input integration occurs in two distinct phases

Any explanation of how the neural trajectories predicted by the models emerge from the interaction of inputs and recurrent dynamics must include the properties of the right eigenvectors of the dynamics matrix. Whereas the left-eigenvectors determine how inputs are coupled to the recurrent dynamics (Fig. 5c), the right eigenvectors determine “where” in activity space the inputs are mapped onto.

We separately consider condition-dependent (CD) and condition-independent (CI) components of the neural trajectories. CD components were the primary focus of past accounts of this data^1^, and late in the trial primarily capture choice-related activity. CI components, on the other hand, capture prominent structure in the trajectories that is common to all conditions and choices, and appears related to the passage of time in a trial. To identify the modes of the dynamics contributing to CD or CI components, we computed the alignment between the right eigenvectors of the dynamics and the dimensions capturing most CD or CI variance at a given time in the trial (Fig. 7, *A*^*cx*^, *B* model, motion context; Extended Data Fig. 8, all models and contexts). Only right eigenvectors that are well aligned with a given CD or CI dimension can contribute to response variance along that dimension.

**Figure 7.**
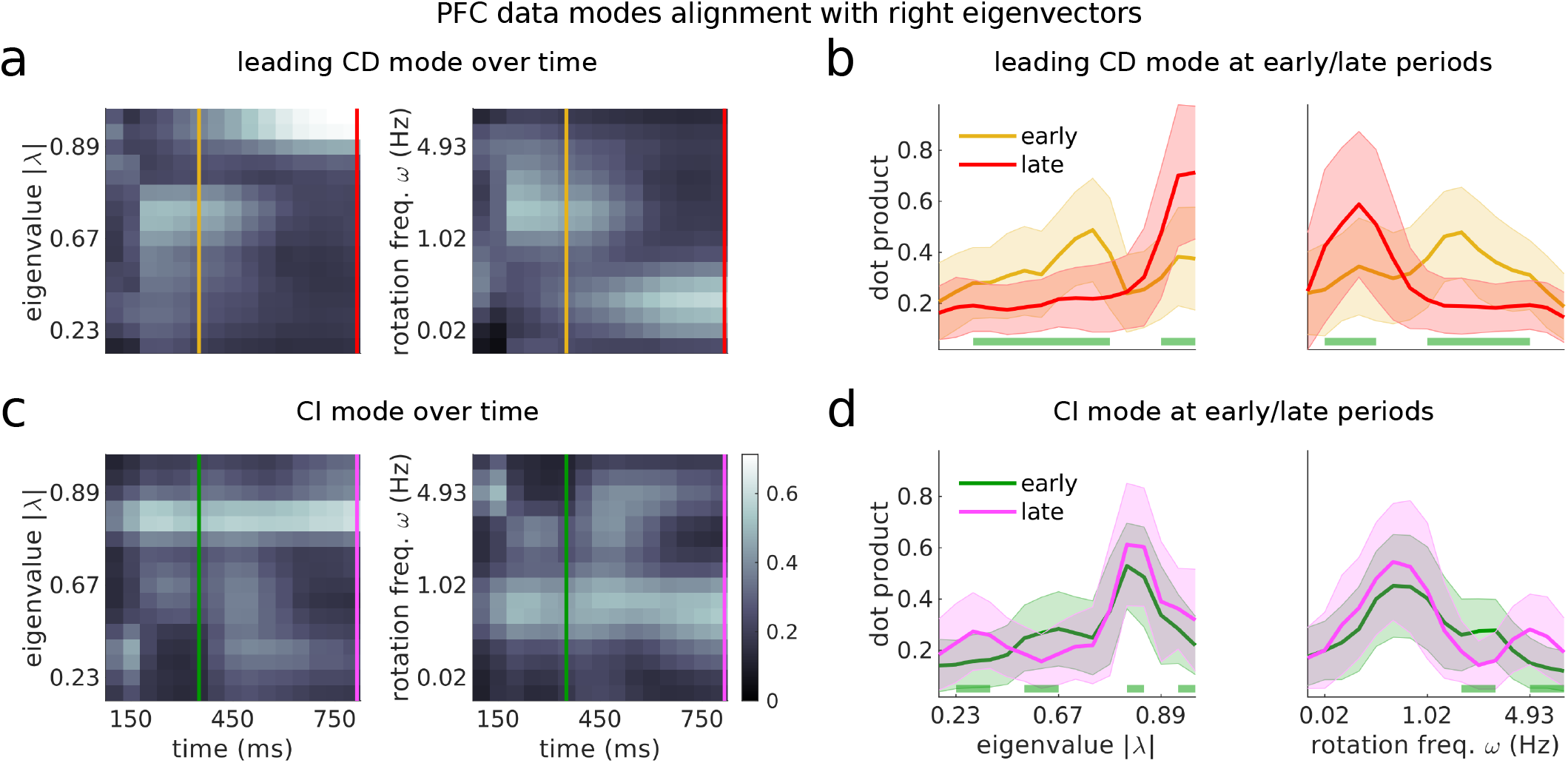
Integration of the relevant inputs occurs in two separate phases of the dynamics. **a**, Largest variance dimension of the condition dependent (CD) data (i.e. the leading singular vector of the data with the across-condition mean subtracted) in the motion context, and at each time step, projected onto the right eigenvectors of the dynamics from the *A*^*cx*^, *B* model (dot products for real eigenvectors, cosines of minimum subspace angles for complex conjugate pairs of eigenvectors, Methods). Left/right panel shows dot products sorted by increasing eigenvalue norm/rotation frequency of the right eigenvectors (averaged across 100 models). Yellow lines mark the early phase of the integration process, (t=350ms, the time at which the integrated motion signal in Fig. 4b,c picks and saturates). Red lines indicate the late phase of the integration process (the last time step of the trial, where decision signals are the strongest^1^). **b**, Mean distribution of alignments across 100 random models at the early and late phases (at times marked in **a**). Shades = std. Green bars indicate the eigenvalues along which the early and late alignment distributions significantly differ (Wilcoxon rank-sum test, p*<*0.001). **c**,**d** Same as panel **a**,**b** but for the CI data vectors (condition-averaged data vectors). Green/purple lines mark the same periods as yellow/red lines. Monkey A data.

The alignment between CD dimensions and right eigenvectors suggests that input integration occurs in two phases characterized by distinct dynamics. Early in the trial, CD responses occur primarily along right-eigenvectors corresponding to modes implementing relatively fast decay and fast rotations (|*λ*| = 0.7 − 0.8, decay time constant *τ* = 140 − 224ms, rotation frequency *f >* 1Hz, Fig. 7a,b, yellow lines). Late in the trial, CD responses instead occur along right-eigenvectors with very slow decay and weak or no rotations (|*λ*| *>* 0.9, *τ >* 475ms, *f <* 0.25Hz, red lines). This transition occurs consistently across model classes, contexts and model initializations (Extended Data Fig. 8). The differences in decay constants and rotational frequencies of the best aligned modes early vs. late in the trial are highly significant (Fig. 7b, Extended Data Fig. 8b, p*<*0.001, Wilcoxon rank-sum test). These observations imply that the relevant input is initially integrated along multiple decaying and rotational modes. Indeed, the relevant inputs load most strongly onto left eigenvectors with intermediate eigenvalues (Fig. 5c). Later in the trial, the relevant input is further integrated and maintained along at set of different, more persistent and non-rotational modes.

The CI components in the responses are mediated by modes that differ from those mediating the CD components, and that appear largely fixed throughout the trial (Fig. 7c,d, Extended Data Fig. 8c,d). At all considered times, the CI components are best aligned with a fixed set of modes that decay more slowly than the early CD-aligned modes, but more quickly than the late CD-aligned modes (|*λ*| = 0.8 − 0.9, *τ* = 224 − 475ms) and are associated with rotational frequencies that are smaller than those in early CD-aligned modes, but faster than late CD-aligned modes (*f* = 0.25 − 1Hz).

The inferred modes of the dynamics can thus be grouped into three non-overlapping sets, accounting for different components in the trajectories. The first and second sets account for early and late choice-related activity, while the third set accounts for choice-independent activity. The existence of these three different components in the PFC responses likely explains why the LDS models infer dynamics that is relatively high-dimensional and involves many modes associated with relatively slow decay.

To further validate the existence of multiple phases of the dynamics, we examined activity trajectories along dimensions aligned with the relevant CD and CI components. We defined context-independent early and late CD dimensions (averaged across contexts), which are primarily aligned with the first and second set of dynamics modes (yellow and red lines in Fig. 7a), and a single CI dimension, which is primarily aligned with the third set of modes (green line in Fig. 7a).

Projections along the early CD dimension and the CI dimension reveal prominent features in the trajectories that are not apparent in other subspaces (Fig. 8a,b), confirming their potential importance in explaining the observed dynamics. The late CD dimension approximately matched the choice axis identified by Mante et al. (average angular difference: 18° across contexts, much less than chance, Extended Data Fig. 6b,c). Consistent with its alignment to slowly decaying modes, it captured a steady build-up of decision signals in both contexts over time^1^ (Fig. 8b, top panel, red dimension, dec). The early CD dimension, which aligns to more rapidly decaying modes, instead captures transient choice-related activity that emerges early in the trial, but later decays (Fig. 8b, top panel, yellow dimension, dec 2). Unlike activity along the input dimensions, which reflects the sign of a given input regardless of context (Fig. 8a,b, middle panels, black: motion coh; blue: color coh), activity along the early CD dimension only reflects the sign of the contextually relevant input, and is not modulated by the irrelevant input (Fig. 8a,b, middle vs. top panels). Finally, projections onto the CI dimension reveal components of the responses that are common to both choices (Fig. 8a,b, bottom panels). As shown above, additional dimensions capture variance due to coherence magnitude (|col| and |mot| in Fig. 4d,e, Extended Data Fig. 9a,b). All the inferred dimensions explained substantial fractions of the data variance (1-9%, Extended Data Fig. 9c) that are comparable to those captured by previously defined task-related dimensions^1^ (Extended Data Fig. 9d).

**Figure 8.**
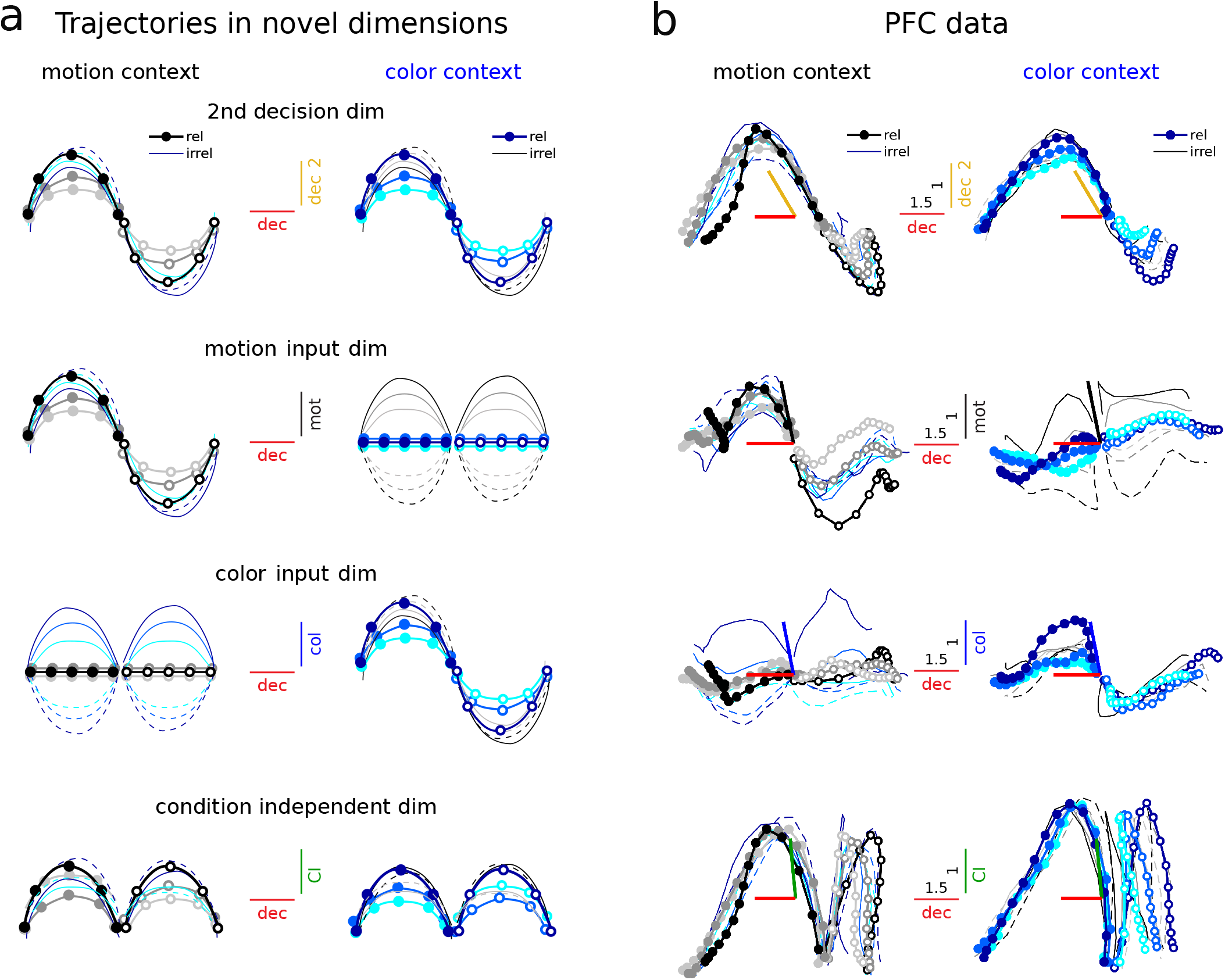
The LDS models help discover novel computational dimensions in PFC. **a**, Expected trajectories along a hypothetical secondary decision dimension (top) that reflects transient decision signals, and a dimension that captures CI signals (bottom), plotted against the evolution along a persistent decision dimension (same plotting conventions as in Fig. 3a). Contrast them with known dimensions that reflect motion and color inputs (middle, as in Fig. 3a). The new dimensions capture novel features of the population trajectories. **b**, PFC data trajectories from monkey A along the early integration (secondary decision), decision and CI dimensions (defined as the 1st svd of the CD data at the early and late periods, and the CI data at the early period, Fig. 7, averaged across contexts). Same plotting conventions as in Fig. 3b. Middle panels show the trajectories along the LDS-identified input coherence dimensions averaged across contexts and models. The data projections along them resembled the input projections found by TDR (Fig. 3b; TDR-LDS input alignments: *A, B*^*cx*^, mot = 55°, col = 42°, for mean input coherence dimensions across contexts and 100 models; *A*^*cx*^, *B*, mot = 44°, col = 31°, mean directions across 100 models; the alignments are higher than expected by chance, Extended Data Fig. 6b). CI variance has been subtracted to the trajectories in the middle panels to emphasise input-related variance. The dimensions have been orthogonalized with a QR-decomposition^1^ (starting with decision, and then dec 2, motion, color and CI). Colored bars show the alignments before the orthogonalization step (note that the inferred LDS coherence input dimensions are almost orthogonal to the decision dimension). Trajectories have been smoothed with a Gaussian filter for visualization (sliding window size, 5-bins).

Overall, these projections support the existence of two phases of integration and illustrate how dimensions based on LDS fits can isolate meaningful components of the computations implemented by the neural dynamics.

### Task-optimized RNNs do not capture all features of the PFC data

The properties of inputs and dynamics in the LDS models appear to differ in several ways from those expected from a line attractor, of the kind implemented by RNNs trained to solve the contextual integration task. Specifically, both LDS models infer multi-dimensional inputs, rely on a large number of slow dynamics modes, and process inputs in two phases (early vs. late choice dimensions). However, it is not immediately clear that these features reflect meaningful differences between the linear LDS models and the non-linear RNNs. Rather, some LDS features may reflect somewhat trivial consequences of approximating non-linear dynamics with a linear system.

We repeated all the above analyses on simulated responses of a trained RNN and found that the highlighted features of PFC dynamics are not captured by the trained RNN (Supplementary Figs. 1 to 5). The LDS fits of the RNN responses infer inputs that are largely one-dimensional (Supplementary Fig. 2c-e) and do not provide any evidence of multiple phases of input integration (Supplementary Fig. 4). In the RNN, as in PFC, the inferred slow dynamics is not limited to a single mode, unlike in a perfect line attractor (Supplementary Fig. 3a and Fig. 5b). The RNN tends to implement integration along a one-dimensional manifold that is curved^1^, rather than perfectly straight, and thus cannot be approximated by a single linear mode. Nonetheless, the number of inferred slow modes in the RNN is substantially smaller than in PFC (20 − 22% of modes are slow on average, N=100 models, Supplementary Fig. 3a; vs. 35 − 55% in the PFC data).

The analyses of RNN responses also reiterate the challenges in establishing which of the two mechanisms implemented by the LDS models is more likely to be implemented by PFC. By design, the inputs to the RNN are not modulated by context, which matches the *A*^*cx*^, *B* models, but not the *A, B*^*cx*^ models. Yet, as for the PFC data, both model classes fit the RNN data equally well (Supplementary Fig. 1). The *A, B*^*cx*^ fits of the RNN responses display some idiosyncratic properties suggestive of parameter fine-tuning, like a very large number of latent dimensions (Supplementary Table 5) and extreme levels of non-normal amplification (Supplementary Fig. 3c-e). Such fine-tuning may reflect the mismatch between the underlying mechanisms of integration. The fits of PFC responses did not display evidence of such fine-tuning, meaning that also in this respect both model classes are equally valid descriptions of the PFC data.

## Discussion

The complex and highly heterogeneous activity patterns observed in prefrontal areas are thought to be critical for the computations implemented in these regions^19^. In this study, we inferred candidate mechanisms for one such computation, contextual decision-making, by fitting interpretable LDS models directly to PFC activity. We found that two distinct mechanism of contextual-integration were consistent with the PFC responses: a switch in recurrent dynamics and a modulation of inputs. Both mechanisms required multi-dimensional inputs and high-dimensional integration dynamics to reproduce the complex dynamical portrait of PFC population activity. This finding stands in contrast to mechanisms inferred with alternative approaches that rely on models with limited complexity, which are easier to interpret but may miss potentially important features of the measured activity^1,20–24^.

The first LDS mechanism is broadly consistent with past accounts of PFC responses in this task^1,20–22^, in that the input selection relies on non-normal, context-dependent recurrent dynamics. Apart from this role in inputs selection, our analysis revealed how non-normal dynamics might additionally result in the transient amplification of relevant inputs in PFC. Non-normal transient amplification was previously proposed to be involved in the processing of external inputs^17,25–28^ and their maintenance in working-memory^29^, in producing transient activations during movement generation^30^ and in mediating robustness to perturbations^31^. Our observation of two distinct stages in PFC dynamics during decision formation is evocative of the proposal that transient amplification may optimally load information onto an attractor^27^. In contrast to such optimal loading, however, we inferred inputs that were not preferentially aligned with the most amplifying dimensions of the dynamics (Extended Data Fig. 6e-g). The loading of inputs across many modes (Fig. 5c) might alternatively reflect optimal input discrimination strategies in non-normal recurrent networks^32^.

The second LDS mechanism relies on a modest modulation of the inputs^33^. Our LDS fits reveal how strong top-down modulation of sensory areas would have to be to explain context-dependent responses in PFC. The inferred modulation strengths (38 ± 14% mot, 22 ± 8% col, Fig. 4b) are in the range of some attentional effects observed in sensory areas^34,35^, although other studies reported weaker or stronger modulation^36–40^. Unlike recent modeling of sensory and prefrontal responses during contextual-decisions in mice^40^ and humans^41^, we found that irrelevant inputs are not completely gated out before reaching PFC. Notably, not just the input strength, but also its direction was modulated by context (Fig. 4d). A change in input direction could be achieved with top-down modulation, if the input originated in multiple areas or subpopulations that are modulated independently (Supplementary Note 3). Alternatively, input amplitude and direction could both be modulated by non-linear dynamics occurring within PFC^21,23^, a possibility that we did not explicitly model here.

Both LDS models implemented input integration in two distinct phases, whereby choice-related signals first emerged along relatively fast decaying dimensions with rotational dynamics, and then transitioned towards orthogonal dimensions with slower, non-rotational dynamics. This finding is consistent with the proposal that individual task-related signals are encoded dynamically along multiple dimensions at different time-scales^12^. Indeed, our early and late choice dimensions were well-aligned with the early and middle choice dimensions in Aoi et al^12^ (35° and 22°, respectively; much more than chance, Extended Data Fig. 6b,c). Our LDS fits show how these multiple choice dimensions could emerge from the interaction of inputs and recurrent dynamics, and lead to a novel interpretation of the underlying mechanisms. Whereas such past accounts of the data concluded that recurrent dynamics in PFC is strongly rotational late in the trial^12^, in the LDS fits rotational recurrent dynamics primarily shapes the early choice responses (Fig. 7). While these features of the LDS differ in several ways from dynamics in RNNs implementing approximate line-attractors^1,16^, in agreement with such simpler models, the late choice signals predominantly emerged along a single, context-independent integration dimension^1^ (Extended Data Fig. 10).

The key features of the inferred mechanisms of context-dependent integration were consistently found across the motion and color inputs in monkey A, and the motion input in monkey F (Supplementary Figs. 6 to 10). However, as previously reported^1,12^, representations of color inputs were instead weak or absent in monkey F (Supplementary Fig. 7a-d). Unlike in monkey A, activity in monkey F also revealed evidence of strong motion integration in both contexts (Supplementary Fig. 7a,b; consistent with the observed choices in that monkey^1^); a somewhat weaker or absent separation of integration into two phases (Supplementary Fig. 9); and overall stronger CI signals, which were highly aligned with choice signals (Supplementary Fig. 6b, Supplementary Fig. 9c,d). Importantly, all these feature of activity in monkey F were captured by models that ultimately relied on similar mechanisms as those in monkey A (Supplementary Figs. 6 to 10).

The LDS models provide several insights into the properties of potential inputs into PFC, beyond their contextual modulation. First, both mechanisms inferred multi-dimensional inputs carrying information about both signed coherence and coherence magnitude. The resulting curved representation of coherence, which might arise from non-linear circuit interactions, agrees with findings in parietal and frontal areas^11,12,15^. Notably, in our models the different input components were inferred entirely from the data, rather than being hand-designed^12^. Second, both models inferred inputs that were somewhat transient, even though the fits penalized large magnitude inputs. The inputs weakened (*A, B*^*cx*^ mechanism) or progressively decayed (*A*^*cx*^, *B* mechanism) late in the trial (Fig. 4b,c). However, models with time-invariant inputs cannot be ruled out, as they performed almost as well (Extended Data Fig. 5b, Extended Data Fig. 2g,h). Critically, such models relied on mechanisms that were analogous to those described above (data not shown), confirming that the complexity of PFC responses is well approximated by linear dynamics and not necessarily inherited from inputs with rich dynamics.

Our models provide an alternative to previously proposed approaches for inferring the properties of inputs into an area. One advantage over past approaches^42–44^ is that we make minimal assumption about the properties of the inputs, like their dimensionality. Several studies have emphasized the importance of inferring inputs to understand cortical computations^11,42,45–49^ but such efforts are complicated by unavoidable model degeneracies that arise when attempting to distinguish inputs from recurrent contributions without access to the upstream areas from which the inputs originate^42,48,50,51^. Our finding that two fundamentally different mechanisms of input selection explain PFC responses equally well is a reflection of such degeneracy. Ultimately, the inferred inputs and choice-related signals may reflect computations distributed across several cortical areas^2,50^.

The LDS models explained the data essentially as well as our novel TFR model, which sets an upper bound to the goodness of fit achievable by an LDS. In PFC, intuitive linear descriptions may thus apply to all regions of state space, and not only to local regions around fixed points^1^. While we fitted activity from only a relatively short time window from each trial (the 750ms of random dots presentation), non-linear models may not outperform linear models in capturing cortical dynamics even on longer time-scales^52^. Nonetheless, analyses based on non-linear models are becoming increasingly common, given their flexibility in capturing complex neural data^42^ and in modeling biological constraints that cannot be captured by linear models^45^ (but see^31^).

A crucial aspect of our data-driven modeling approach is that it is well-suited to testing multiple alternative hypotheses about the mechanisms underlying the observed dynamics, but caution must be taken in selecting among competing theories when modeling complex systems like the brain^14,53^. Indeed, several LDS mechanisms explained the data similarly well (*A, B*^*cx*^ and *A*^*cx*^, *B* models with time-varying 3D inputs, Fig. 2a, Fig. 3c,d; 2D inputs, Extended Data Fig. 2c,d; and time-constant 3D inputs, Extended Data Fig. 5b, Extended Data Fig. 2g,h), whereas others explained the data less well (models with time-varying 1D inputs, Fig. 2a, Extended Data Fig. 2e,f) or only poorly (a *A, B* model, fully constrained across contexts, with time-varying 3D inputs, Fig. 2a, Extended Data Fig. 2b). Our best LDS models share key features with mechanisms of context-dependent integration recently inferred by a study in rats^24^, which relied on pulsatile inputs to distinguish between alternative mechanisms of input selection and integration. Similarly, the two candidate mechanisms we identified could be distinguished by their dynamics following perturbations along random state-space directions, which would differ between mechanisms due to their different degree of non-normality (Fig. 6a). Alternatively, input and recurrent contributions to the dynamics may sometimes be distinguished based on the properties of trial-by-trial variability in simultaneously recorded population responses^50^.

Methods for inferring neural population dynamics of the kind proposed here will likely play a key role in uncovering the neural computations underlying behavior. While abstract mental processes were originally hypothesized to reflect structural changes at the level of single neurons (Santiago Ramón y Cajal, see^54^), more recent evidence suggest that cognitive functions arise at the neural population level and depend critically on the ability of neural circuits to flexibly switch between dynamical regimes^55–58^. Ultimately, a complete description of neural computations will also explain how neural dynamics emerges from the rich and dynamic structural components of biological circuits^59–61^. The lawful characterization of population level dynamics amounts to a theoretical abstraction of the neural computations emerging from such a rich neural circuit, and provides a key bridge in linking lower-level biological structure to behavior.

## End Notes

## Acknowledgements

We thank Renate Krause for providing the RNN data and for valuable discussions. We thank Lea Duncker, Asma Motiwala, Saray Soldado Magraner and Gabriela Michel for providing feedback on the manuscript and for valuable discussions. This work was supported by the Gatsby Charitable Foundation.

## Author Contributions

J.S.M. and M.S. conceptualized the methodological approach, developed the models and performed data analysis. V.M. conceived the experiments, collected the data and conceptualized data analysis. All authors actively participated in the interpretation of the data. J.S.M and V.M wrote the paper.

## Code Availability

The code will be made publicly available on GitHub upon peer-reviewed publication.

## 1 Methods

### 1.1 Experimental procedures and data

#### 1.1.1 Subjects and task

Two adult male rhesus monkeys were trained in a contextual two-alternative forced-choice visual discrimination task. The monkeys had to discriminate either the color or the motion of a random dots display based on context, which was indicated by the fixation cue. The presentation of the random dots lasted for 750ms, after which the monkeys had to wait for a variable delay and report their decision. This was done by saccading to one of two diametrically opposite targets, as indicated by the color or motion evidence. The strength of the evidence was modified by varying the motion and color coherence of the random dots. This was determined by the percentage of dots moving coherently or that were colored the same. Six different coherence settings were used: three strength levels and two directions. The later indicated whether the evidence was pointing towards or away from one of two choice targets—placed at the receptive field (RF) location of the recorded neurons. When the evidence pointed towards the RF of the neurons, their FRs typically increased above baseline. Therefore, positive values were used to define the in-RF evidence. On the contrary, when the evidence pointed away from the RF of neurons, their FRs typically decreased, and hence negative values were used to define the out-RF evidence. Considering all possible motion and color coherence value pairings (6x6), 36 different random dots configurations were presented, which defined the 36 task conditions. Importantly, the motion and color evidence in a given trial could be congruent or incongruent. When incongruent, it was necessary for the monkey to ignore the irrelevant signals in order to perform the correct decision. For further details on the animal procedures and task we refer to the original study^1^.

#### 1.1.2 Neural data

Electrophysiological recordings were performed during the task in PFC regions, likely comprising the frontal eye fields (FEF) and surroundings. Both single-unit and multi-unit activity was isolated from the recordings. We referred to them as neurons, for simplicity. Only a few neurons were recorded simultaneously in each trial, but their activity was collected for multiple trials under the 36 different task conditions. Population responses were then constructed by pooling the condition-averaged activity of all neurons. For that, the firing rate of the neurons was computed in each trial using a 50ms sliding square window from spike trains sampled at 1ms. Activity was then averaged across trials under the same condition and z-scored, as in^1^. However, we did not apply any smoothing to the data prior to fitting the models (only in the analysis, for visualization purposes). Thus, the data consisted of a pseudo-population of raw per-condition averaged PSTHs. The population size was N=727 for monkey A and N=574 for monkey F. We included only neurons which had recorded activity under all conditions and for all times. As in the original study, we focused our analysis on the period of random dots presentation (750ms, from 100ms after dots onset to 100ms after dots offset) and we analysed only correct trials.

### 1.2 Models

#### 1.2.1 Linear Dynamical System (LDS) model

The linear dynamical system model considered was a non-probabilistic version of a standard LDS or state space model, with equations

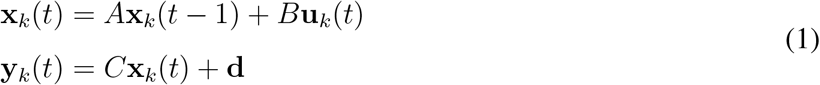

where the vector **x**_*k*_(*t*) represents the latent state at time step *t* and task condition *k*, **y**_*k*_(*t*) are the observations (a vector containing the PFC condition-averaged PSTHs) and **u**_*k*_(*t*) the external input vector. The dynamics matrix *A* determines the transition between subsequent latent states. The matrix *B* defines the input dimensions. The external inputs drive the dynamical system at each time step and define input vectors (*B***u**(*t*)) that live in the latent subspace spanned by the columns of *B*. Therefore, the external inputs are assumed linearly mixed in the population at each time step. Note that the input vectors (*B***u**(*t*)) can point in different directions over time, but these changes are always confined within the input subspaces. The input term in equation (1) can be decomposed to make explicit its color and motion components *B*_*m*_**u**_*m*_(*t*) + *B*_*c*_**u**_*c*_(*t*). The loading matrix *C* maps the low-dimensional latent state onto the high-dimensional neural space. The constant vector **d** acts as a bias. This LDS model can be seen as a low-dimensional RNN that reads-out onto a high-dimensional output space.

To capture changes in activity across contexts 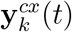, we fitted an LDS model jointly to the PFC data from each context. The model could learn independent parameters for each context (based on the data from each context) or a single parameter across contexts (using the joint data from both contexts). Both the dynamics matrix *A*^*cx*^ and the motion and color subspaces 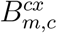 could be context-dependent (*cx* = mot or col context). The 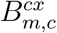 matrices could have different norms, and hence, contextual modulation of inputs could be implemented through changes in both input subspace orientation and norm. The external input signals **u**_*m,c*_(*t*) and the mapping *C* were assumed fixed across contexts.

For each motion and color input dimension, 6 external input time courses were learned, corresponding to the 6 different coherence values in the task (3 strength levels and 2 directions). These were inferred pooling data from all task conditions were a particular coherence level and direction was presented, and therefore, were shared across task conditions (i.e. there were 36 task conditions, but only 12 input traces were inferred per input dimension). The model incorporated additional input constraints, which simplified their temporal structure and were found to improve generalization performance. Time courses were constrained to be the same for all coherence levels of the same direction. That is, a single time course was shared for positive coherences (in-RF evidence) and another one for negative coherences (out-RF evidence). The coherence strength level was learned as a scalar value that multiplied the time course *u*(*t*) = *T*_*in,out*_(*t*) *coh*_1,…,6_. We also fitted a model constrained to learn fixed inputs in time, with *T*_*in,out*_(1, …, *t*) = 1. The resulting input vectors (*B***u**) for this model also live in the input subspace defined by the input matrices *B*, but unlike the input vectors for the time varying input model (*B***u**(*t*)), these do not move within the input subspaces over time, and remain fixed throughout the trial (both in strength and direction).

A different vector of initial conditions was also learned for each context 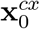. This parameter helped the model recreate the separation of trajectories in state space found across contexts (contextual axis in the Mante et al. study^1^). Note that this feature cannot account for the contextual differences in input integration, since the model is linear, so the relationship between inputs and dynamics modes is the same everywhere in state space. Indeed, a fully constrained model across contexts, with flexibility only in the initial conditions, fails to selectively integrate and poorly reproduces the data (Fig. 2a *A, B* model, Extended Data Fig. 2b). The initial conditions simply add a shift to the overall dynamics in an input independent manner, since **x**_0_ is the same across all task conditions, so it could only capture baseline changes across contexts. This can be seen in the next equation, which illustrates the unfolding of the dynamics from the initial state and makes the dynamics and inputs convolution explicit

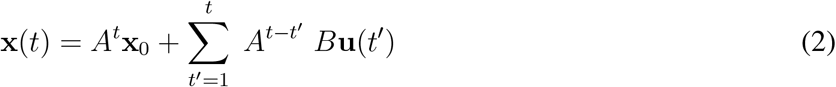

This equation also illustrates the presence of a summation degeneracy in the model. The first term defines condition independent (CI) effects, but these can also be captured by the input term. For this reason, in figure Fig. 4, Fig. 8, Extended Data Fig. 3b, Extended Data Fig. 4, Extended Data Fig. 9 and associated Supplementary Figures, we subtracted out CI effects from the input/data trajectories along the input dimensions.

The model was implemented in Python and optimized using gradient descent (ADAM algorithm) to minimize the data reconstruction mean squared error (MSE), with learning rate of 0.009, and the rest of parameters set to the default. The convergence criteria was set to ΔMSE *<* 10^−5^, maximum iterations to 10,000 and minimum iterations to 5,000. The cost function incorporated an input norm penalty to constrain the space of possible solutions and to favour learning small inputs. This encouraged that task-related variables in the data other than the inputs, in particular integration signals, were generated dynamically by the model.

Incorporating the penalty minimally impacted performance and helped provide consistent solutions across fits even when parameters were initialized at random. Therefore, we incorporated such penalty in all our model fits and randomly initialized all parameters. The resulting objective function was:

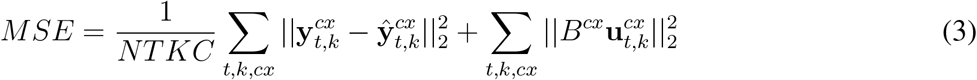

Where N = number of neurons, T = trial duration, K = number of conditions and C = number of contexts. Since the data was z-scored, the MSE captured the fraction of unexplained variance in the data by the model.

Note that the LDS was simply optimized to minimize the MSE of the condition-averaged PSTHs. We did not learn any observations noise model or inferred a latent state distribution, contrary to more standard formulations of the LDS, which are fully probabilistic (and typically infer Gaussian latents, or Gaussian latents combined with Poisson observations^62^). We considered this simpler case given that our data was trial-averaged. Furthermore, our focus was to analyse the parameters of the dynamical model, which are part of the prior distribution over the latents in the probabilistic LDS, and not the data-corrected posterior distribution.

#### 1.2.2 Tensor Factor Regression (TFR) model

The model consists of a factorization of the data tensor structure into three main low-rank tensors

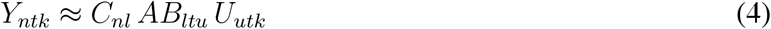

where *n* = number of neurons, *t* = time steps, *k* = conditions, *l* = latent dimensionality, *u* = input dimensionality + 1D baseline. The tensor *C* (an orthonormal matrix) sets the rank of the factorization and maps the low-dimensional core tensor *AB* into the high-dimensional neural space. The inputs tensor *U* captures the condition-dependent effects in the data and acts as a regressor, when this is known. When learned, as it is the case here, it is used to capture task-related variables, such as motion and color input signals. Note that similar to the LDS, these signals are assumed linearly mixed in the population at each time step.

In the previous equation, for clarity (as in Fig. 2c), we omitted an indicator tensor *T* that emulates the LDS-like convolution of inputs and dynamics

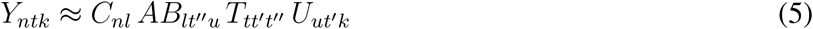

where *T*_*tt*′t″_ = *δ*(*t* − *t*′ = *t*″). One can see how this model encompasses the LDS by writing

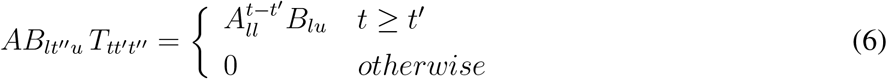

where *A* and *B* correspond to the LDS dynamics and input subspace matrices, respectively.

The inputs incorporated constraints analogous to the LDS. First, inputs were repeated across conditions with an additional indicator tensor *Q*

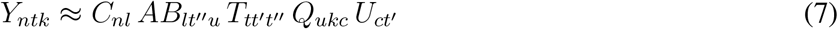

where *c* = (6 × *u*) + 1 indexes the six coherence conditions, plus baseline (that captures CI effects). In this way, the tensor *U* is designed to extract common task-related variables across conditions. Second, the temporal structure of the inputs was constrained to be the same for coherences of the same direction. For that, the input tensor *U* was factorized further as follows

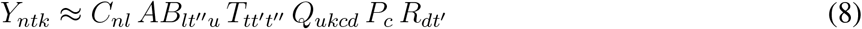

where *d* = 2 indexed the two possible coherence directions.

The parameters of the TFR model can be computed by alternating the estimation of the tensors *W* = *CAB* and *U* . For that, one can consider the tensor unfolding *Y*_(*n*)(*tk*)_ and compute *C* and *AB* via reduced rank regression, with fixed *U* . Then, knowing *W*, the least squares estimate of *U* can be computed. In practice, we estimated the parameters following the same optimization procedure we used for the LDS, which provided identical results. That is, the model was implemented in Python and optimized using ADAM, with objective given by the data reconstruction MSE.

The TFR model is related to existing regression-based methods that discover task-related variance in the data^1,12,63^, but with the difference that TFR incorporates task regressors that are themselves learned from the data. Another key distinction is that TFR considers a joint factorization of the whole data tensor structure, similar to other studies^64^, but the tensor components relate to the parameters of the task and are themselves low-dimensional.

#### 1.2.3 Recurrent Neural Network (RNN) model

We generated data from a RNN model of the same type as used by Mante and colleagues^1^.

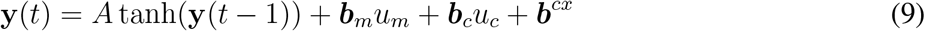

Briefly, the model was a non-linear RNN trained using back-propagation to solve the same contextual decision-making task as the monkeys. Contrary to the LDS, the RNN was not optimized to reproduce the complex and heterogeneous responses of PFC neurons, i.e. to match PFC’s dynamics. This network was designed with the same built-in assumptions as in the original model (Fig. 1c). Namely, that the external coherence input signals *u*_*m*_ and *u*_*c*_ were noisy but constant in time, with mean proportional to the strength of the coherence evidence, and that these reached the circuit through two fixed input dimensions across contexts **b**_*m*_ and **b**_*c*_. The model had the flexibility to learn different contextual input vectors **b**^*cx*^, whose activation changed the dynamics of a fixed, non-linear recurrent network (with connectivity *A*). This allowed the model to switch its state between two approximately linear regimes 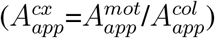, performing different computations in each context. Namely, selecting the contextually-relevant input signals for integration towards a choice and dynamically discarding the irrelevant ones. In the original study, the RNN population activity 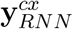 was analysed and qualitatively compared with the PFC activity, revealing some shared features that were suggestive of a common contextual-integration mechanism between PFC and the network. The network could be “reverse-engineered” in order to understand the mechanism underlying such computation, by linearizing the dynamics around the identified fixed points of the system (obtaining different local dynamics matrices 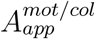, which however were similar in dynamics and could be averaged^1^). In this work, we instead focused on analysing the properties of LDS models fit to the RNN population activity 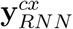 (the z-scored condition-averaged responses, as in the PFC data, but from 100 RNN units), and recovered one or two dynamics matrices (*A*^*mot/col*^ in the *A, B*^*cx*^ model; *A*^*mot*^ and *A*^*col*^ in the *A*^*cx*^, *B* model) that approximated the global dynamics of the RNN population in both contexts. For further details on the RNN training and analysis we refer to the original study^1^.

### 1.3 Dynamics analysis

#### 1.3.1 Eigenspectrum and time constants

The eigenspectrum of the LDS dynamics matrices contains both real and imaginary eigenvalues (Extended Data Fig. 6a), which come in complex-conjugate pairs

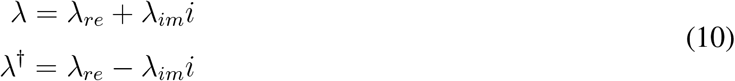

The absolute value of the eigenvalues determines the rate of decay or growth of each dynamic mode^65^. Modes are stable if they either decay or persist

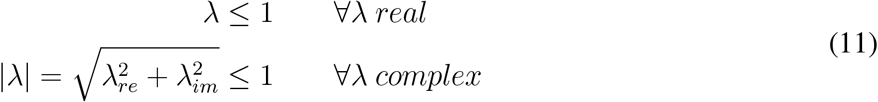

The slower the decay, the slower or more persistent a given mode is, and the greater input information is preserved along it. The time constant measures the time at which the initial state will have decayed by 37% (1/e=0.37) along a given mode. Considering that each time step is 50ms (the data binning size)

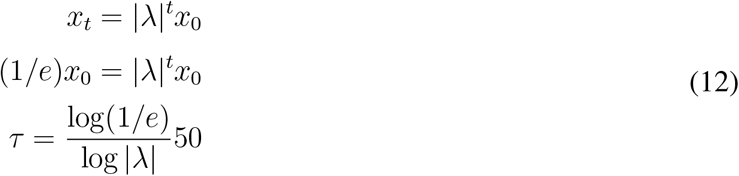

We classify a mode as slow if it has a norm close to one, that is, if |*λ*| *>* 0.8. This corresponds to a decay time constant of *τ >* 224ms, which encompasses approximately a third of the trial duration. Given that the inferred external inputs in the two models are strong for the first third of the trial (Fig. 4a,b), inputs mapped onto such slow modes largely persist until the end of the trial, albeit with some decay for modes |*λ*| = 0.8 − 0.9. In particular, by the second third of the trial, inputs would have decayed by at most 37%. We consider the slowest modes to have |*λ*| *>* 0.9 and time constant *τ >* 475ms. These are strongly persistent and preserve most input information until the end of the trial. The relatively fast decaying modes (|*λ*| = 0.7 − 0.8, *τ* = 140 − 224ms) are somewhat persistent, but loose most input information by the end of the trial.

Many of the eigenvalues were imaginary, indicating the presence of rotational dynamics in the data^56^. Some of the eigenvalues were negative, which also indicate the presence of oscillations^66^. A few models identified slightly unstable eigenmodes (with eigenvalue norm slightly bigger than 1), but this is expected when learning from finite trial lengths and limited data samples^67^. However, the models inferred from monkey F data, in particular for the *A, B*^*cx*^ model, seemed to use instability properties of the dynamics in order to capture specific features of the data (Supplementary Fig. 8a,d).

#### 1.3.2 Rotational dynamics measure

As mentioned above, the existence of complex eigenvalues indicates the presence of rotational dynamics in the data. Rotations are confined to the planes defined by pairs of complex-conjugate eigenvectors, with directions spanned by the real and imaginary components of the vectors. On each plane, state trajectories are shaped by the rotation matrix *J*, which derives from the dynamics matrix *A* expressed in the Jordan normal form^65^. As an example, for a 2d system with two distinct complex eigenvalues, which come as a complex-conjugate pair *λ* − *λ*^†^ (equation (10)), if we consider their phase plane representation in polar coordinates

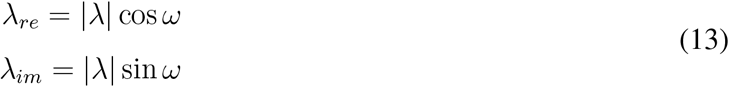

where

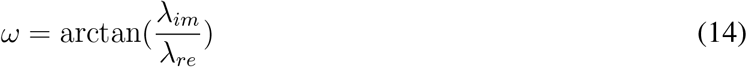

the rotation matrix *J* is given by

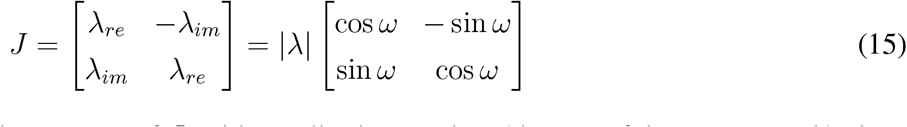

Rotations evolve in time following powers of *J*, with amplitude over time (the rate of decay or growth) given by the absolute value of the eigenvalues, and with rotation frequency *ω*

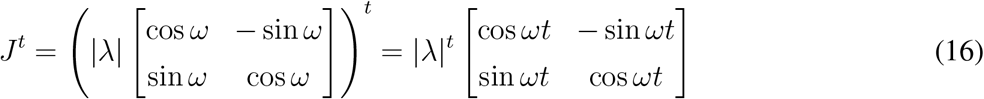

Note that the frequency increases when the ratio 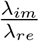 is big. The rotation frequency *ω* is given in *rad/s* and *f* = *ω/*(2*π*) in Hz. Since the data was down-sampled at 20Hz (50ms bins), the frequency is given by *f* = 20*ω/*(2*π*) in Hz (the value reported in Fig. 7). For real modes, the rotation frequency is zero.

#### 1.3.3 Non-normality measure

The Henrici’s index measures the degree of non-normality of the the dynamics, and is given by^68^

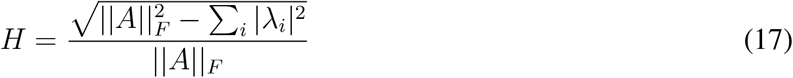

This is a normalized metric with values between 0 and 1, with 0 indicating that the system is normal, and 1 that is maximally non-normal. A system is normal when its dynamics can be described with an orthonormal eigenvector basis. A system is non-normal when its eigenvectors do not necessarily form an orthonormal basis, and the transformation to eigenvector coordinates may involve a strong distortion of the phase space^68^. Importantly, in normal linear networks, the network responses are explained with a linear combination of exponentially decaying modes (if the system is stable), with timescales defined by the corresponding eigenvalue (equation (12)). In non-normal stable networks, however, more complex patterns can emerge, which often involve transient responses where the network activity temporarily grows, but eventually decays as in normal systems.

A crucial property of non-normal systems is that they have different left and right eigenvectors.

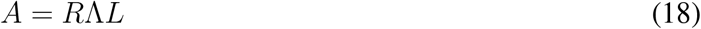

with *L* = *R*^−1^, whereas for normal systems *L* = *R*^†^ (†=conjugate transpose). This non-normal property allowed the RNN to change the leading left eigenvectors across contexts, while keeping the right eigenvectors pointing in the same direction^1^.

#### 1.3.4 Input loads

To compute the input load onto the modes of the dynamics, we start by expressing the latents in the left eigenvectors basis.

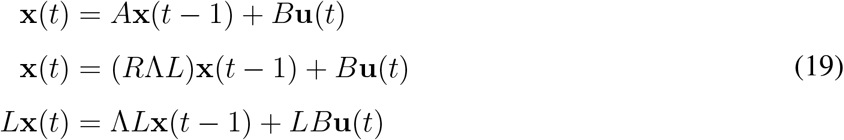

where we have taken the eigendecomposition of the matrix *A*, with *R* containing the right eigenvectors in its columns and *L* = *R*^−1^ the left eigenvectors in its rows. We have then left-multiplied by *L*. Defining ***α***(*t*) = *L***x**(*t*) we obtain

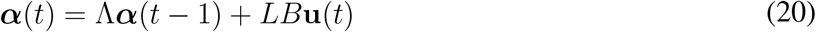

The evolution of the latents in this basis is independent, that is, decoupled from one another—given that the matrix Λ is diagonal. Unrolling this equation in time we obtain

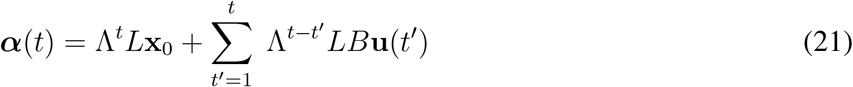

As the eigenmodes are independent, we can write down a set of uncoupled equations that describe the evolution of each eigenmode, one for each entry of the vector ***α***, given by *α*_*l*_ with l indexing the latent variable dimension

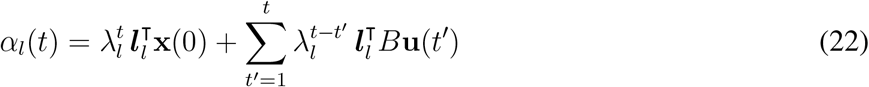

and ***l***_*l*_ being the lth left eigenvector. The input “loads” are defined by the last term of the summation, which correspond to the non-normalized projection of the inputs onto the left eigenvectors (given that neither the input vectors nor the left eigenvectors are unit norm).

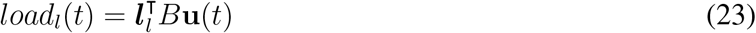

This term specifies how strongly the inputs are mapped onto the dynamic modes, at each time step *t*, before being processed by the dynamics (i.e., in this basis, before being scaled by *λ*). The extend to which the inputs are mapped or “loaded” onto each mode depends on the alignment between the input vectors and each left eigenvector, as well as the norm of both vectors. For each pair of complex modes, the load is given by

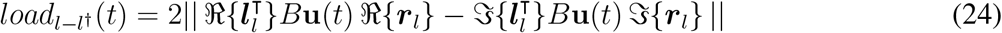

Where *ℜ{*.*}* and *𝔍{*.*}* take the real and imaginary components of their arguments. The rationale for the expression above comes from the following. For complex modes, equation (23) contains imaginary numbers, since the left eigenvectors are complex, so we cannot interpret the loads in this basis. However, we can do it in the original state vector basis **x**(t), which is real. To change basis, we use ***α***(*t*) = *L***x**(*t*) and express **x**(*t*) as a linear decomposition of the state along each right eigenvector dimension. The coefficients of the linear decomposition are given by *α*_*l*_(*t*), which contains the input loads

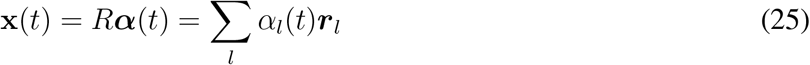

We can now make explicit the contribution due to real eigenmodes and complex eigenmodes, which come in complex conjugate pairs (*l* − *l*^†^).

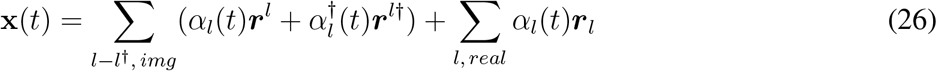

Due to the complex conjugacy, the imaginary numbers end up cancelling out in the summation, and only real terms survive. This is why in this basis, the state vector **x**(t) is real. In particular, the way the complex roots end up contributing to the state dynamics is given by their real and imaginary parts. This is because for each pair of complex conjugate roots, two complementary real solutions exist, which are given by the sum and difference modes 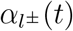

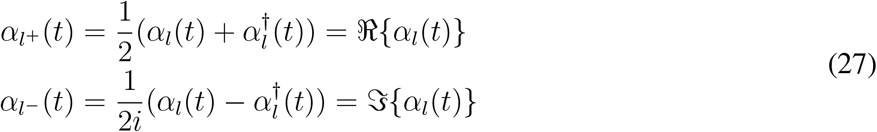

This can be seen by expanding the complex term in the state equation

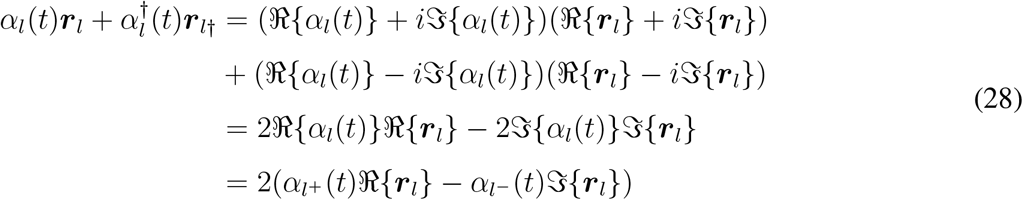

Thus

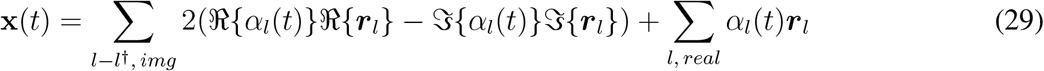

To understand how the inputs are loaded at each time step *t* into the dynamic modes to affect the latent state, we focus on the last term of the summation in the *α*_*l*_(*t*) equation (22), as we did before

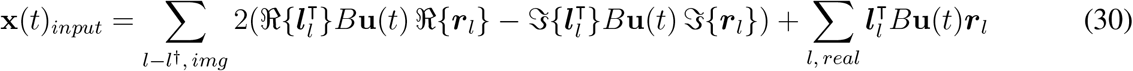

The last term contains the input loads along each real mode, 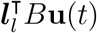, which gives equation (23). This value indicates how much of the input is mapped along each right eigenvector direction ***r***_*l*_ (for *l* real). Thus, considering only this term, the latent state vector is reconstructed with a linear combination of real right eigenvectors, weighted by the input loads. Note however, that the right eigenvectors are not orthogonal, so the result of the sum could be non-trivial, if for instance some of this vectors cancel out, or give rise to amplification (Supplementary Note 2, Extended Data Fig. 7). The total input contribution or load along each direction ***r***_*l*_ is thus given by the norm of the vector 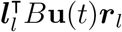. Since the real right eigenvectors are normalized, this is equal to 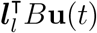, which gives equation (23). Similarly, the load for each complex conjugate pair of modes is given by the norm of the vector 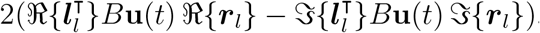, which gives equation (24). This vector lives within the 2D plane spanned by the real and imaginary components of the complex-conjugate right eigenvector pairs.

To compute the loads in Fig. 5c, we use the inferred inputs for the largest motion and color positive coherence values, and project them along the coherence dimension. Thus, the loads are computed using the coherence component of *B***u**(*t*), for all times and all 100 randomly initialized models, and then averaged across time and models. For complex modes, the same load is shared across both complex conjugate pairs, and is computed using equation (24).

#### 1.3.5 Most amplifying dimensions

The most amplifying modes were found following^27^, by computing the Observability Gramian and its associated eigenvectors. The most amplifying modes are defined by the eigenvectors with the largest associated eigenvalues. We computed the Observability Gramian by solving the following Lyapunov equation

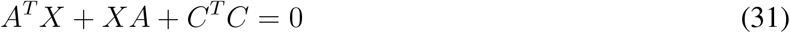

where A is the LDS models dynamics matrix and C is the loading matrix. We considered only stable models^27^, which in our case were 90% of the 100 *A, B*^*cx*^ models and 85% (mot cx), 60% (col cx) of the *A*^*cx*^, *B* models in monkey A.

### 1.4 Additional analysis methods

#### 1.4.1 Alignment metrics

We report alignments between different dimensions using either dot products or angles (in degrees). When computing alignments between a given vector and complex eigenvector dimensions, we consider the plane spanned by the real and imaginary vector components of the pair of complex conjugate modes, and compute the minimum subspace angle between the vector and the plane.

#### 1.4.2 Statistical tests

To asses statistical significance of differences between distributions, such as the relevant vs. irrelevant load distributions in Fig. 5c, we used a Wilcoxon rank-sum test with significance levels (p-values) set at p*<*0.001 (Fig. 4b,c, Fig. 7b, Extended Data Fig. 8b, and associated Supplementary Figures) or p*<*0.05 (Fig. 5c and associated Supplementary Figures, also Supplementary Fig. 4). This is a two-sided rank sum test of the null hypothesis that two independent samples come from distributions with equal medians.

---

**Extended Data Figure 1.**
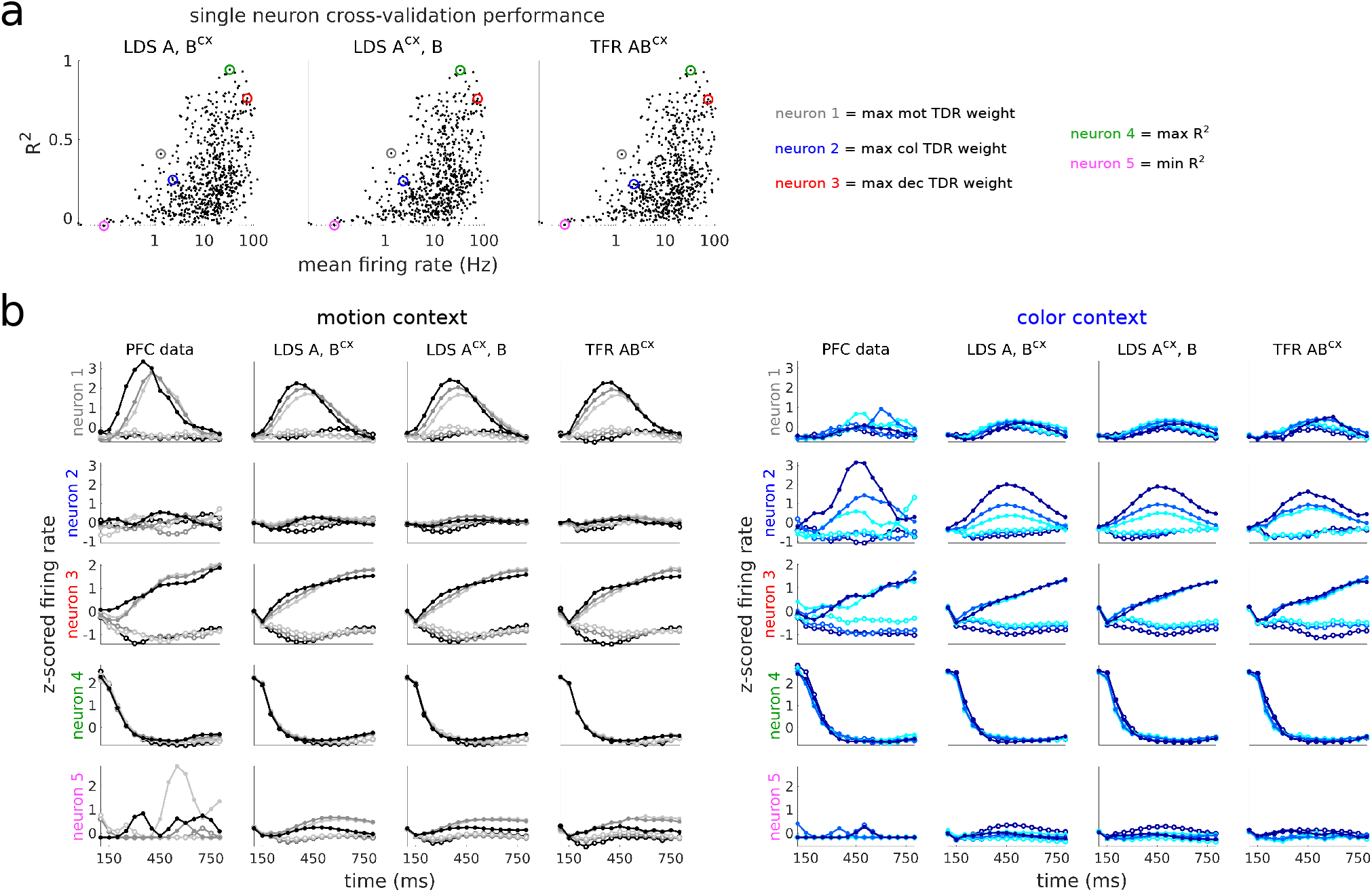
Individual neuron performance for the best LDS and TFR models. **a**, LDS and TFR models *R*^2^ for all individual neurons, sorted by their mean firing rate as in Aoi et al.^12^. Highlighted in colors are five example neurons. Three of them have maximum selectivity to either motion, color or decision (in grey, blue, red), as measured by their weight onto the motion, color and decision population vectors found using targeted dimensionality reduction (TDR)^1^. The two other neurons were selected based on model performance (best neuron captured, max *R*^2^, in green, and worse neuron captured, min *R*^2^, in pink). **b**, PFC data and LDS/TFR models cross-validated PSTHs for the 5 example neurons. The PSTHs are computed from z-scored data and model responses sorted by the relevant coherence value in each context (motion in the motion context, left, and color in the color context, right) and averaged across irrelevant coherence conditions, as in Mante et al.^1^. Color shades and filled/hollow circles indicate the strength and direction of the coherence evidence, respectively (same notation as in Fig. 1a). PFC data responses have been smoothed with a Gaussian kernel (*σ*=40ms) for visualization^1^. Data is from monkey A.

**Extended Data Figure 2.**
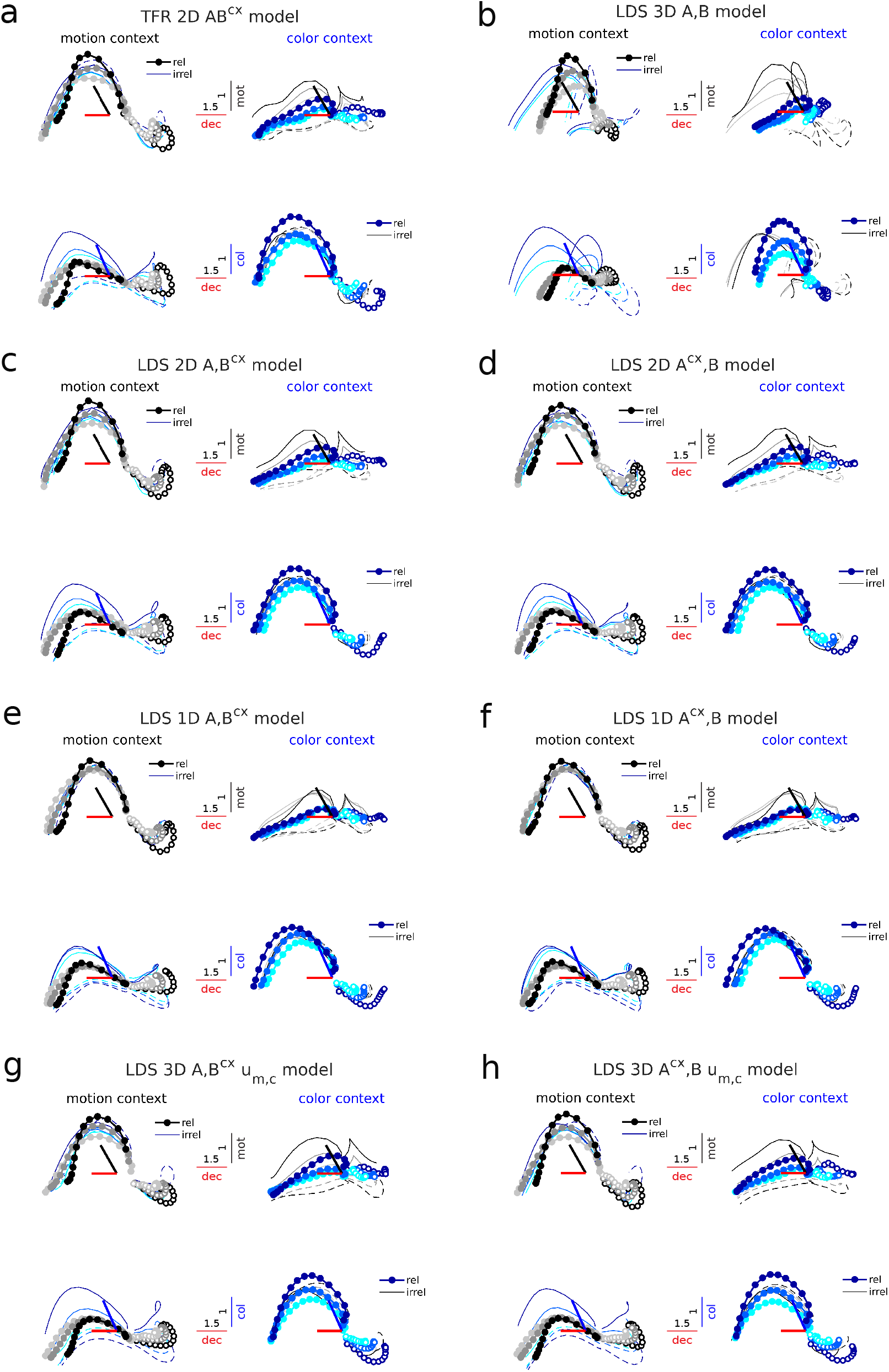
Population trajectories of the TFR model and alternative LDS models in the task-related subspace. Cross-validated model trajectories (LOOCV) for the best TFR model and additional LDS models with various input dimensionalities and input/contextual constraints. **a**, TFR *AB*^*cx*^ model with 2D inputs. **b**, LDS *A, B* model with 3D inputs. This model poorly captures the trajectories, specially along the decision dimension. **c**,**d**, LDS *A, B*^*cx*^ and *A*^*cx*^, *B* models with 2D inputs. These capture the trajectories almost as well as the 3D models (Fig. 3c,d) **e**,**f**, LDS *A, B*^*cx*^ and *A*^*cx*^, *B* models with 1D inputs. Note that the trajectories along the input dimensions are not accurately captured, in particular for the irrelevant inputs, where trajectories poorly separate by coherence condition. **g**,**h**, LDS *A, B*^*cx*^**u**_*m,c*_ and *A*^*cx*^, *B***u**_*m,c*_ models with time-constant 3D inputs. These also perform nearly as well as the 3D LDS models with time-varying inputs (Fig. 3c,d, Extended Data Fig. 5b). Same conventions as in Fig. 3. All trajectories have been smoothed with a Gaussian filter for visualization (sliding window size, 5-bins). This step did not change much the LDS trajectories, since they are inherently smooth, but it helped smooth-out substantially the TFR model trajectories, given that this model has no dynamical constraints that enforce smoothness. Monkey A data.

**Extended Data Figure 3.**
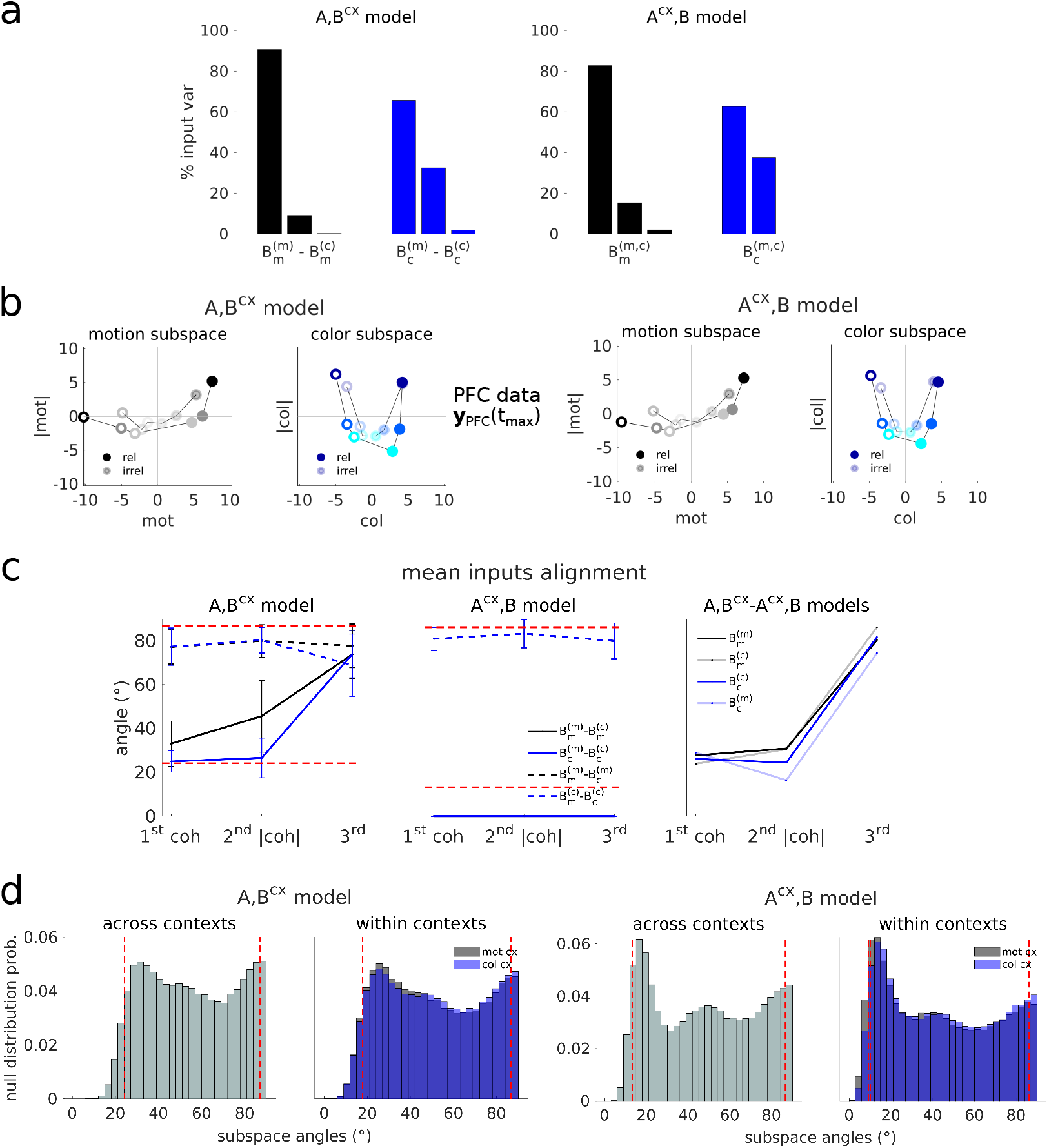
Input variance, PFC data in the 2D input subspaces and alignment statistics. **a**, External inputs variance in the three orthonormalized input dimensions from both LDS models. For the *A, B*^*cx*^ model variance is computed along the mean dimensions across contexts (first taking the average across contexts: mot avg. 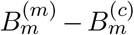, for dims 1-3, and col avg. 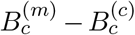, for dims 1-3, then re-orthogonalizing the three input dimensions). Note that most of the input variance is concentrated in the first two input dimensions (the 2D plane capturing coherence an coherence magnitude variance, found via regression, and then orthonormalised using a QR factorization^1^) whereas the third dimension carry almost no input variance (see also Extended Data Fig. 4c,f, top panels). Averages across 100 models. **b**, PFC data at t=250ms for all coherences and both contexts, projected onto the same 2D coh-|coh| input planes as in Fig. 4d,e. The PFC data contains a curved representation of coherence information. CI signals have been subtracted. **c**, Alignment between the motion and color input vectors within contexts (dashed lines), and between the motion or color input vectors across contexts (filled lines), for each of the three input dimensions, and the two LDS models (left and middle panels). Dashed red lines, 5th and 95th percentiles of a null distribution of alignments (see **d**). Error bars, std across a 100 randomly initialized models. Note that the inferred coherence and coherence magnitude dimensions for both motion and color are largely stable across contexts in the *A, B*^*cx*^ model (i.e. they are highly aligned, filled lines). However, the across-contexts alignments for the third input dimension are close to orthogonal, indicating that this dimension is not common across contexts. Right panel, alignments between the mean input directions (across 100 models) from each LDS model class. The coh and —coh— directions are highly consistent across models. **d**, Null distribution of alignments (subspace angles) from randomly sampled 3D subspaces within and across contexts drawn aligned to the data covariance^69^, orthonormalized, and then projected onto the low-d subspaces from each LDS model class (defined by the columns of the loading matrices *C*). The null distributions from the two model classes are different since they learned different *C* matrices. Random samples s≈33,000 orthonormal subspaces, or 100,000 vectors. Dashed red lines, 5th and 95th null distribution percentiles. Alignments not expected by chance fall in regions ≤ the 5th or ≥ the 95th percentiles of the control distributions. Additionally, given the binomial nature of the distribution, where both high and low alignments are expected, random alignments should on average lie around 50° and have large variance. This is not what is typically obtained from the data (panel **c**). Furthermore, in all 100 models from the *A, B*^*cx*^ class, the highest alignments consistently occurred between the two color and two motion dimensions across contexts, and not between the motion and color dimensions within contexts (first panel in **c**, filled vs. dashed lines). This was true only for the first two input dimensions, but not the third, where all alignments are very low. Similarly, the first two mean input dimensions were highly aligned across model classes, but not the third (**c**, third panel). Monkey A data.

**Extended Data Figure 4.**
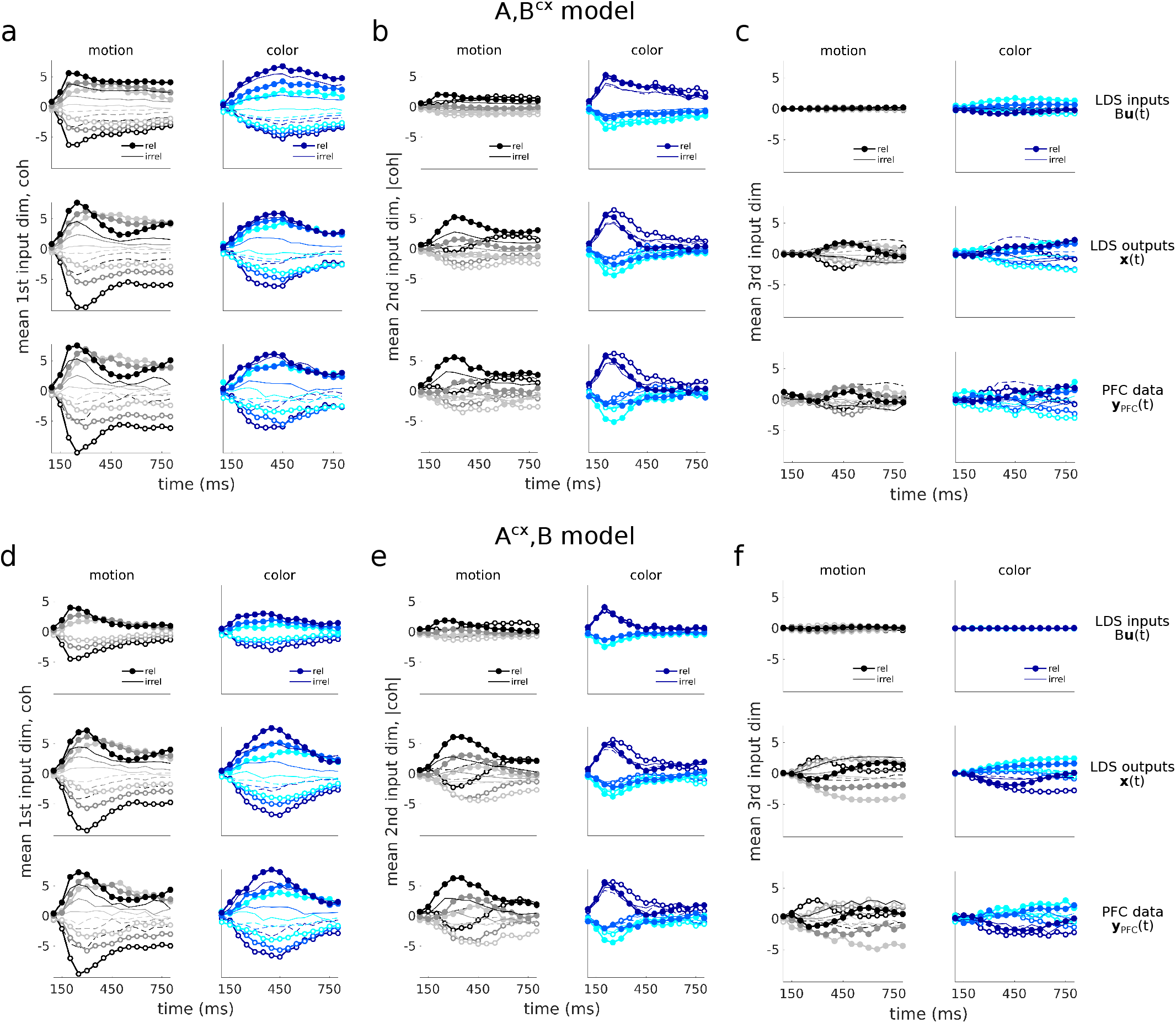
LDS models external inputs, latents and PFC data in the LDS input dimensions. LDS external inputs, LDS cross-validated latents (outputs) and PFC data trajectories projected along the three input dimensions found in the *A, B*^*cx*^ (**a-c**) and *A*^*cx*^, *B* (**d-f**) models, for all coherence conditions and contexts (relevant vs. irrelevant). **a**,**d**, First input dimension, capturing coherence related variance (coh). **b**,**e**, Second input dimension, capturing coherence magnitude related variance (|coh|). **c**,**f**, Third input dimension, orthogonal to the coh and |coh| dimensions, capturing little input and relatively little output/data variance compared to the other dimensions (in particular, for color). For the *A, B*^*cx*^ model, projections are shown onto the direction bisecting the two color and two motion directions found for each context. All data is from means over 100 models initialized at random. The three mean color and motion input dimensions are orthogonalized with QR-factorization. For all trajectories the mean across conditions has been subtracted out to remove condition independent signals (CI). Latents and data trajectories are generated for all 36 task conditions and plotted along the motion/color input dimensions with color/motion conditions averaged out, as in^1^. Same plotting conventions as in Fig. 3. Monkey A data.

**Extended Data Figure 5.**
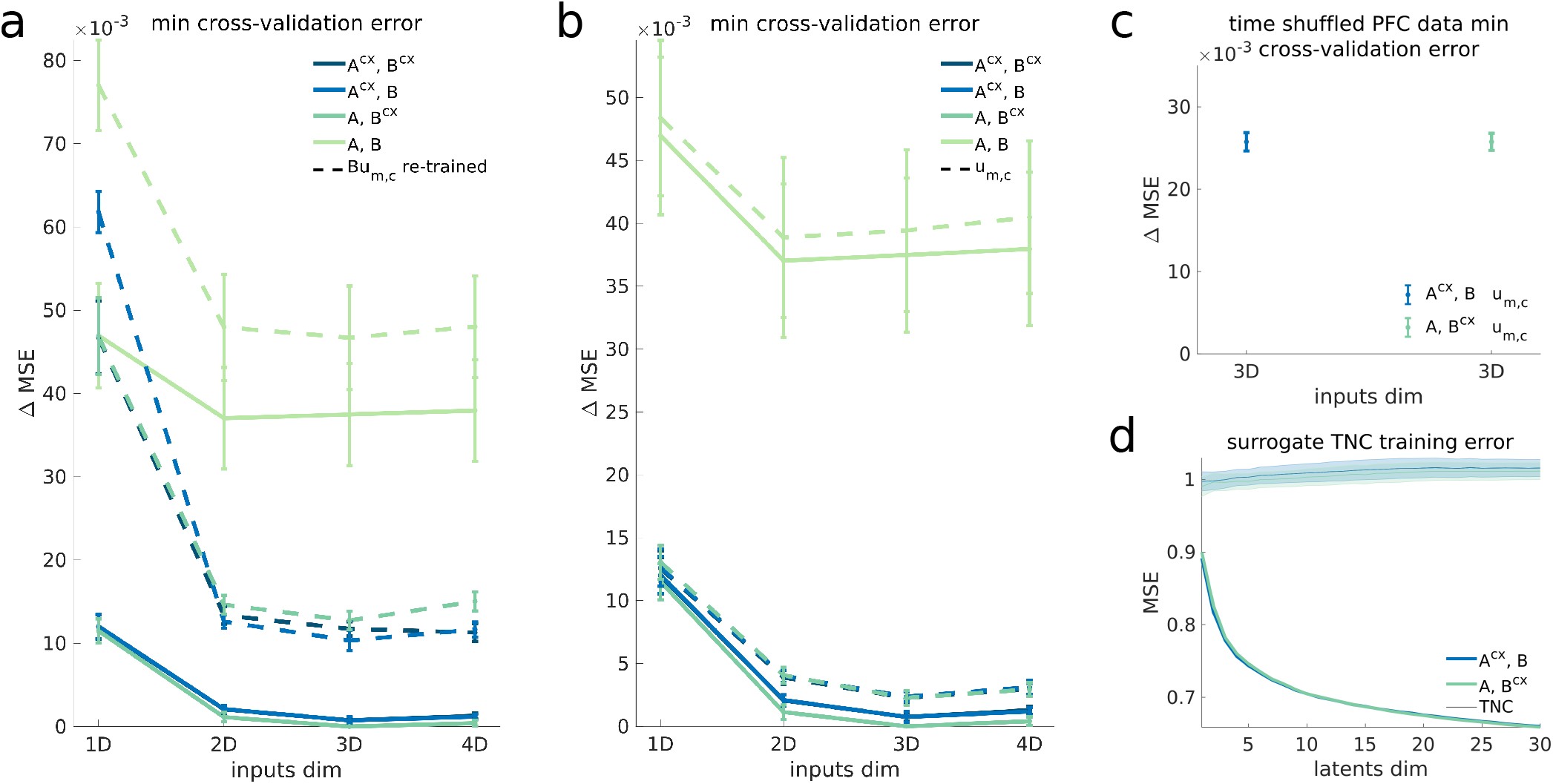
Performance of LDS models with time-constant inputs and randomized data controls. **a**, Leave-one-condition-out cross-validation performance (LOOCV) of the LDS models with time-varying inputs in Fig. 2a (filled lines) and the same models after re-training their input parameters *B***u**_*m,c*_, but constraining **u**_*m,c*_ to be constant in time, and with the rest of the parameters kept the same (dashed lines). See next panel for performance of a similar model but fully optimized (all parameters re-trained). **b**, Performance of LDS models with **u**_*m,c*_ constant in time where all parameters, including the dynamics matrix, are optimized to fit the data (dashed lines). The time-varying input models from Fig. 2a are also shown for reference (filled lines). **c**, Performance of the best time-constant models (*A, B*^*cx*^**u**_*m,c*_ and *A*^*cx*^, *B***u**_*m,c*_ 3D models, dashed lines in **b**) when fitted to time-shuffled PFC data. Note that the performance drops substantially (Δ MSE = 26), being worse than the 1D input models and nearly as bad as the most contextually constrained *A, B* models (see **b**). For all three subpanels (**a**-**c**) the minimum cross-validation errors are shown relative to the best performing LDS model (the time-varying *A, B*^*cx*^ model with 3D inputs). Error bars indicate the standard error mean across LOOCV folds. **d**, Training performance of the best time-varying LDS models on surrogate data sets randomized across time, neurons and conditions (TNC), but designed to preserve the primary statistics of the data^13^. The LDS models perform poorly on these type of data. To obtain the randomized TNC data sets the tensor maximum entropy method (TME) was used. Shades indicate standard error mean across 30 surrogates. Monkey A data.

**Extended Data Figure 6.**
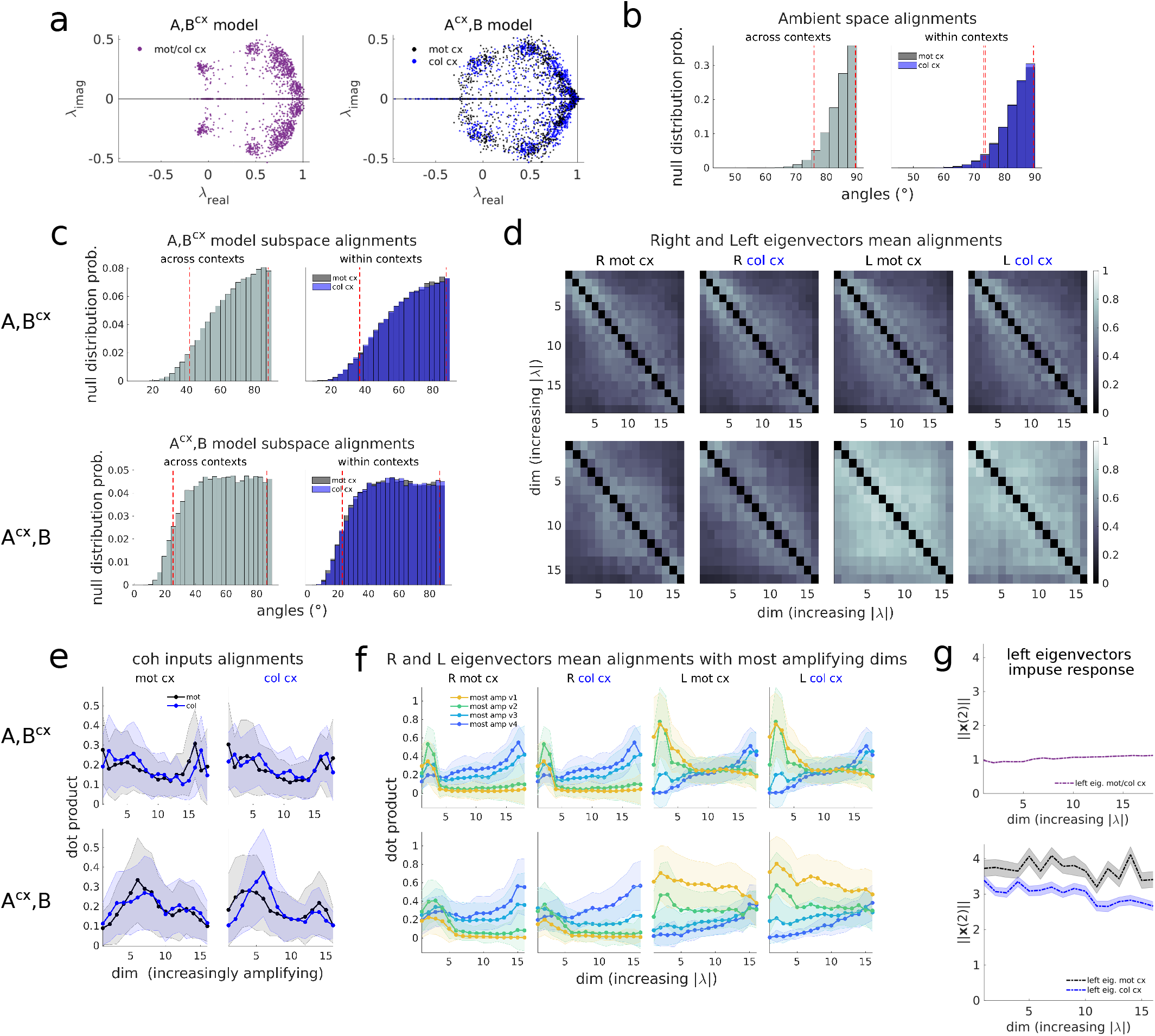
LDS models dynamics properties. **a**, LDS eigenspectrums for the 100 randomnly initialized models from each model class. **b**,**c**, Null distribution of alignments for randomly sampled pairs of vectors, within contexts or across contexts, drawn aligned to the data covariance^69^ in the ambient (high-d) space (**b**) or projected onto the low-d LDS subspaces from each model class (**c**, defined by the columns of the loading matrices *C*). Random samples s=100,000 vectors. Dashed red lines, 5th and 95th null distribution percentiles. Alignments not expected by chance fall in regions *<* the 5th or *>* the 95th percentiles of the control distributions. **d**, Mean alignments among the left/right eigenvectors from each model class. *A, B*^*cx*^, top, *A*^*cx*^, *B*, bottom, averages across 100 models from each class. Note that the left eigenvectors in the *A*^*cx*^, *B* model are highly aligned, which explains its strong non-normal dynamics (Fig. 6c). **e**, Motion and color input coherence vector alignments with respect to dynamic dimensions of various degrees of amplification, sorted from the least to the most amplifying modes (Methods), for the two contexts (left and right panels) and the two LDS model classes (top, bottom). Mean ± std across 100 models. Motion and color inputs do not strongly align with the most amplifying dimensions. Note that in Fig. 5c we found that coherence inputs were strongly loaded onto the relatively fast decaying left eigenmodes. Indeed, these intermediate left eigenmodes do not align particularly strongly to the most amplifying modes, compared to the alignments for the fastest and the slowest eigenmodes (see **f**, L panels). **f**, Right (R) and Left (L) eigenvectors alignments with respect to the four most amplifying modes of the dynamics (Methods), for both models and both contexts. Mean ± std across 100 models. Note that all left eigenvector dimensions, from fast, to intermediate, to slow, have moderate to strong alignments with the most amplifying modes. In fact, all left eigenvector directions amplify inputs similarly (see next panel) **g**, Impulse response along each left eigenvector direction, measured at the time right after the unit norm perturbation (t=2). The state at this time indicates the degree of transient amplification immediately after the pulse (see also Fig. 6a for response over time, averaged across all left eigenvectors). The state norm at t=2 is slightly bigger than 1 for all *A, B*^*cx*^ left eigenvectors, indicating that all these directions slightly amplify inputs. For the *A*^*cx*^, *B*, in both contexts, the impulse response is much bigger than one for all left eigenvectors, indicating that all these directions strongly amplify inputs. This confirms that the dimensions where the inputs are mostly loaded, the intermediate or relatively fast decaying dimensions, are not particularly amplifying, relative to the fastest and slowest dimensions. Monkey A data.

**Extended Data Figure 7.**
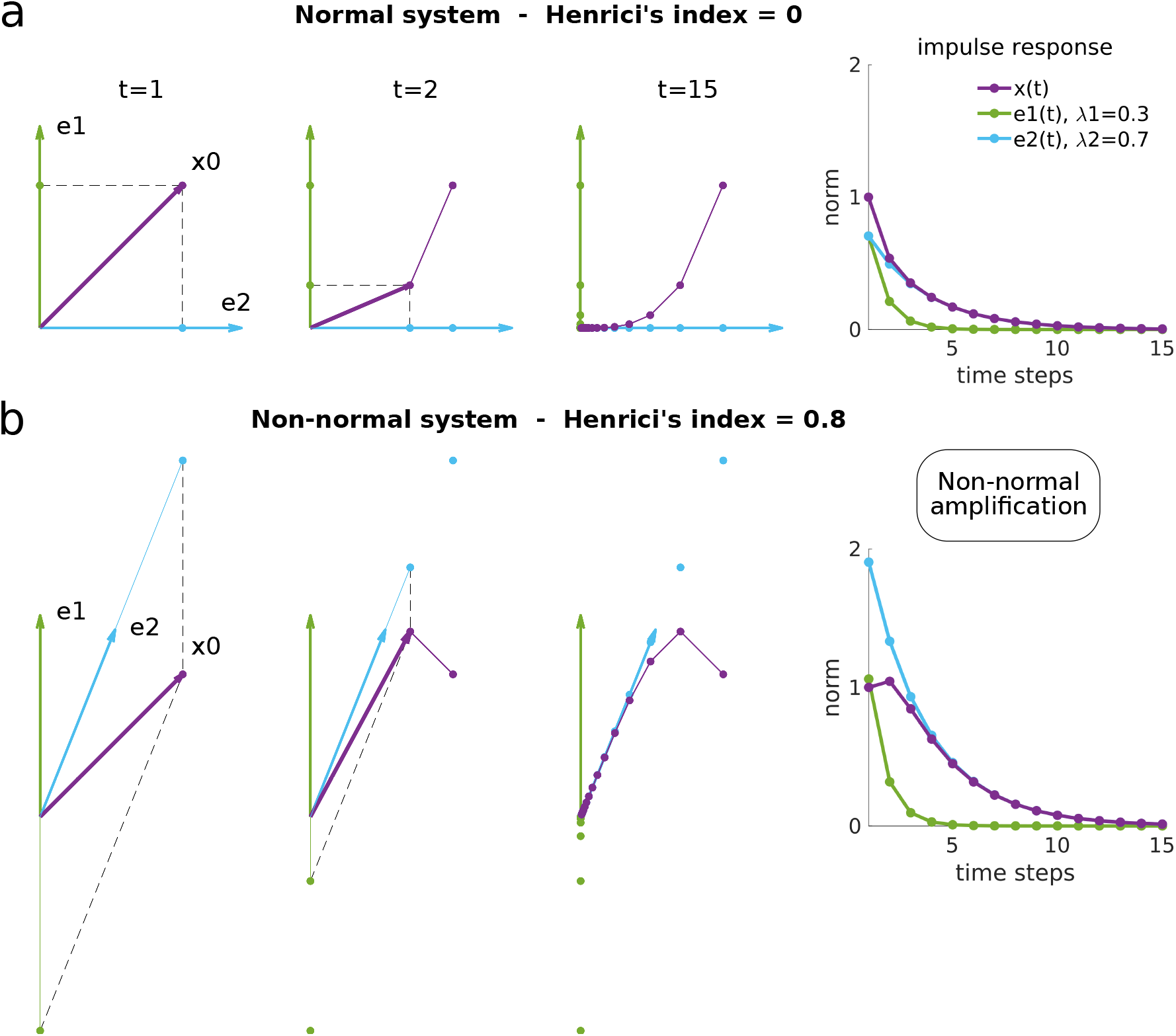
Non-normal transient amplification of inputs. Impulse response behavior of two 2D linear dynamical systems with normal (**a**) and highly non-normal (**b**) dynamics. The normal system, by construction, had two orthogonal right eigenvectors (*e*1, *e*2, left panels, green and blue arrows) and the non-normal system non-orthogonal ones (set to be closely aligned to each other). The left eigenvectors are not shown, since they are not relevant in this example. Both models were set to have identical eigenvalues, one being small (*λ*1 = 0.3, fast dynamics, in green) and the other large (*λ*2 = 0.7, slower but also decaying dynamics, in blue). The rightmost panels show the impulse response of both systems (the norm of the system state ||*x*(*t*)|| over time under a pulse input, or unit norm perturbation). The perturbation was given along a direction bisecting the plane spanned by the two right eigenvectors in the normal model (*x*0, left panels). For the normal system (**a**), the projection of the state onto the eigenvectors’ orthonormal basis at each time step (dashed lines, middle panels) gives the evolution along the dynamic modes (dots on *e*1 and *e*2 directions). For the non-normal system (**b**), the state cannot be decomposed using an orthogonal projection since the eigenvector basis is not orthogonal. Instead, the eigenvector components are given by the linear combination coefficients along two non-orthogonal basis vectors (here reconstructed using the parallelogram vector addition rule, dashed lines, left and middle panels). After the pulse, activity along the dynamic modes *e*1 and *e*2 decays exponentially for the normal system (right panel in **a**, in green and blue), and so does the state norm (in purple). For the non-normal system, the dynamic modes also decay exponentially (right panel in **b**, in green and blue). However, the state norm experiences a transient increase, followed by exponential decay (in purple), as a consequence of the non-orthogonal state-vector decomposition and the difference in decay rates of the two eigenmodes (left and middle panels in **b**). This phenomenon is known as non-normal transient amplification.

**Extended Data Figure 8.**
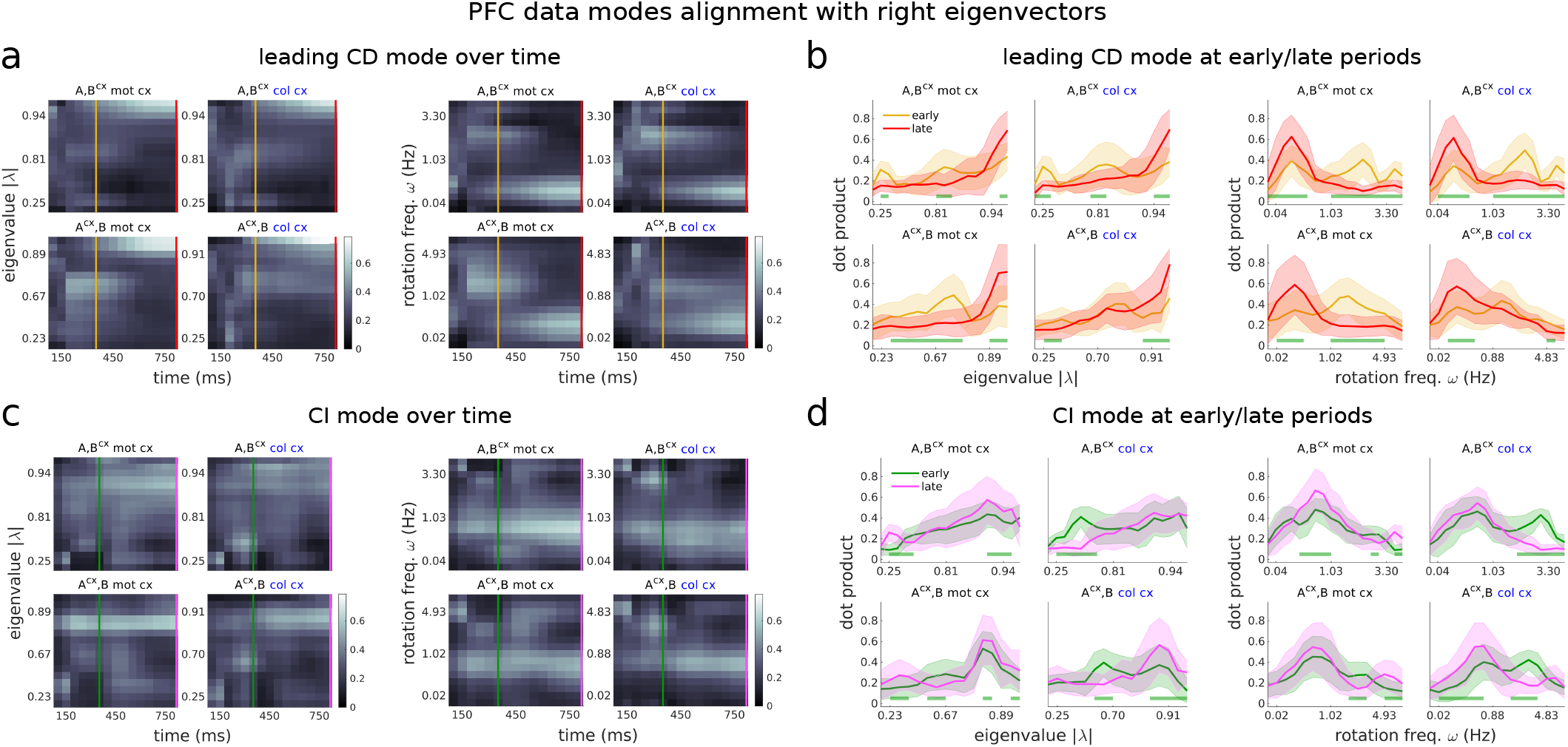
PFC integration phases for both LDS models across contexts. Same as in Fig. 7 but for both models and contexts. **a**,**b**, The two distinct integration phases are consistently observed across model classes (*A, B*^*cx*^ and *A*^*cx*^, *B*), contexts (mot and col cx) and model instantiations (**b**, the distribution of alignments across 100 randomly initialized models are fairly narrow, mean ± std). Note that for the *A*^*cx*^, *B* model in the color context, the early and late alignment distributions are not significantly different along the intermediate set of eigenmodes (at significance level p*<*0.001, **b**). However, the early distribution clearly peaks around the intermediate modes, which have relatively fast decaying dynamics (**b**, left) and fast rotations (**b**, right). The early and late alignment distributions are always significantly different along the largest modes, which have the slowest dynamics (**b**, left) and very small rotation frequencies (**b**, right). **c**,**d**, CI signals are integrated along a different set of slow modes than CD signals, consistently across models and contexts (the peaks of the CI distributions, **d**, lie in different ranges of slow eigenvalues than the early and late CD distribution peaks, **b**). The integration of CI signals do not clearly separate in two phases given that the alignments are largely steady across the trial (**c**). Indeed, the distribution of alignments early vs. late largely overlap (**d**, green and magenta distributions). Monkey A data.

**Extended Data Figure 9.**
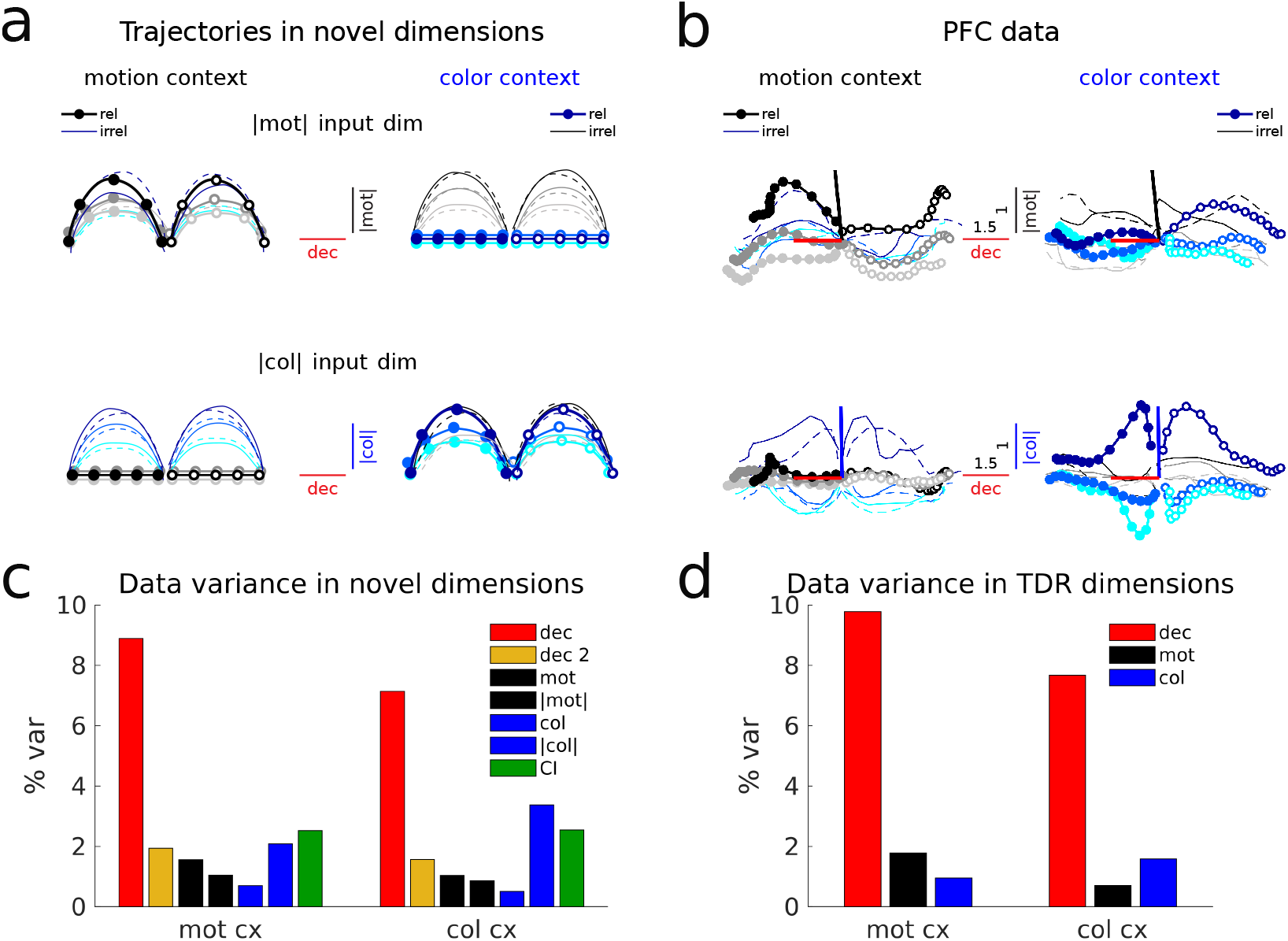
PFC data trajectories in coherence magnitude dimensions and data variance in the novel and TDR dimensions. **a**,**b**, Same as in Fig. 8, but for the LDS-inferred coherence magnitude dimensions (averaged across contexts and models). The condition independent (CI) variance has been subtracted-out from the trajectories to emphasise input-related variance. **c**, PFC data variance in the novel decision, secondary decision, motion coherence, motion coherence magnitude, color coherence, color coherence magnitude and condition independent (CI) dimensions. Note that the variance that is reflected in each dimension is the total variance, and not the isolated task-related variance. For instance, there is substantial CI variance in the input dimensions, in particular the color coherence magnitude ones, which we removed in panel **b** to emphasize input-related features. **d**, Variance in TDR decision, motion and color dimensions^1^. Monkey A data.

**Extended Data Figure 10.**
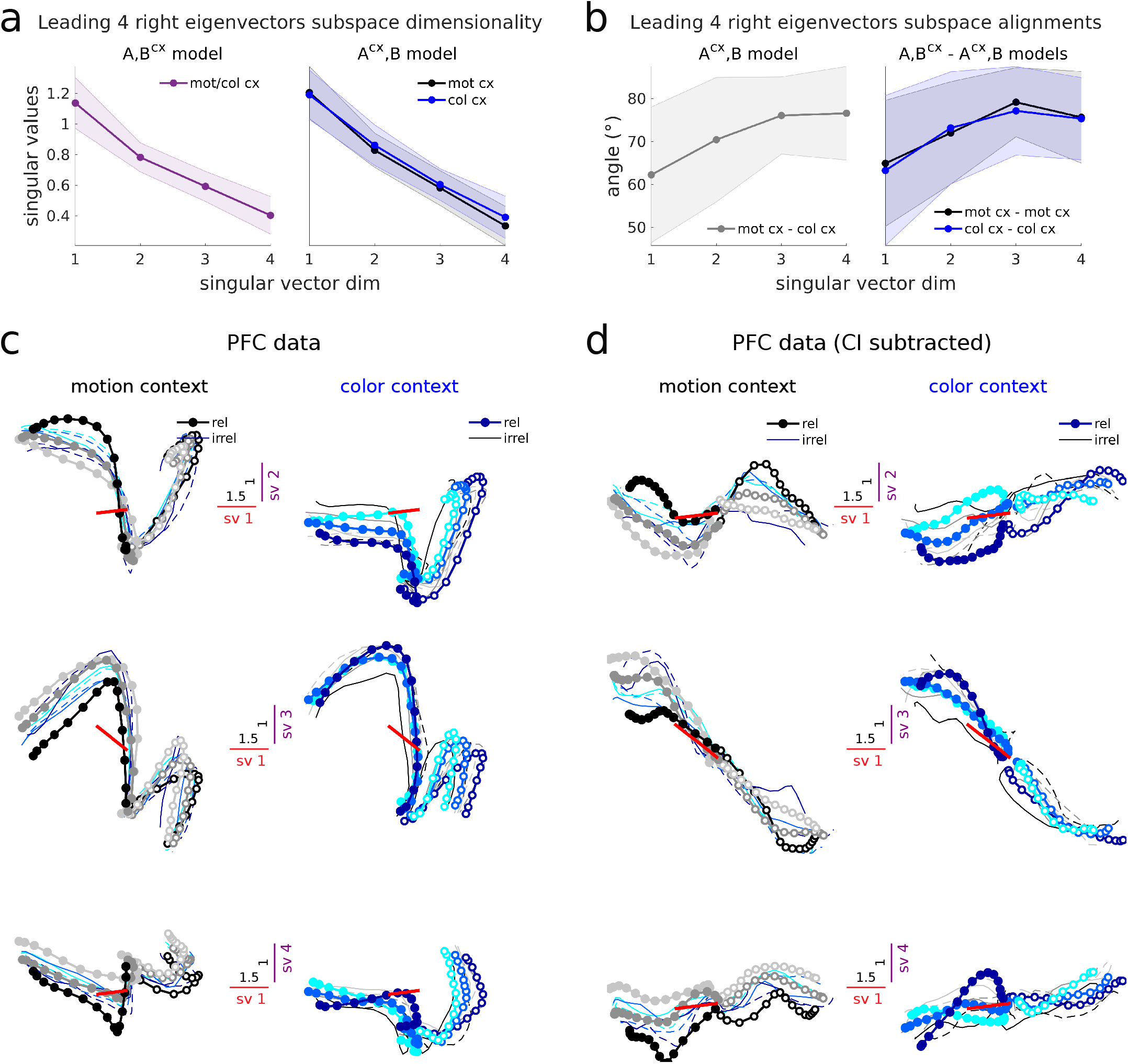
Choice signals in the slowest LDS subspace largely evolve along a single dimension across contexts. **a**, The four slowest dimensions inferred by the LDS models have very long time constants (|*λ*| *>* 0.9, time-constant *τ >* 475ms, average across 100 models, Fig. 5b), so they are expected to define a 4D subspace that contains highly persistent signals. However, the associated right eigenvectors are not orthogonal (Extended Data Fig. 6d), so these signals could occupy fewer than four dimensions. We measured the effective dimensionality of the subspace spanned by the four slowest right eigenvectors by taking the SVD of the matrix containing them. We found that the right eigenvectors effectively span four dimensions, since all singular values are well above zero. Mean ± std across 100 models. The dominant dimension (1st singular vector) is also the one most highly aligned across contexts in the *A*^*cx*^, *B* model and also across models (see next panel). **b**, Alignments of the slowest subspace dimensions found across contexts in the *A*^*cx*^, *B* model (left panel) and across models (right panel). Mean ± std across 100 models. The alignments are moderate to weak and expected by chance from a control distribution of random vector alignments in the low-d LDS subspaces (alignments are within the 5th and the 95th percentiles of the control distributions, Extended Data Fig. 6c). However, the largest alignments occur precisely for the first dimension. The moderate alignments may be explained by the effect of CI signals across contexts (see next panel) **c**, PFC data projected in the four singular vector (sv) dimensions of the slowest subspace (averaged across models and contexts, and then re-orthonormalized). Importantly, the first dimension captures decision information, but dimensions 2 to 4 mostly capture condition independent (CI) variance, and some contextual variance (in particular sv2, where the trajectories from each context are slightly shifted vertically). **d**, Same as **c** but after subtracting CI signals. Red bars show the alignment of the decision dimension found by TDR with respect to the four averaged singular vector dimensions (sv1: 36°, sv2: 86°, sv3: 65°, sv4: 86°). The first singular vector dimension highly aligns with the decision dimension (36°), and the third one moderately aligns to it (65°). These alignments are not expected by chance, considering a control distribution of random alignments in the ambient (high-d) space (alignments *<* 5th percentiles of the control distributions, Extended Data Fig. 6b). The 2D subspace spanned by these two singular vector dimensions (middle panels) contains trajectories mainly evolving along a single dimension, when CI signals are subtracted out (compare to middle panels in **c**). This dimension is common across contexts and strongly aligns to the decision axis found by Mante et al. (red bars). Similar results are obtained when projecting the data into the sv dimensions computed for each context and model independently. Monkey A data.

## Supplementary Information

1. **Supplementary Notes 1–3**
2. **Supplementary Figures 1–10**
3. **Supplementary Tables 1–5**

### 1 Supplementary Notes

#### 1.1 Additional LDS model-fitting controls

To rule out the possibility that the LDS models were learning dynamically complex external input signals **u**_*m,c*_(*t*) to capture the data, we re-trained all input parameters and constrained **u**_*m,c*_(*t*) to be constant in time **u**_*m,c*_. This resulted in a relatively small drop in performance (Extended Data Fig. 5a, for models with input dimensionalities higher than 2D). In particular, the performance of re-trained models with 3D constant inputs (dashed lines) dropped to the level of the 1D time-varying input models (filled lines). Newly fitted models with **u**_*m,c*_ constant in time, but where all parameters were optimized, performed nearly as well as the time-varying models (Extended Data Fig. 5b), and accurately captured the PFC trajectories (Extended Data Fig. 2g,h). Notably, in these two control model classes the optimal input dimensionality was consistently 3D, and the latent dimensionalty was also close to the time-varying input models’ dimensionality (*A*^*cx*^, *B* - *A, B*^*cx*^, dim = 16-22 for the time-varying models with inputs retrained to be time-constant; 15-16 for the newly optimized time-constant models and 16-18 for the original time-varying models). Most importantly, the time-constant input models could only rely on their recurrent dynamics to capture the temporal complexity of the PFC data. Therefore, the complexity of the PFC responses is well approximated by linear dynamics and is not necessarily inherited from the external inputs’ dynamics.

We also quantitatively assessed whether the time-constant input models were indeed able to capture complex temporal structure in the data. For this, we asked how well these models performed on time-shuffled data, which had no correlational structure across time. If there was no drop in performance, this would indicate that the time-constant input models were uniquely capturing time-unrelated structure, such as correlations across neurons and conditions. The performance of the best time-constant models (which had 3D inputs, Extended Data Fig. 5b) dropped substantially, being worse than the 1D input models and nearly as bad as the most contextually constrained *A, B* models (Extended Data Fig. 5c). This indicates that the simple, time-constant LDS models were indeed capturing the complex time-related structure present in the PFC data. However, these models still captured a substantial fraction of the time-shuffled data variance (24% on shuffled data vs 27% on the original data). This suggests that the LDS models might be in gran part capturing correlational structure across neurons and conditions, besides time-related variance (which makes sense, given that the LDS is a low-d model that extracts common structure across neurons, and across conditions via shared inputs). Indeed, surrogate data sets randomized across conditions, neurons and time were very poorly captured, even by the best time-varying LDS models (Extended Data Fig. 5d). These data sets were designed to preserve the primary statistics of the data, and thus these results also indicate that the LDS models were not merely capturing basic features of the data^13^.

#### 1.2 Understanding non-normal transient amplification

Transient amplification is a property of dynamical systems that have non-normal dynamics matrices^17,18^. These systems present non-trivial dynamical properties that are not predicted by their steady state behavior. In particular, such systems can transiently amplify inputs before decaying to a steady state. To illustrate how the transient amplification mechanism takes place, we built two simplified dynamical system models with identical specifications and only two dimensions, and made one normal (Extended Data Fig. 7a, degree of non-normality or Henrici’s index=0, see Methods) and the other highly non-normal (Extended Data Fig. 7b, Henrici’s index=0.8). In Extended Data Fig. 7a,b, we show the two right eigenvectors of the dynamics for each system (*e*1, *e*2, in green and blue), which define the “output” dimensions were activity evolves over time. In the normal system, by definition, these are orthogonal vectors. In the non-normal system, they need not be, as it is the case in this example. Two additional left eigenvectors exist, but in a normal system they are the same as the right eigenvectors. In a non-normal system the left eigenvectors are distinct from the right ones (see Methods), but these are not shown here since they are not crucial to understand this picture. We set one of the eigenvalues of the system to be small (*λ*1 = 0.3, fast dynamics) and the other large (*λ*2 = 0.7, slow dynamics), in both models. We then analysed the impulse response properties of both systems (for t=15 time steps, as in the PFC data). For this, we provided an input pulse of unit norm along a random direction *x*0 (in this case, for illustration purposes, a direction that bisected the plane spanned by the two right eigenvectors in the normal system, left panels in Extended Data Fig. 7a,b, purple arrow). We then looked at how each system processed such input by looking at the evolution of the system state over time *x*(*t*) and its norm (middle and right panels, purple lines). In the normal system, the input decayed exponentially towards its steady-state, at zero, as expected from a dynamical system with real eigenvalues smaller than one (rightmost panels). The evolution of the two modes of the dynamics was governed by the time-constant of the eigenvalues (right panel). The smallest eigenvalue mode decayed faster (in green) and the largest slower (in blue), with the slower eigenmode largely determining the evolution of the state at later stages (in purple). The evolution along the two dynamics modes could be obtained by projecting the state at each time step onto the orthonormal basis defined by the two right eigenvectors (left and middle panels). The non-normal system, however, had non-orthogonal eigenvectors. In particular, we purposely set the eigenvectors to be closely aligned to each other, to obtain strong non-normal dynamics. This meant that the system state could no longer be decomposed using an orthogonal projection. Instead, the state was constructed from a linear combination of a non-orthogonal eigenvector basis set (left and middle panels, by applying the parallelogram law of vector composition). We provided the same input perturbation as for the normal system. Despite the similarities in design (in particular, having the same eigenvalues), the impulse response properties of this system were very different. Most notably, right after the pulse, there was a transient increase in the state norm (middle and right panels). However, the state eventually decayed to zero. This is because the long-term behaviour of a non-normal system is still governed by its eigenvalues. The smallest eigenvalue mode decayed faster and the largest slower, and they did so exponentially, as in the normal system (rightmost panel). Yet, initially a transient amplification effect was observed. This happens precisely because of the difference in the decay rates of the modes, combined with the fact that the state is constructed with a non-orthogonal eigenvector basis (left and middle panels, note how the state vector, in purple, is reconstructed at each time step). The more aligned the eigenvectors are, and the stronger the difference between their eigenvalues, the larger the degree of non-normal amplification. Note that because of the non-trivial decomposition of the state along the non-orthogonal basis, the modes’ initial norm was very large (left and right panel, specially for the mode in blue). This resulted in the state experiencing a more sustained decay than in the normal system (right panel, compare blue and purple lines in both systems), i.e. the input pulse was transiently “persistent”. This precise behavior was found for the relevant motion and color inputs in the *A*^*cx*^, *B* model, but not the irrelevant inputs; or any inputs in the *A, B*^*cx*^ model (Fig. 6b).

#### 1.3 Biological implementation of the top-down input modulation mechanism: fixed input subspaces with context-dependent external inputs

The *A, B*^*cx*^ mechanism can be realized even in the case where the input dimensions *B* are fixed across contexts, which may be more intuitively aligned with biology—given that long-range projections are known to be anatomically stable. In this alternative view, the direction of input vectors *B***u** within the fixed input subspace can still be changed, provided that the external inputs are context-dependent **u**^*cx*^, and that the dimensionality of the input subspace is higher than 2D. This can be achieved by changing the external inputs’ strength in each context independently along each of the input dimensions (i.e. for 2D inputs, 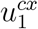 and 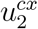). This effectively changes the input vector *B***u** coordinates within the input basis *B*. This change of coordinates can be used to rotate and stretch the inputs within the input subspace differently across contexts (i.e. 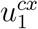 and 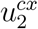 can be set to define any vector in the 2D input plane). The dimensionality of *B* (its columns) determines the number of inputs that originate from different subpopulations—within an area or across multiple areas. Changes in input direction could then be achieved through top-down modulation of the different subpopulations independently (i.e. modulation of subpopulation 1 would change 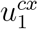 and of subpopulation 2 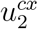, resulting in a change of *B***u** norm and direction).

### 2 Supplementary Figures

**Supplementary Figure 1.**
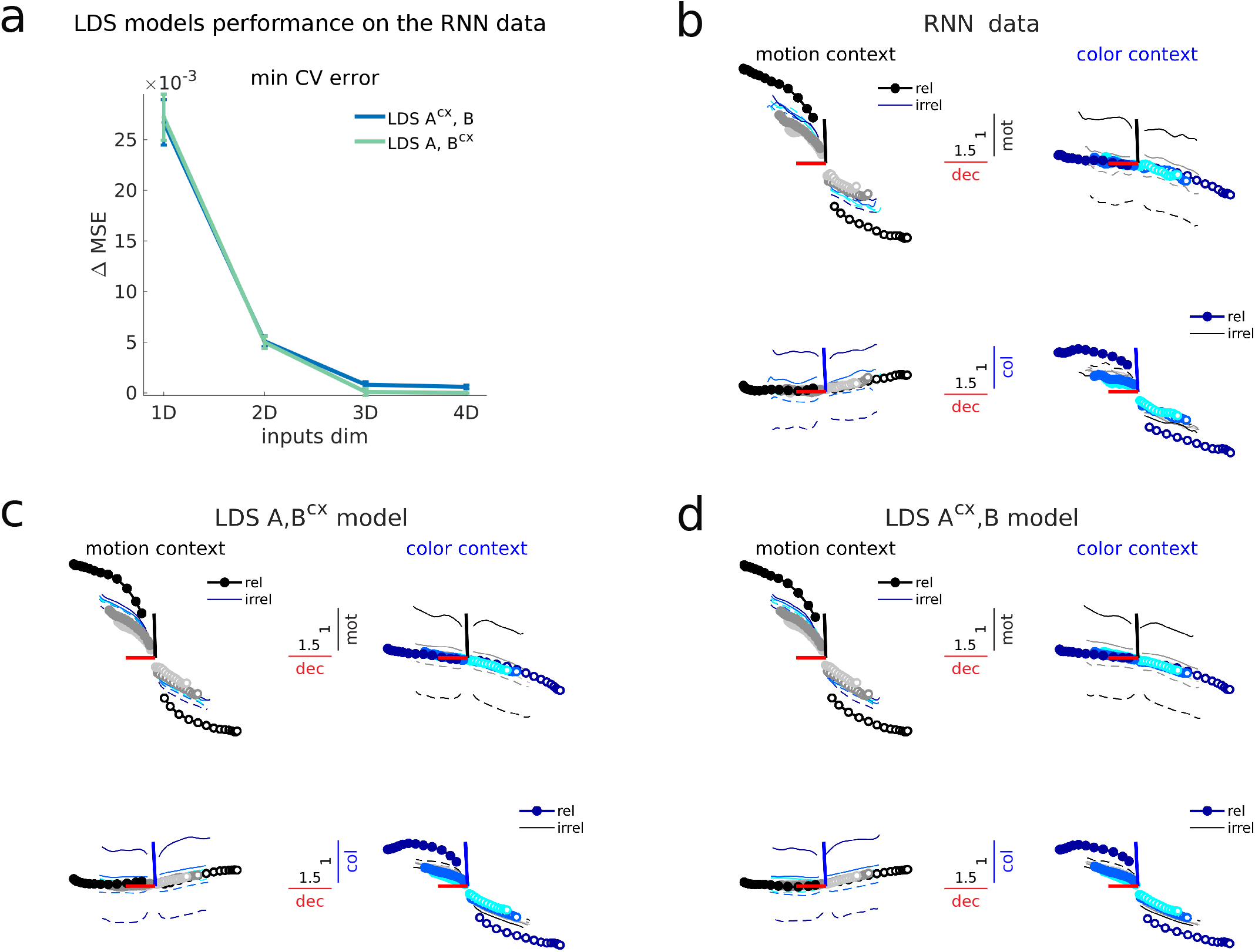
The LDS models accurately capture the RNN data. **a**, Same as Fig. 2a, but for the *A, B*^*cx*^ and *A*^*cx*^, *B* models fitted to the RNN data. In this case the best performing model was the *A, B*^*cx*^ model with 4D inputs. However, the performance of the *A, B*^*cx*^ model with 3D inputs was not significantly different (Supplementary Table 5). Therefore, in the RNN data the best performing models required at least 3D inputs, as for the PFC data (Fig. 2a). This was greater than the actual input dimensionality of the RNN model, which had 1D inputs (Fig. 1c), but see Supplementary Fig. 2 for an explanation. The two LDS model classes performed equally well (Supplementary Table 5), as in the PFC data (Fig. 2a), but the optimal latent dimensionality for the 3D *A, B*^*cx*^ model was 26 and for the *A*^*cx*^, *B* model 14 (Supplementary Table 5). The difference in dimensionality between the two model classes is larger than in the PFC data (18D vs. 16D, Supplementary Table 1). This suggests that the *A, B*^*cx*^ model struggles to capture the RNN data, since it needs more parameters. Note that the error for the two LDS models is close to zero (Supplementary Table 5, LOOCV MSE≈0, or ≈0% of variance missed), unlike for the PFC data from both monkeys (MSE≈0.73, or ≈73% of variance missed, Supplementary Tables 1 and 3). Thus, the LDS models are able to approximate near perfectly the dynamics of the non-linear RNN in each context. This further validates the adequacy of our linear dynamics model-fitting approach. **b-d**, Same as Fig. 3b-d but for the RNN data. Trajectories in the RNN task-related subspace are well captured by the both LDS models. Axes are defined by the RNN motion and color input vectors and the output vector (the decision readout) after training^1^. Note that the *A, B*^*cx*^ model can capture well the input variance along the RNN input dimensions, which by design are fixed across contexts (Fig. 1c). In spite of this, this LDS model must have changed the inputs to achieve contextual integration (given that it can only change *B*), accurately capturing the substantial amount of variance along the decision axis.

**Supplementary Figure 2.**
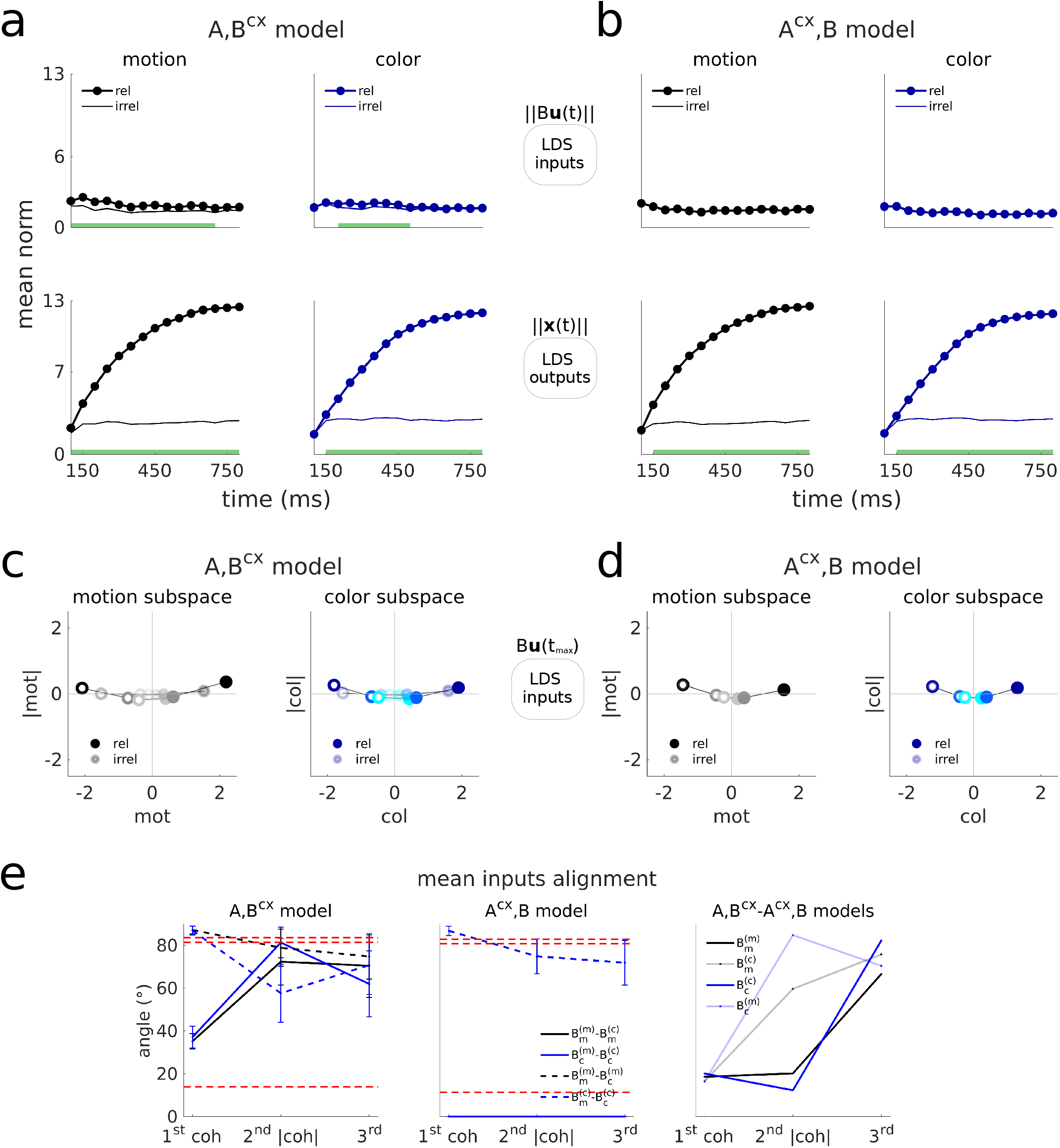
The LDS inputs inferred from the RNN data are largely constant over time and one-dimensional. **a-d**, Same as Fig. 4b-e but for the RNN data. **a**,**b**, Inputs are learned nearly flat in both the *A, B*^*cx*^ and *A*^*cx*^, *B* models, and almost identical across contexts in the *A, B*^*cx*^ model, which is consistent with the ground truth RNN inputs being time-constant and fixed across contexts (Methods). Yet, both models, including the *A, B*^*cx*^ model, strongly amplify the relevant inputs across contexts, but not the irrelevant ones. Note that the relevant inputs are much more strongly amplified than the irrelevant ones in the RNN, as expected from an optimal selective integration strategy^1^, but the same is not found in the PFC data, were the irrelevant outputs are prominent and persist throughout the trial (Fig. 4b,c). **c**,**d**, In the RNN the coherence representations are largely 1D since very little coherence magnitude modulation exists, unlike what is found for the PFC data (Fig. 4d,e). This is consistent with the ground truth RNN inputs being 1D and lacking a coherence magnitude component. **e**, Same as Extended Data Fig. 3c but for the RNN data. The alignments of the second inferred dimension (coherence magnitude) are not consistent across contexts in the *A, B*^*cx*^ model (first panel), and neither across models for the irrelevant inputs (third panel, transparent lines), unlike what is found for the PFC data (Extended Data Fig. 3c). Similar weak alignments are observed for the third input dimension, both in the RNN and the PFC data. This indicates that the coherence magnitude dimension in the RNN is not an invariant input feature. This is consistent with the ground truth RNN inputs being 1D and lacking a coherence magnitude component. Instead, the additional input dimensions might be learned to compensate for the linear approximation.

**Supplementary Figure 3.**
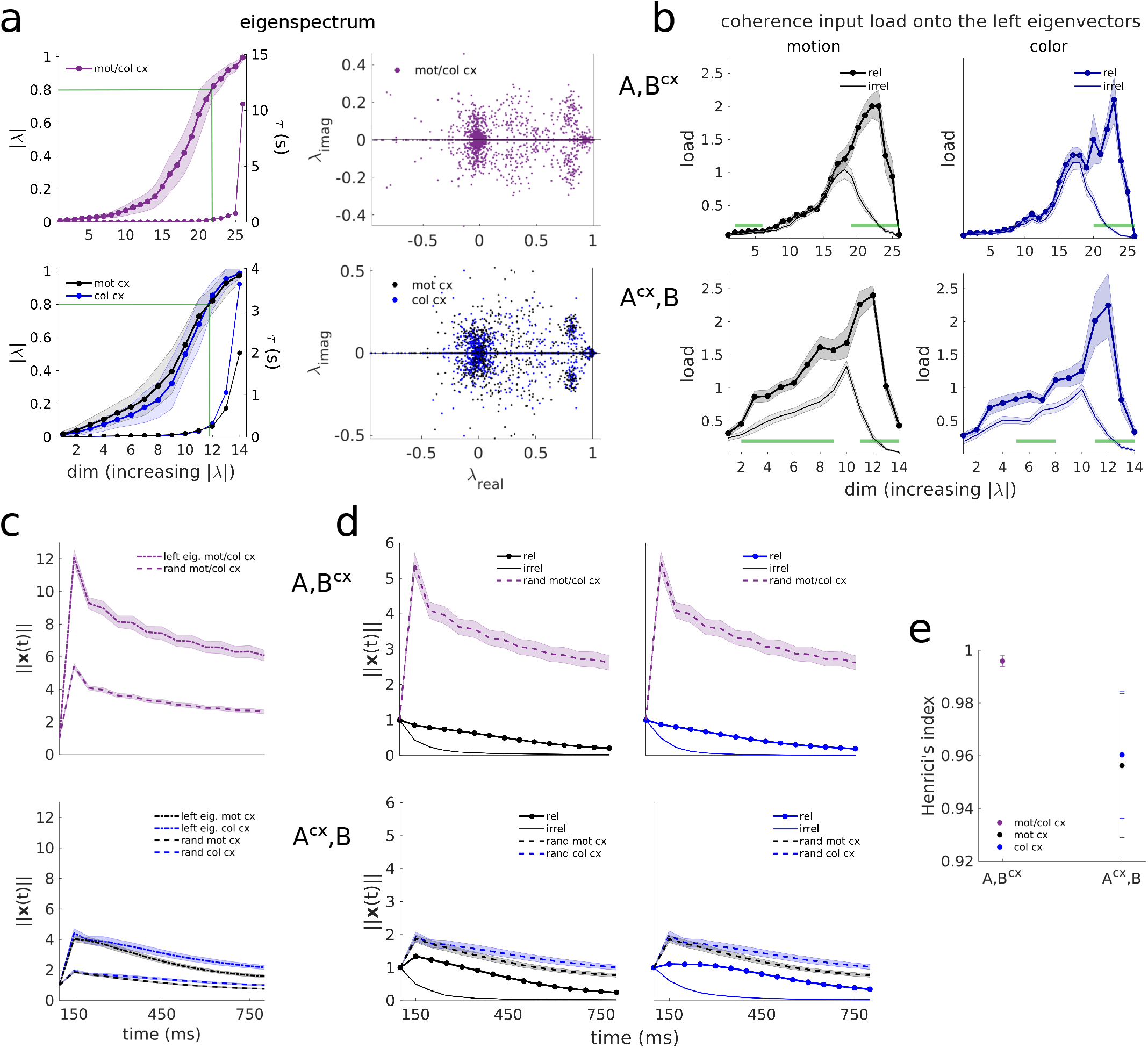
The LDS dynamics inferred from the RNN data are high-dimensional, mediate relevant input selection largely through slow modes, and implement transient amplification of relevant inputs in the two models—but through extreme non-normality for the *A, B*^*cx*^ model. **a**,**b**, Same as Fig. 5b,c and Extended Data Fig. 6a, but for the RNN data. **a**, The LDS models inferred from the RNN data are higher dimensional than expected for an idealized line attractor solution, with a single slow dimension and the rest of the dimensions fast decaying^16^. In particular, the LDS models learned several slow modes (|*λ*| *>* 0.8, green lines), as happened in the LDS data (Fig. 5b). However, the LDS models inferred from the RNN data had a smaller fraction of slow modes than the ones inferred from the PFC data (*A, B*^*cx*^, 20 ± 5%; *A*^*cx*^, *B*, 21 ± 7% mot cx/ 22 ± 5% col cx, mean±std across 100 models; vs. 35 − 55% in the PFC data). Furthermore, for the rest of the modes, most of them were very fast decaying (|*λ*| *<* 0.4, *τ <* 55ms), unlike what was found in the PFC data, where most of the modes had either intermediate (|*λ*| = 0.4 − 0.8, *τ* = 98 − 224ms) or slow eigenvalues (|*λ*| *>* 0.8). The largest eigenvalue was 1.00 ± 0.02 for the *A, B*^*cx*^ model and 0.98 ± 0.04 / 0.99 ± 0.04 (mot/col cx) for the *A*^*cx*^, *B* model. The second largest eigenvalue was 0.94 ± 0.03 for the *A, B*^*cx*^ model and 0.93 ± 0.08 / 0.95 ± 0.06 (mot/col cx) for the *A*^*cx*^, *B* model. The additional slow modes could have been learned to capture the curvature of the line attractor^1^, or alternatively, to capture CI and contextual variance, as found in the PFC data (Extended Data Fig. 10, see also Supplementary Fig. 5). Another possibility is that the higher dimensionality is a necessary feature of the mapping between low-rank linear RNNs and LDS models^70^ (note that the RNN dynamics was low-rank, and approximately linear in each context^1^). Similarly, this might also in part explain the dimensionality of the LDS models inferred from the PFC circuit. **b**, In both models, the coherence inputs inferred are most strongly loaded onto slow modes, rather than intermediate modes as in the PFC data (Fig. 5c). However, the inputs do not preferentially load onto the slowest modes (|*λ*| *>* 0.9, *τ >* 475ms), as we found in the data—but in the RNN the slowest modes carry CI signals, rather than decision-related CD ones (Supplementary Fig. 4) **c-e**, Same as Fig. 6a-c but for the RNN data. **c**, Left-eigenvector perturbations as well as random perturbations result in very strong amplification for the *A, B*^*cx*^ model, unlike what is found in the PFC data (Fig. 6a). Accordingly, the degree of non-normality for this model is extremely high (**e**), unlike what is found in the PFC data (Fig. 6c). On the contrary, the response behavior of the *A*^*cx*^, *B* fitted to the RNN data is similar to the behavior in the PFC data (Fig. 6a). **d**, Random perturbations are very strongly amplified by the *A, B*^*cx*^ model, but the inputs are not. This indicates a high degree of specificity in the model, which might imply fine-tuning. The *A*^*cx*^, *B* model, on the contrary, processes inputs in a similar way as found in the PFC data (Fig. 6b), with the difference that the relevant inputs in this case are transiently “amplified”, rather transiently “persistent”. This might be simply due to the fact that for the RNN data the inputs are learned much weaker than for the PFC data (Fig. 4c, Supplementary Fig. 2b), but the input pulse is provided with the same strength in both cases (unit norm). Also, the *A*^*cx*^, *B* model is slightly more non-normal for the RNN data (**e**) than the PFC data (Fig. 6c). This relatively high level of non-normality might also indicate some level of fine-tuning due to model mismatch—which might as well apply to the models from the PFC data, but to a lesser degree.

**Supplementary Figure 4.**
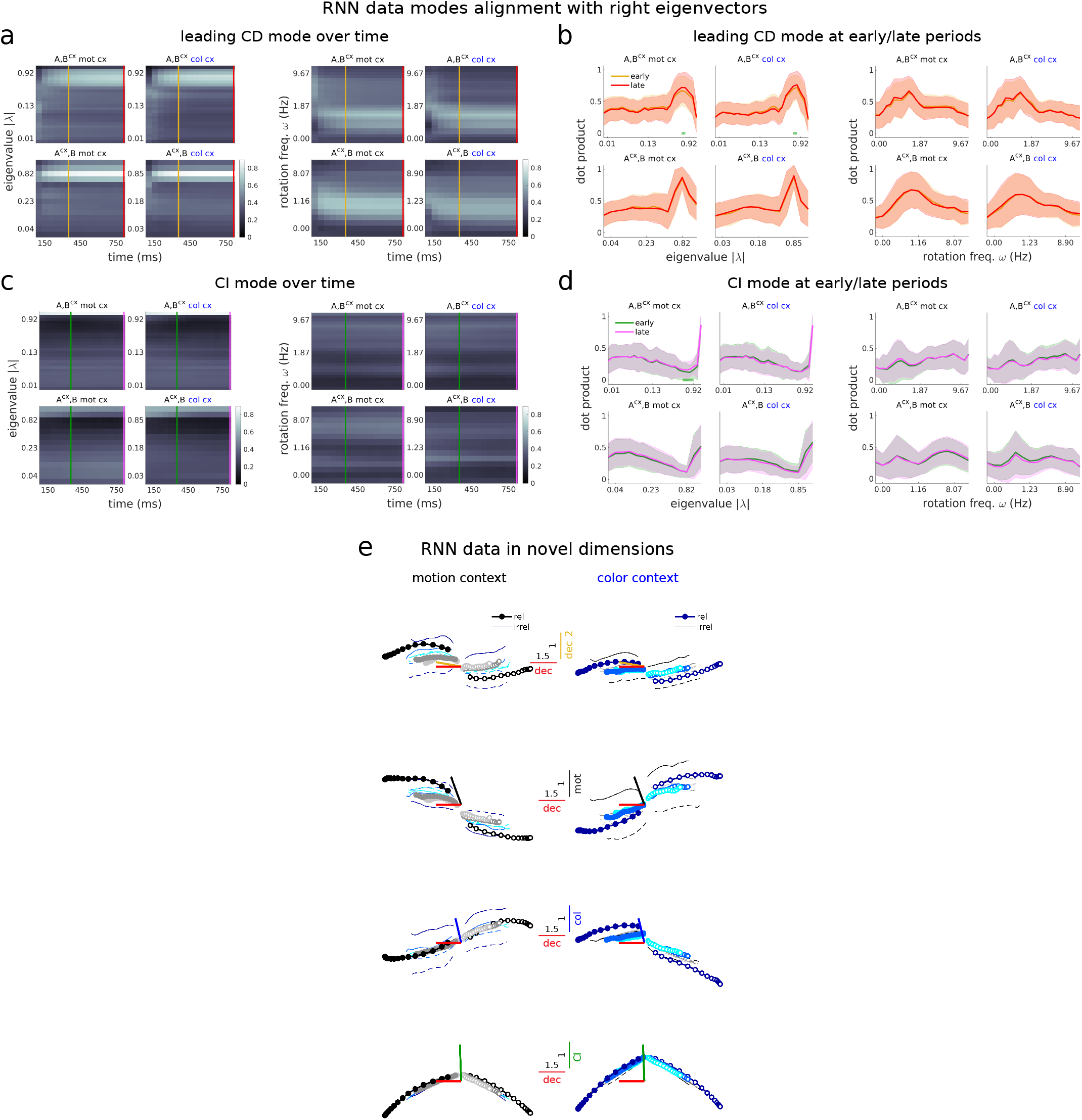
The RNN integration process did not separate in two phases. **a-d**, Same as Fig. 7, Extended Data Fig. 8, but for the RNN data. **a**,**b**, The largest CD mode for the RNN data projects to slow modes throughout the whole trial, unlike what is found in monkey A’s PFC data, where the CD vector projects to relatively fast decaying modes first and to the slowest modes later in the trial. Thus, the CD projection pattern in the RNN does not change over the course of the trial. Indeed, the distributions of projection early vs. late in the trial are practically indistinguishable (absence of green bars in **b**, Wilcoxon rank-sum test, p*<*0.05). The CD vector aligns to slow modes (|*λ*| *>* 0.8), but not preferentially to the slowest modes (|*λ*| *>* 0.9), unlike what is found in monkey A’s PFC data. The dynamics was also largely non-rotational (≈1Hz), and not significantly different early vs. late in the trial (right panels in **b**). **c**,**d**, The largest CI mode for the RNN data projects most strongly to the slowest modes, unlike what is found for monkey A’s PFC data, where the CI vector targets slow modes, but not the slowest. **e**, Same as Fig. 8b, but fore the RNN data. Top, the dimension inferred early in the trial highly aligns to the decision axis (small angle between red and yellow bars). This is consistent with the fact that early in the trial the RNN CD vector already projects to slow modes (**a**,**b**), and not to a different set of dimensions (the relatively fast decaying modes), as is found in monkey A’s PFC data, which define a secondary decision dimension (Fig. 8b). Middle panels, trajectories in the LDS motion and color input coherence dimensions found from the RNN data fits. The coherence input vectors found by the LDS models are only moderately aligned with the ground truth RNN input vectors (*A, B*^*cx*^: mot = 45°, col = 55°, for mean coherence input dimensions across 100 models and across contexts; *A*^*cx*^, *B*, mot = 43°, col = 54°, for mean across 100 models), but higher than expected by chance (Extended Data Fig. 6b). However, the RNN trajectories along the LDS coherence input dimensions (middle panels) are qualitatively similar to the trajectories along the ground truth RNN input vectors, and separate coherence information similarly well (Supplementary Fig. 1b).

**Supplementary Figure 5.**
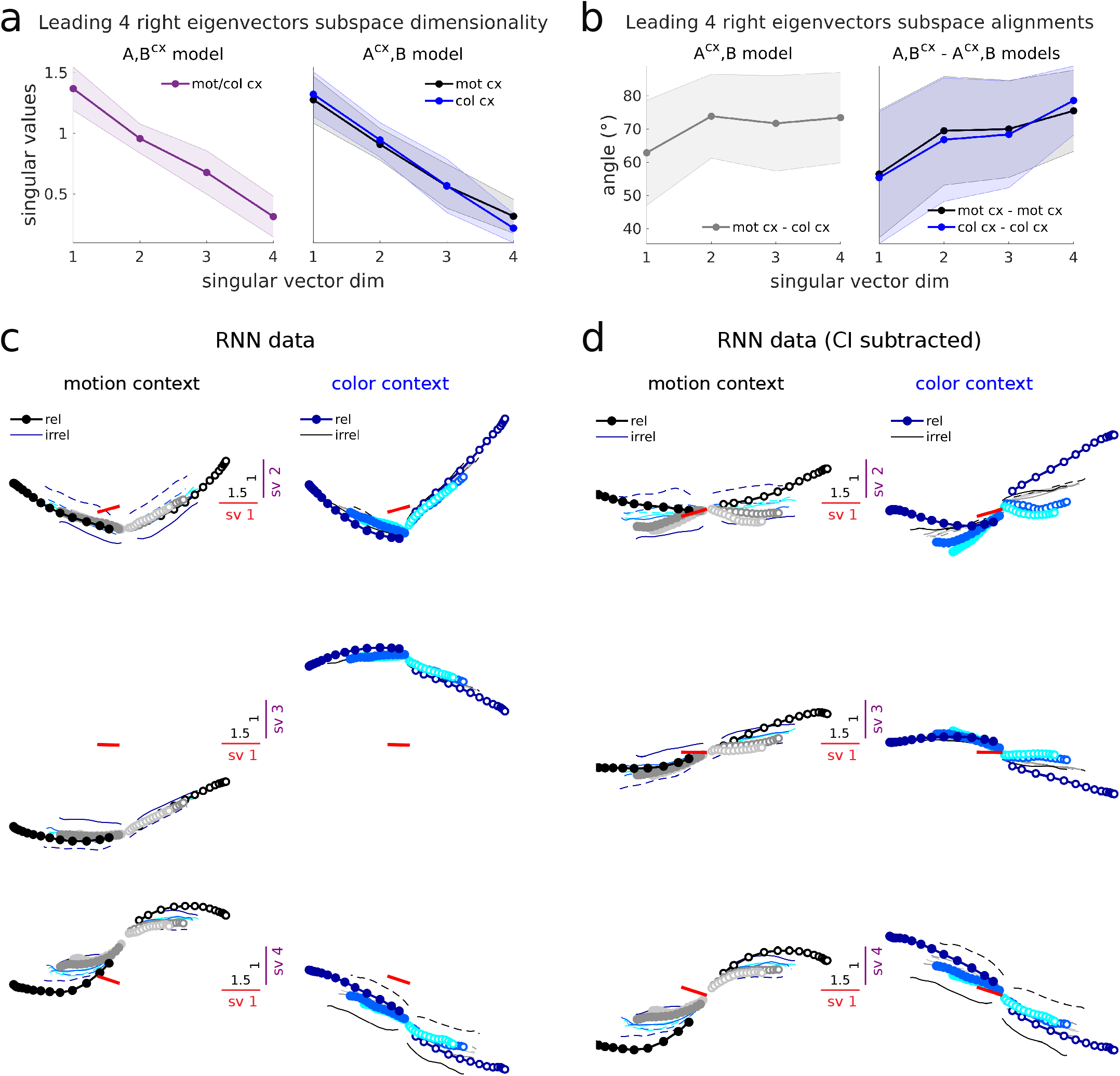
Choice signals in the slowest LDS subspace largely evolve along a single dimension across contexts in the RNN data. Same as Extended Data Fig. 10, but for the RNN data. **a**,**b**, The right eigenvectors effectively span four dimensions, since the singular values are not close to zero, as is found in the PFC data. Furthermore, the first first singular vector dimension is also the most aligned across contexts in the *A, B*^*cx*^ model and across models. **c**,**d**, The first singular vector dimension (sv) captures decision information. However, sv dimensions 2 to 4 mostly capture condition independent (CI) variance and contextual variance. Indeed, only the first dimension aligns well with the RNN decision dimension (red bars), which is defined by the RNN output vector or readout after training^1^ (sv1: 61°, sv2: 85°, sv3: 90°, sv4: 84°). Thus, the RNN trajectories mainly evolve along a single dimension in this 4D subspace, which aligns with the decision (or output) axis of the network. The other dimensions are used to capture the curvature of the approximate line attractor (sv4), CI features (sv2), and contextual separation (sv3, sv4).

**Supplementary Figure 6.**
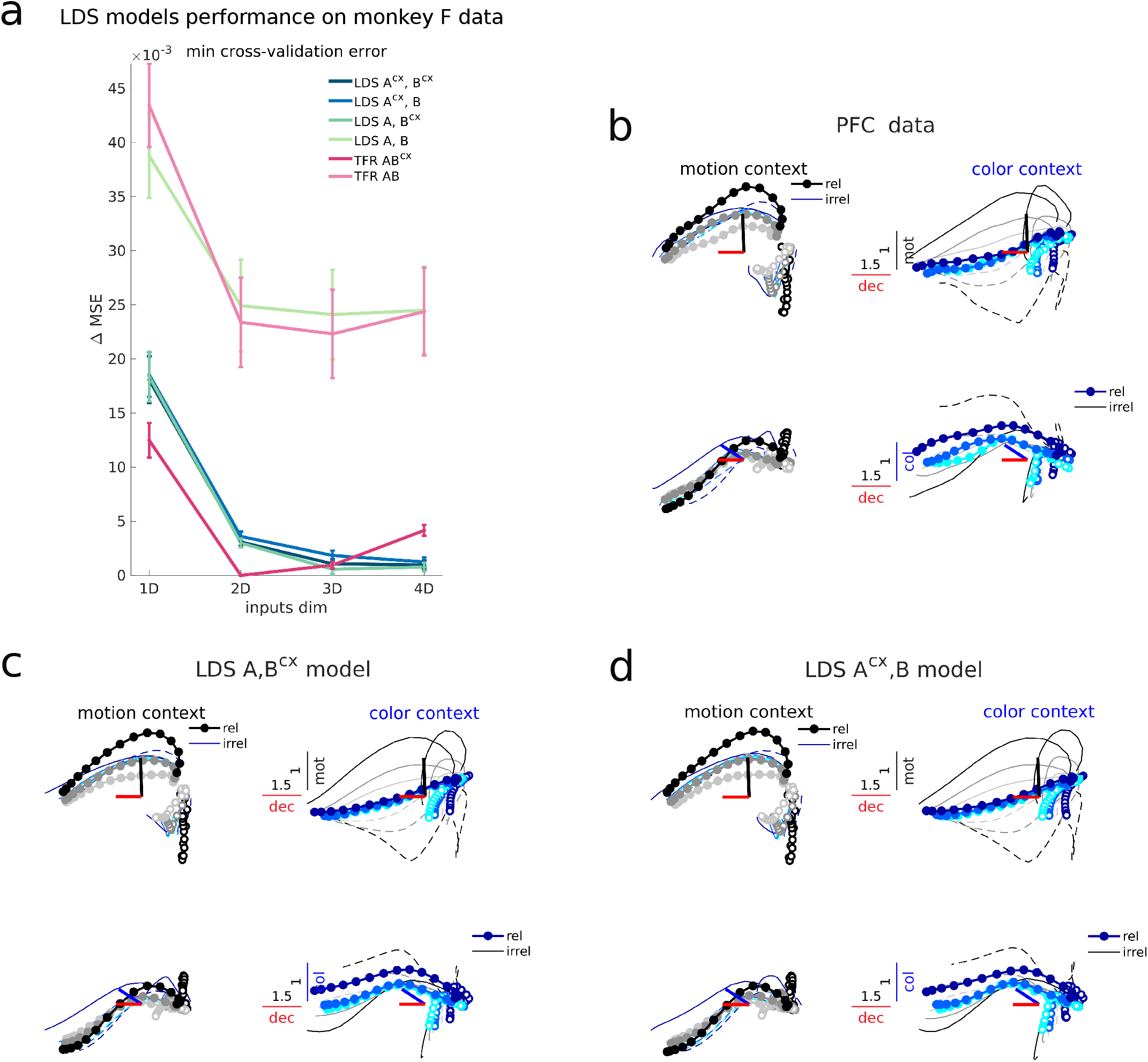
The LDS models accurately capture monkey F’s data, perform comparably to TFR and require multi-dimensional inputs. Same as Supplementary Fig. 1, but for monkey F data. **a**, The pattern of errors across models and input dimensions closely follows the one obtained for monkey A (Fig. 2a). The best performing model is also the *A, B*^*cx*^ model with 3D inputs. The *A*^*cx*^, *B* model with 3D inputs performs similarly well (Supplementary Table 3). **b-d** Same as Fig. 3b-d but for monkey F data. The LDS models accurately capture the trajectories in the contextually-stable task-relevant subspace. Note the presence of strong condition independent (CI) signals along the choice axis^1^, which are also captured by the models.

**Supplementary Figure 7.**
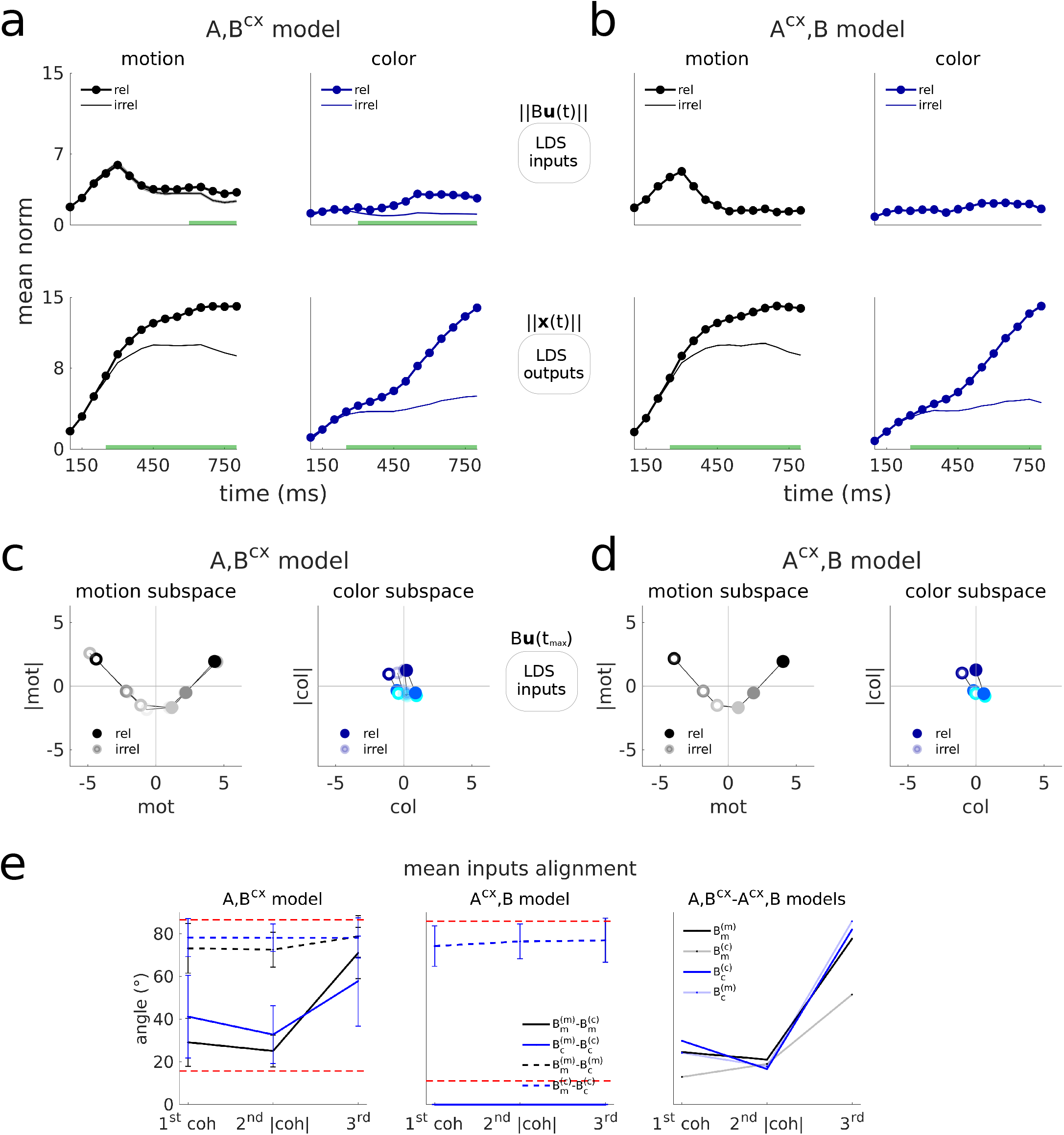
Monkey F’s LDS inputs are largely stable across contexts and span curved manifolds, with motion inputs being strongly integrated in both contexts. Same as Supplementary Fig. 2, but for monkey F data. **a**,**b**, The motion inputs from both models are similar to the inputs inferred for monkey A, in that these are transient in both the *A, B*^*cx*^ and *A*^*cx*^, *B* models, but more sustained in the *A, B*^*cx*^ model (top left panels). However, the *A, B*^*cx*^ model infers motion inputs that are nearly identical in strength across contexts (**a**, top left), unlike what is found for monkey A (Fig. 4b). Accordingly, the motion inputs are strongly integrated in both contexts, with the relevant outputs being only slightly stronger than the irrelevant ones (**a**, bottom left). Another difference is that the inferred color inputs are very weak, although these increase slightly towards the end of the trial when relevant (**a**, top right). Yet, the *A, B*^*cx*^ model selectively and strongly integrates the color inputs (**a**, bottom right). Both models generate identical outputs (**a**,**b**, bottom). The color outputs increase more sharply in the middle of the trial, and continue growing until the end of the trial. On the contrary, motion outputs saturate towards the end of the trial (bottom left). This saturation is also observed in monkey A, for both the motion and the color outputs (Fig. 4b,c). Thus, color inputs might be integrated later in the trial in monkey F. An alternative explanation could be that color signals arriving into monkey F’s PFC circuit are already integration signals, and our LDS models learn this particular input-ouput solution due to the fact that they incorporate an input penalty encouraging weak inputs (Methods). In line with this interpretation, we found that the LDS coherence input dimension is highly aligned with the decision dimension (Supplementary Fig. 9e). **c**,**d**, The motion coherence representations inferred by both models are strongly curved in this monkey, unlike for monkey A. The color inputs are weak along both the coherence and the coherence magnitude dimensions. **e**, The pattern of alignments between the different input dimensions across models and contexts is consistent across monkeys. The inferred coherence and coherence magnitude dimensions for both motion and color are largely stable across contexts in the *A, B*^*cx*^ model, but not the third dimensions (filled lines). One difference is that the motion coherence magnitude dimension is highly aligned across contexts in the *A, B*^*cx*^ model in this monkey, but less so in monkey A (Extended Data Fig. 3c). This is consistent with the motion coherence representations having a stronger curvature in monkey F than in monkey A (**c**,**d** vs. Fig. 4d,e).

**Supplementary Figure 8.**
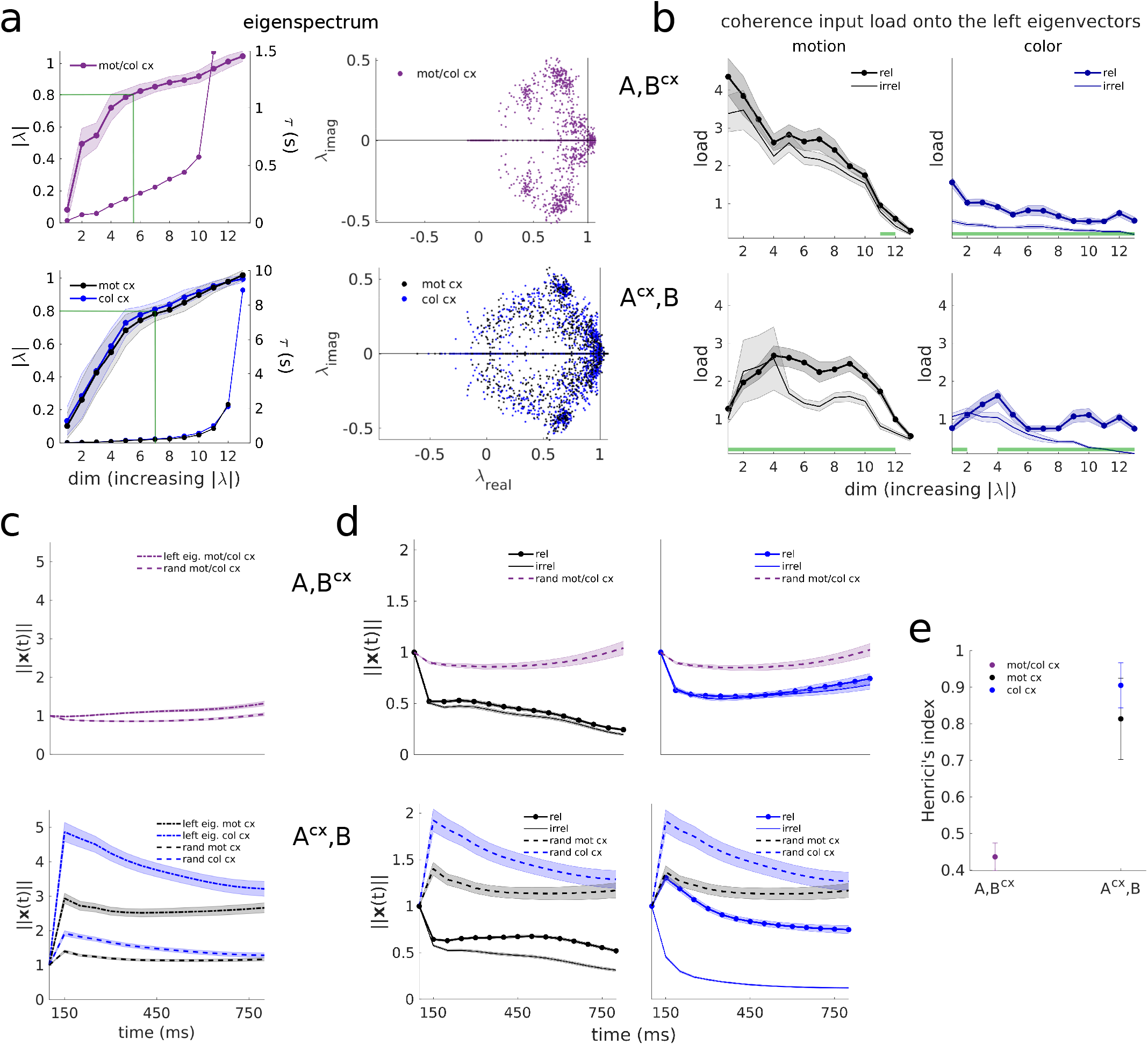
Monkey F’s linear dynamics are high-dimensional, perform context-dependent input selection through several intermediate and slow modes and implement transient amplification of relevant inputs only in the *A*^*cx*^, *B* model. Same as Supplementary Fig. 3, but for monkey F data. **a**, The inferred eigenspectrum from both models presents multiple slow modes, as in monkey A. The *A, B*^*cx*^ model had also a larger fraction of slow modes than the *A*^*cx*^, *B* model (63 ± 7% vs. 46 ± 8% mot cx/ 53 ± 8% col cx, mean±std across 100 models). Note that the last two eigenvalues of the *A, B*^*cx*^ model are slightly larger than 1 (unstable), so the time constant is not shown. Same for the last eigenvalue of the *A*^*cx*^, *B* model in the motion context. **b**, The coherence inputs are not preferentially loaded onto the slowest modes, but rather, intermediate and fast modes, as in monkey A’s data. However, the *A, B*^*cx*^ model loads the motion inputs selectively only onto the second and third slowest modes (green bar). **c** In the *A, B*^*cx*^ model the average impulse response across random perturbations grows over time, indicating the presence of unstable dynamics (top), which was not found in monkey A. On the contrary, the *A*^*cx*^, *B* model exhibited similar transient responses as monkey A (bottom). **d** The instability effect is also observed for perturbations along the color input dimension (top right panel). This feature might help integrate color input signals selectively, given that color inputs are very weak (Supplementary Fig. 7a). The *A*^*cx*^, *B* model response to motion and color pulses is broadly similar to that of monkey A’s. However, the responses were more amplified for the relevant color inputs in this monkey, and the relevant motion inputs exhibited transient amplification effects later in the trial, rather than immediately after the pulse. **e**, The degree of non-normality of the LDS models is similar across monkeys (Fig. 6c).

**Supplementary Figure 9.**
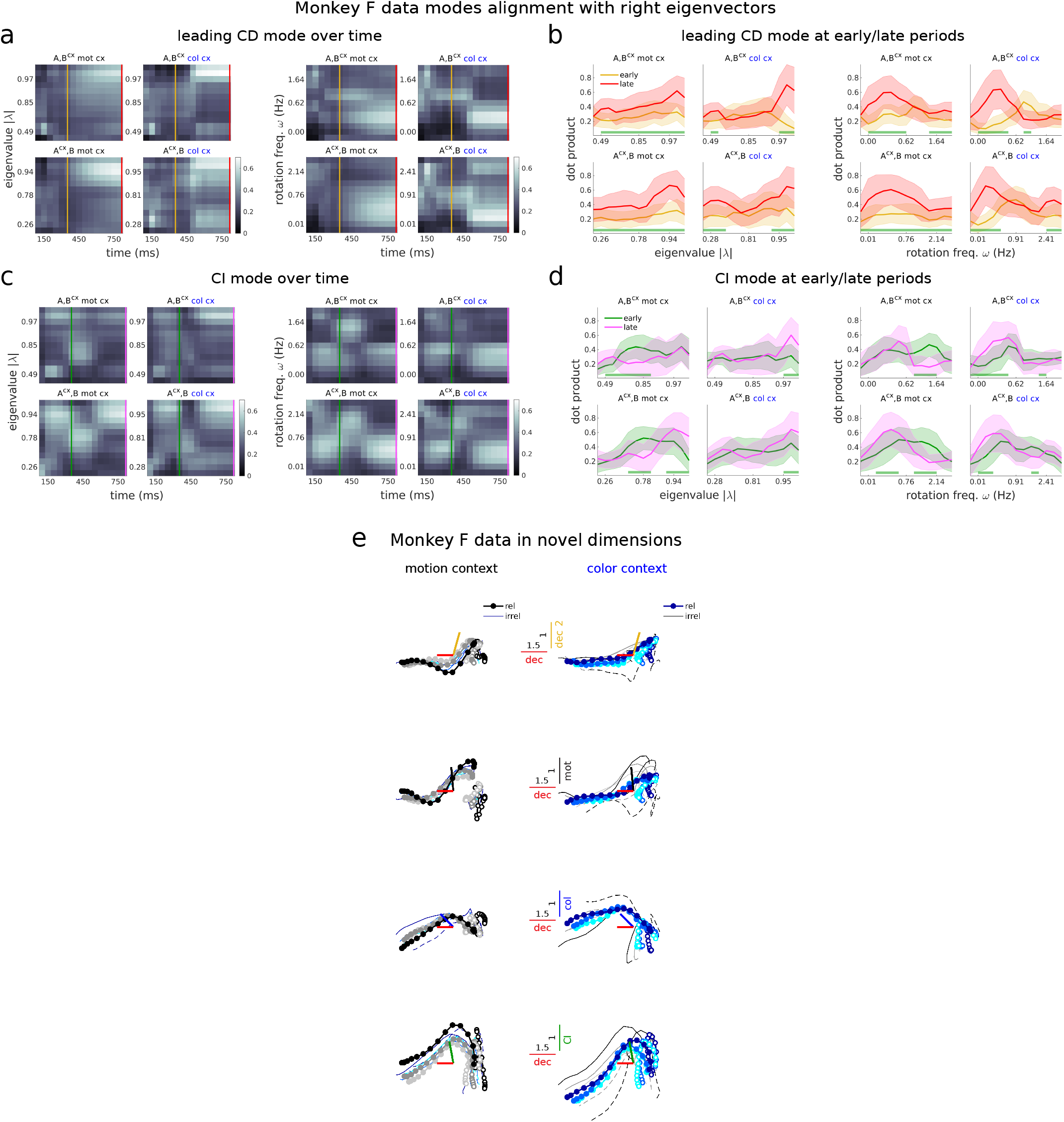
Monkey F’s integration process did not clearly separate in two phases, and CI signals were integrated along with choice signals. Same as Supplementary Fig. 4, but for monkey F data. **a**,**b**, Monkey F did not present a clear separation of the integration process into different phases, unlike monkey A. In particular, early in the trial the leading CD variance vector did not preferentially project onto relatively fast decaying modes (**b**, left panels, yellow lines, the pattern of projections is nearly flat), unlike for monkey A (Fig. 7b, Extended Data Fig. 8b, left panels, yellow lines, projections pick at intermediate modes). However, the pattern of early vs. late projections was similar to that of monkey A when splitting the modes by their rotation frequency (**b**, right panels, yellow lines in the color context, and in the motion context for the *A, B*^*cx*^ model, pick on intermediate rotation frequencies, as for monkey A, Fig. 7b, Extended Data Fig. 8b, right panels). The modes targeted during the late phase of the trial (red line projections) are consistent across monkeys both in time constant and rotation frequency. This late projections significantly differ from the early projections (green bars in **b**, Wilcoxon rank-sum test, p*<*0.001). **c**,**d**, The CI data vector for monkey F projects most strongly onto the slowest modes, in particular at the end of the trial (**d**, pink lines). This indicates that CI signals are integrated along the same dimensions as the CD signals, i.e., the dimensions that integrate coherence inputs (compare with red lines in **b**), unlike what is found for monkey A (Fig. 7c,d, Extended Data Fig. 8c,d, left panels, pink lines pick onto slow modes, but not the slowest). This is consistent with the fact that CI signals are strongly present along the decision axis^1^. The block-like structure found in the pattern of projections in panels **a**,**c** comes from the fact that the variance along the first singular vector dimensions is close to the variance along the next dominant singular value dimensions (i.e. that the covariance structure of the data is largely spherical), which leads to switches in the estimation of the dominant variance direction. **e**, The early-phase dimension or secondary decision dimension (yellow) does not capture much structure of the trajectories in monkey F (top). The late-phase dimension (red) is strongly aligned with the TDR decision axis (18°). The LDS coherence input dimensions were well aligned with the TDR dimensions in the *A*^*cx*^, *B* model, specially for color, but not in the *A, B*^*cx*^ model (*A, B*^*cx*^: mot = 54°, col = 71°, for mean coherence input dimensions across 100 models and across contexts; *A*^*cx*^, *B*, mot = 44°, col = 31°, for mean across 100 models). The averaged coherence input dimension across contexts and models highly aligned with the decision dimension (angle between blue and red bars in middle panels). This alignments were higher than expected by chance (Extended Data Fig. 6b).

**Supplementary Figure 10.**
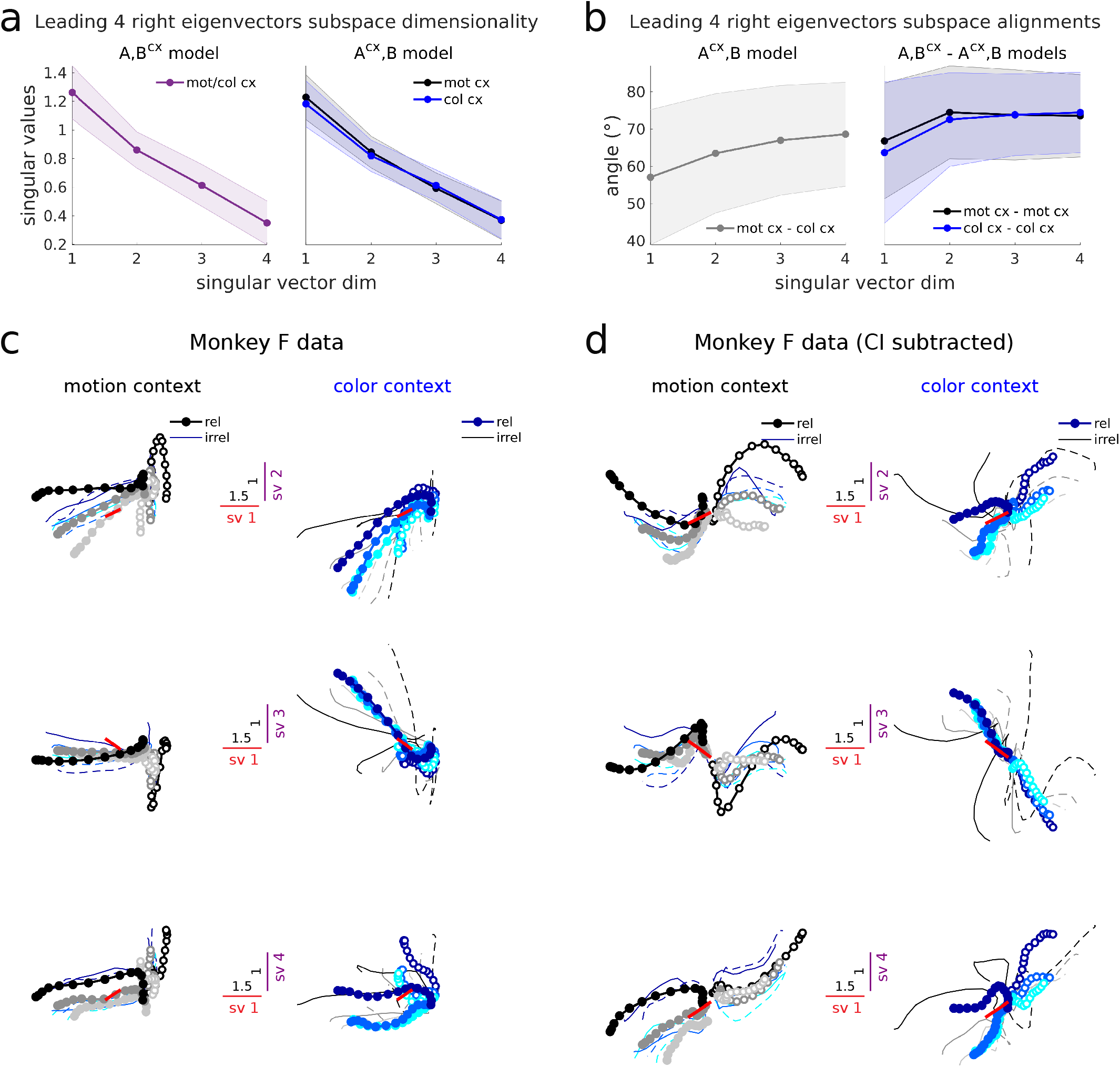
Choice signals in the slowest LDS subspace largely evolve along a single dimension across contexts in monkey F’s data. Same as Supplementary Fig. 5, but for monkey F data. **a**,**b**, The right eigenvectors effectively span four dimensions, since the singular values are not close to zero, as found in the other monkey. Furthermore, the first singular vector dimension is also the most aligned across contexts in the *A, B*^*cx*^ model and across models (Extended Data Fig. 10a,b). **c**,**d**, The first singular vector dimension (sv) captures decision information. However, sv dimensions 2 to 4 mostly capture condition independent (CI) variance and contextual variance. Indeed, only the first dimension aligns well with the TDR decision dimension (red bars) (sv1: 58°, sv2: 80°, sv3: 75°, sv4: 76°).

### 3 Supplementary Tables

**Supplementary Table 1.** Monkey A data minimum LOOCV MSE ± sem across k folds (36 conditions) and corresponding latent dimensionality for which it is achieved, for LDS models with different input dimensionalities and contextual constraints. Highlighted in black is the model that achieved the minimum MSE. It cannot be appreciated in the 4th column (min MSE) due to rounding error, but can be seen in the last column (min Δ(MSE ± sem)), where performance is given relative to the best performing model (TFR 2D *AB*^*cx*^ model, see Supplementary Table 2), so the differences across models, albeit small, can be revealed. This quantity is the one reported in Fig. 2a.

| Input dim | Contextual constraints | Latent dim | min MSE | min $\Delta\text{MSE} (\times 10^{-3})$ |
| --- | --- | --- | --- | --- |
| 1D | $A^{cx}, B^{cx}$ | 13 | $0.74 \pm 0.02$ | $13 \pm 1$ |
| | $A^{cx}, B$ | 15 | $0.74 \pm 0.02$ | $13 \pm 1$ |
| | $A, B^{cx}$ | 15 | $0.74 \pm 0.02$ | $12 \pm 1$ |
| | $A, B$ | 16 | $0.78 \pm 0.02$ | $48 \pm 6$ |
| 2D | $A^{cx}, B^{cx}$ | 15 | $0.73 \pm 0.02$ | $2.7 \pm 0.4$ |
| | $A^{cx}, B$ | 15 | $0.73 \pm 0.02$ | $2.7 \pm 0.4$ |
| | $A, B^{cx}$ | 16 | $0.73 \pm 0.02$ | $1.7 \pm 0.6$ |
| | $A, B$ | 17 | $0.77 \pm 0.02$ | $38 \pm 6$ |
| 3D | $A^{cx}, B^{cx}$ | 14 | $0.73 \pm 0.02$ | $1.3 \pm 0.4$ |
| | $A^{cx}, B$ | 16 | $0.73 \pm 0.02$ | $1.3 \pm 0.3$ |
|  | **$A, B^{cx}$** | **18** | **$0.73 \pm 0.02$** | **$0.6 \pm 0.5$** |
| | $A, B$ | 17 | $0.77 \pm 0.02$ | $38 \pm 6$ |
| 4D | $A^{cx}, B^{cx}$ | 14 | $0.73 \pm 0.02$ | $1.9 \pm 0.4$ |
| | $A^{cx}, B$ | 14 | $0.73 \pm 0.02$ | $1.8 \pm 0.4$ |
| | $A, B^{cx}$ | 16 | $0.73 \pm 0.02$ | $1.0 \pm 0.5$ |
| | $A, B$ | 17 | $0.77 \pm 0.02$ | $39 \pm 6$ |

**Supplementary Table 2.** Monkey A data minimum LOOCV MSE ± sem, min Δ(MSE ± sem) and corresponding latent dimensionality for which it is achieved, for TFR models with different input dimensionalities and contextual constraints. Same conventions as in Supplementary Table 1. Performance in the last column is given relative to the best performing model (TFR 2D *AB*^*cx*^ model).

| Input dim | Contextual constraints | Latent dim | min MSE | min ΔMSE ($\times 10^{-3}$) |
| --- | --- | --- | --- | --- |
| 1D | $AB^{cx}$ | 13 | $0.74 \pm 0.02$ | $9 \pm 1$ |
| | $AB$ | 14 | $0.78 \pm 0.02$ | $51 \pm 7$ |
| 2D | **$AB^{cx}$** | **14** | **$0.73 \pm 0.02$** | **$0 \pm 0$** |
| | $AB$ | 16 | $0.76 \pm 0.02$ | $36 \pm 6$ |
| 3D | $AB^{cx}$ | 14 | $0.73 \pm 0.02$ | $1.5 \pm 0.4$ |
| | $AB$ | 18 | $0.76 \pm 0.02$ | $36 \pm 6$ |
| 4D | $AB^{cx}$ | 14 | $0.73 \pm 0.02$ | $4.3 \pm 0.5$ |
| | $AB$ | 14 | $0.77 \pm 0.02$ | $38 \pm 6$ |

**Supplementary Table 3.** Monkey F data minimum LOOCV MSE ± sem, min Δ(MSE ± sem) and corresponding latent dimensionality for which it is achieved, for LDS models with different input dimensionalities and contextual constraints. Same conventions as in Supplementary Table 1. Performance in the last column is given relative to the best performing model (TFR 2D *AB*^*cx*^ model, see Supplementary Table 4).

| Input dim | Contextual constraints | Latent dim | min MSE | min ΔMSE ($\times 10^{-3}$) |
| --- | --- | --- | --- | --- |
| 1D | $A^{cx}, B^{cx}$ | 12 | $0.75 \pm 0.02$ | $18 \pm 2$ |
| | $A^{cx}, B$ | 14 | $0.75 \pm 0.02$ | $19 \pm 2$ |
| | $A, B^{cx}$ | 12 | $0.75 \pm 0.02$ | $18 \pm 2$ |
| | $A, B$ | 15 | $0.77 \pm 0.02$ | $39 \pm 4$ |
| 2D | $A^{cx}, B^{cx}$ | 13 | $0.73 \pm 0.02$ | $3.1 \pm 0.3$ |
| | $A^{cx}, B$ | 14 | $0.73 \pm 0.02$ | $3.6 \pm 0.4$ |
| | $A, B^{cx}$ | 13 | $0.73 \pm 0.02$ | $3.0 \pm 0.4$ |
| | $A, B$ | 15 | $0.75 \pm 0.03$ | $25 \pm 4$ |
| 3D | $A^{cx}, B^{cx}$ | 13 | $0.73 \pm 0.02$ | $1.1 \pm 0.5$ |
| | $A^{cx}, B$ | 13 | $0.73 \pm 0.02$ | $1.8 \pm 0.4$ |
| | $A, B^{cx}$ | **13** | **$0.73 \pm 0.02$** | **$0.6 \pm 0.5$** |
| | $A, B$ | 14 | $0.75 \pm 0.03$ | $24 \pm 4$ |
| 4D | $A^{cx}, B^{cx}$ | 13 | $0.73 \pm 0.02$ | $1.0 \pm 0.4$ |
| | $A^{cx}, B$ | 13 | $0.73 \pm 0.02$ | $1.2 \pm 0.4$ |
| | $A, B^{cx}$ | 12 | $0.73 \pm 0.02$ | $0.8 \pm 0.4$ |
| | $A, B$ | 13 | $0.75 \pm 0.03$ | $24 \pm 4$ |

**Supplementary Table 4.** Monkey F data minimum LOOCV MSE ± sem, min Δ(MSE ± sem) and corresponding latent dimensionality for which it is achieved, for TFR models with different input dimensionalities and contextual constraints. Same conventions as in Supplementary Table 1. Performance in the last column is given relative to the best performing model (TFR 2D *AB*^*cx*^ model).

| Input dim | Contextual constraints | Latent dim | min MSE | min ΔMSE ($\times 10^{-3}$) |
| --- | --- | --- | --- | --- |
| 1D | $AB^{cx}$ | 12 | $0.74 \pm 0.02$ | $12 \pm 2$ |
| | $AB$ | 13 | $0.77 \pm 0.02$ | $43 \pm 4$ |
| 2D | $AB^{cx}$ | **12** | **$0.73 \pm 0.02$** | **$0 \pm 0$** |
| | $AB$ | 13 | $0.75 \pm 0.03$ | $23 \pm 4$ |
| 3D | $AB^{cx}$ | 12 | $0.73 \pm 0.02$ | $1.0 \pm 0.3$ |
| | $AB$ | 12 | $0.75 \pm 0.03$ | $22 \pm 4$ |
| 4D | $AB^{cx}$ | 12 | $0.73 \pm 0.02$ | $4.2 \pm 0.5$ |
| | $AB$ | 12 | $0.75 \pm 0.03$ | $24 \pm 4$ |

**Supplementary Table 5.** RNN data minimum LOOCV MSE ± sem, min Δ(MSE ± sem) and corresponding latent dimensionality for which it is achieved, for LDS models with different input dimensionalities and contextual constraints. Same conventions as in Supplementary Table 1. Performance in the last column is given relative to the best performing model (LDS 4D *A, B*^*cx*^ model).

| Input dim | Contextual constraints | Latent dim | min MSE | min ΔMSE ($\times 10^{-3}$) |
| --- | --- | --- | --- | --- |
| 1D | $A^{cx}, B$ | 23 | $0.039 \pm 0.003$ | $27 \pm 2$ |
| | $A, B^{cx}$ | 26 | $0.039 \pm 0.003$ | $27 \pm 2$ |
| 2D | $A^{cx}, B$ | 13 | $0.017 \pm 0.001$ | $5.1 \pm 0.5$ |
| | $A, B^{cx}$ | 26 | $0.017 \pm 0.001$ | $5.0 \pm 0.6$ |
| 3D | $A^{cx}, B$ | 14 | $0.013 \pm 0.001$ | $0.8 \pm 0.3$ |
| | $A, B^{cx}$ | 26 | $0.012 \pm 0.001$ | $0.1 \pm 0.4$ |
| 4D | $A^{cx}, B$ | 15 | $0.013 \pm 0.001$ | $0.6 \pm 0.2$ |
| | $A, B^{cx}$ | **24** | **$0.012 \pm 0.001$** | **$0 \pm 0$** |

